# Dietary manipulation of intestinal microbes prolongs survival in a mouse model of Hirschsprung disease

**DOI:** 10.1101/2025.02.10.637436

**Authors:** Naomi E. Butler Tjaden, Megan J. Liou, Sophie E. Sax, Nejia Lassoued, Meng Lou, Sabine Schneider, Katherine Beigel, Joshua D. Eisenberg, Emma Loeffler, Sierra E. Anderson, Guang Yan, Lev Litichevskiy, Lenka Dohnalová, Yixuan Zhu, Daniela Min Jing Che Jin, Jessica Raab, Emma E. Furth, Zachary Thompson, Ronald C. Rubenstein, Nicolas Pilon, Christoph A. Thaiss, Robert O. Heuckeroth

## Abstract

Enterocolitis is a common and potentially deadly manifestation of Hirschsprung disease (HSCR) but disease mechanisms remain poorly defined. Unexpectedly, we discovered that diet can dramatically affect the lifespan of a HSCR mouse model (*Piebald lethal*, *sl/sl*) where affected animals die from HAEC complications. In the *sl/sl* model, diet alters gut microbes and metabolites, leading to changes in colon epithelial gene expression and epithelial oxygen levels known to influence colitis severity. Our findings demonstrate unrecognized similarity between HAEC and other types of colitis and suggest dietary manipulation could be a valuable therapeutic strategy for people with HSCR.

**Abstract:** Hirschsprung disease (HSCR) is a birth defect where enteric nervous system (ENS) is absent from distal bowel. Bowel lacking ENS fails to relax, causing partial obstruction. Affected children often have “Hirschsprung disease associated enterocolitis” (HAEC), which predisposes to sepsis. We discovered survival of *Piebald lethal* (*sl/sl*) mice, a well-established HSCR model with HAEC, is markedly altered by two distinct standard chow diets. A “Protective” diet increased fecal butyrate/isobutyrate and enhanced production of gut epithelial antimicrobial peptides in proximal colon. In contrast, “Detrimental” diet-fed *sl/sl* had abnormal appearing distal colon epithelium mitochondria, reduced epithelial mRNA involved in oxidative phosphorylation, and elevated epithelial oxygen that fostered growth of inflammation-associated *Enterobacteriaceae*. Accordingly, selective depletion of *Enterobacteriaceae* with sodium tungstate prolonged *sl/sl* survival. Our results provide the first strong evidence that diet modifies survival in a HSCR mouse model, without altering length of distal colon lacking ENS.

**Highlights:**

- Two different standard mouse diets alter survival in the *Piebald lethal* (*sl/sl*) mouse model of Hirschsprung disease, without impacting extent of distal colon aganglionosis (the region lacking ENS).
- *Piebald lethal* mice fed the “Detrimental” diet had many changes in colon epithelial transcriptome including decreased mRNA for antimicrobial peptides and genes involved in oxidative phosphorylation. Detrimental diet fed *sl/sl* also had aberrant-appearing mitochondria in distal colon epithelium, with elevated epithelial oxygen that drives lethal *Enterobacteriaceae* overgrowth via aerobic respiration.
- Elimination of *Enterobacteriaceae* with antibiotics or sodium tungstate improves survival of *Piebald lethal* fed the “Detrimental diet”.

**Graphical abstract:** 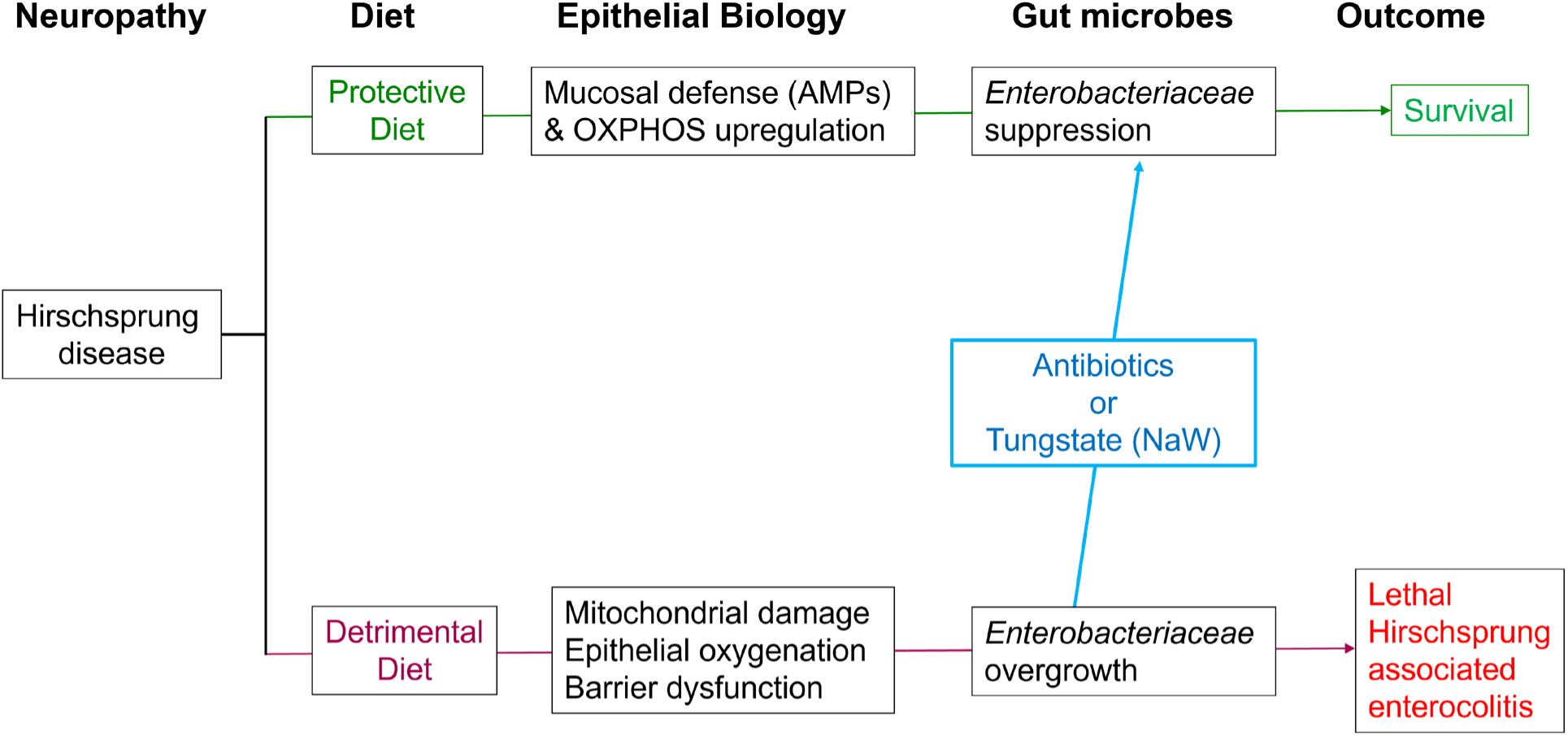

## Introduction

Hirschsprung disease (**HSCR**) is defined by absence of the enteric nervous system (**ENS**) from distal bowel at birth^1–6^. The ENS is an extensive network of neurons and glia that controls most bowel functions including motility, blood flow, immune function, and epithelial biology^7, 8^. When the ENS is missing, the bowel does not work well. HSCR symptoms include intractable constipation, abdominal distension, malnutrition, growth failure, bilious vomiting, bowel perforation and a predisposition to bowel inflammation called Hirschsprung disease associated enterocolitis (**HAEC**)^2, 9, 10^. When HAEC occurs, sepsis may ensue, causing death in childhood. Current therapy for HSCR is “pull-through” surgery to remove the aganglionic bowel lacking ENS and re-attach “normal” bowel to near the anal verge^5, 11^. This surgery is lifesaving, but not always curative. HAEC symptoms, including fever, lethargy, abdominal distension, and explosive (sometimes bloody) diarrhea, remain common after pull-through surgery^2, 10, 12, 13^. Current therapy for HAEC is limited to rectal irrigation, antibiotics, diverting ostomy, and “re-do” pull-through surgery^14–16^. HAEC recurrence is common^17^ and HAEC remains the leading cause of HSCR-associated morbidity and mortality^2, 18^ before and after surgery.

HAEC mechanisms remain incompletely defined but are thought to include epithelial barrier dysfunction, abnormal bowel mucus, and defects in mucosal immune defense that facilitate bacterial translocation across the epithelial barrier and predispose to bowel inflammation^2, 9, 10, 12, 19–27^. Remarkably, although distal bowel “aganglionosis” (absence of the ENS) is always present at birth, some children develop HAEC as neonates and may be critically ill, while others are nearly asymptomatic for months or years after birth^1, 2, 28, 29^. Furthermore, while some children have repeated HAEC episodes, others never develop HAEC^21, 30^ and appear virtually “cured” after pull-through surgery. These differences in HSCR symptom onset, character, and severity occur even in people with similar extents of distal bowel aganglionosis, suggesting additional genetic or environmental factors contribute to HAEC risk. Although we now know a lot about HSCR genetics^2, 31–34^, the only potent genetic modifier known to increase HAEC occurrence is Down syndrome^12, 35–38^, a condition that may cause immune dysregulation^39^. These observations suggest non-genetic factors like diet and gut microbes could critically influence HAEC risk in HSCR but this topic is barely studied.

In other diseases, dietary metabolites, gut microbes, and microbe-derived metabolites are well known to impact intestinal epithelium and mucosal immune system^40–44^, altering risk of inflammation and microbial translocation across the intestinal barrier. For example, specific diets are used clinically to reduce inflammation in Crohn disease^44^, but diet has not been studied in HSCR despite parental desire and attempts to use diet to treat HSCR symptoms^45–47^. Recently, a remarkable prospective human HSCR study revealed that exclusive breastfeeding reduces post-operative HAEC by 40%^21^, demonstrating therapeutic potential for dietary interventions. However, human milk contains many biologically active components that are not present in other foods^48–52^, and breastmilk is not available for all affected children.

To study HAEC, we use the *Piebald lethal* (*sl/sl*) HSCR mouse model^53^ that has an inactivating mutation in *endothelin receptor B* (*Ednrb*)^54^. *EDNRB* mutations also cause human HSCR^55^. EDNRB is a G protein coupled receptor that is expressed by enteric neural crest derived precursors (ENCDC), activated by endothelin-3 (EDN3), and required for ENCDC to colonize distal bowel during fetal development^54–57^. In our colonies, all *sl/sl* mice have distal colon aganglionosis as others reported^54,58^. Like human HSCR, the distal aganglionic colon in *sl/sl* mice fails to relax and lacks propagating contractions causing functional obstruction. Proximal colon migrating motor complex is also absent in *sl/sl* mice^59^. As a result, proximal ENS-containing colon becomes dilated (megacolon) and all *sl/sl* develop megacolon and bowel inflammation prior to death. Serendipitously, we discovered *sl/sl* live 5.7-fold longer at The Children’s Hospital of Philadelphia Research Institute (CHOP) than at University of Quebec at Montreal (UQAM)^60^. Here we show that diet is the primary driver for this survival difference between mouse colonies. The diets we compare are both “standard rodent chow” prepared from natural foods but differ significantly in dietary metabolites and food components. Stool microbes, fecal metabolites, and bowel epithelial transcriptome in *sl/sl* also differ depending on diet in ways that plausibly explain diet-dependent differences in *sl/sl* survival. These observations suggest dietary manipulation or specific nutrient supplementation might also prevent human HAEC.

## Results

### Diet influences survival of *sl/sl* mice

Our *sl/sl* mice were obtained from The Jackson Laboratory where this strain originated^53^ and were maintained at CHOP for several years by breeding *s/sl* x *s/sl* mice before transferring a few breeding pairs to UQAM (with permission from The Jackson Laboratory). Surprisingly, median *sl/sl* survival at CHOP was 149 days, whereas within 2 generations at UQAM, median *sl/sl* survival was only 26 days (**Figure 1A**, p<0.0001) as we previously reported^60^. All *sl/sl* mice at CHOP and UQAM developed dilated stool filled colon (**Figure 1B**), with proximal bowel dilation that becomes more severe with age.

**Figure 1:**
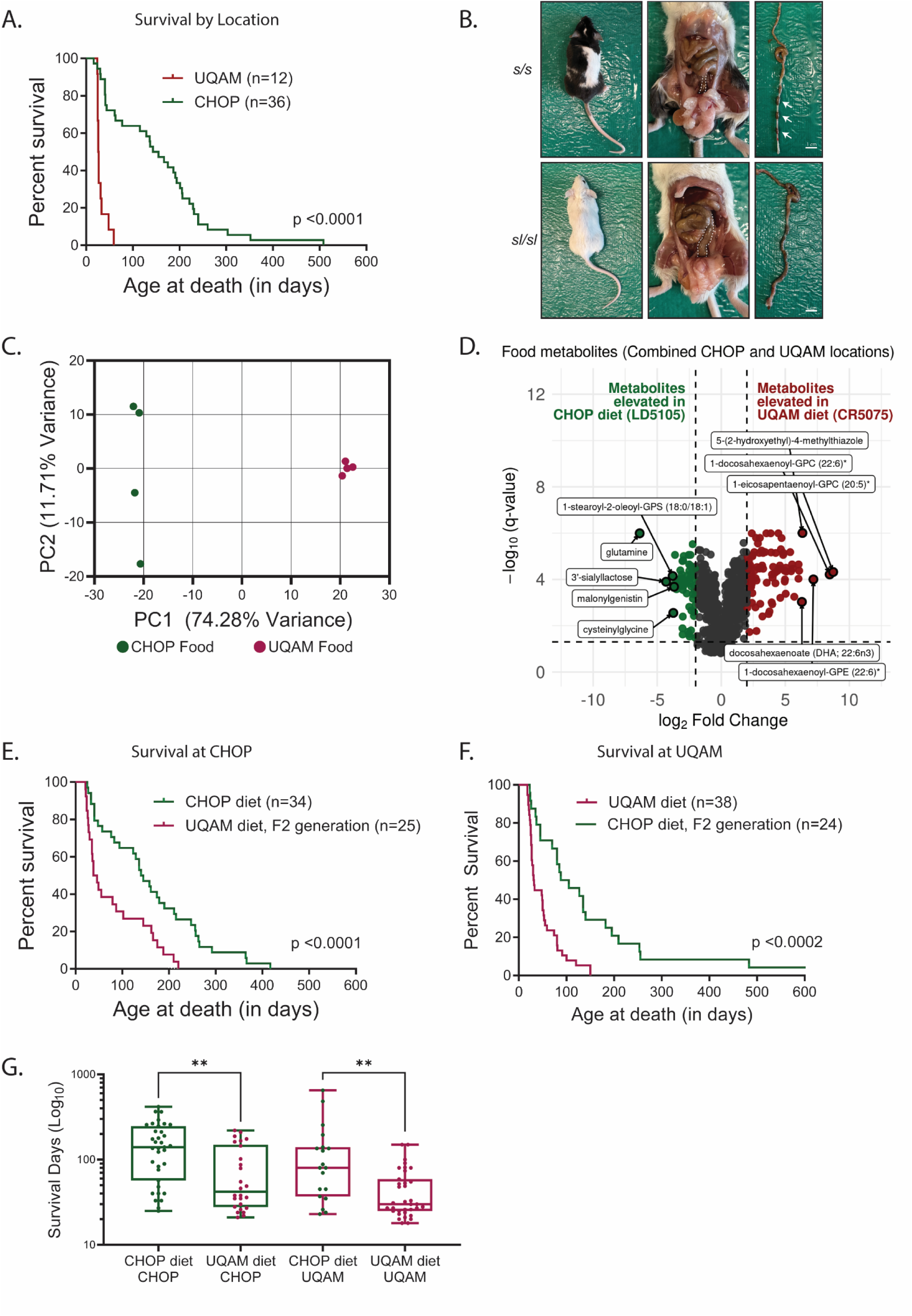
Survival of the Piebald Lethal sl/sl Hirschsprung model is diet-dependent. A. Kaplan-Meyer survival curves show that *sl/sl* housed at CHOP lived much longer than *sl/sl* housed at UQAM. CHOP is The Children’s Hospital of Philadelphia Research Institute; UQAM is University of Quebec at Montreal. For each curve, mice were housed at this location for at least two generations. Log-rank (Mantel-Cox) test p<0.0001. Median age of survival at CHOP was 149 days (n=36); median age at UQAM was 26.5 days (n=12). *Reprinted from Gastroenterology, 159(5), Soret R, Schneider S, Bernas G, Christophers B, Souchkova O, Charrier B, Righini-Grunder F, Aspirot A, Landry M, Kembel SW, Faure C, Heuckeroth RO, Pilon N, Glial Cell-Derived Neurotrophic Factor Induces Enteric Neurogenesis and Improves Colon Structure and Function in Mouse Models of Hirschsprung Disease, pages 1824-1838.e17, Copyright (2000), with permission from Elsevier.* B. At P100 *s/s* (top row) can be distinguished from *sl/sl* (bottom row) by coat color since *Ednrb* is needed for melanocyte and ENS development. P100 colon in *sl/sl* mice was often dilated, filled with stool and lacked discrete stool pellets that are easily distinguished in *s/s* colons (white arrows). Middle panel shows gross anatomy with colon highlighted by dashed lines. Right panel shows colon with attached terminal ileum. Images are from mice housed at CHOP but phenotypes were similar at CHOP and UQAM. Scale bars = 1cm. C. Principal coordinate analysis of food metabolites for the “Protective” (LabDiet 5015, green = standard mouse diet at CHOP) and “Detrimental” (Charles River 5075, magenta = standard mouse diet at UQAM) diets based on untargeted mass spectrometry, which defined relative abundance for 580 metabolites. 4 food pellets from each diet were analyzed. Dots represent individual food pellets. A. Volcano plot showing relative abundance of 581 metabolites versus statistical significance of difference in metabolite abundance for “Protective” and “Detrimental” diets. Green dots = More abundant in “Protective” diet. Magenta dots = More abundant in “Detrimental” diet. Vertical dotted lines indicated Log_2_ fold difference in metabolite abundance = 2. Dots above the horizontal dashed line have adjusted p-value (q-value) < 0.05 (Welch two-sample t-test). Left of center indicates metabolites more abundant in CHOP diet LabDiet5015; Right of center represents metabolites more abundant in UQAM diet CharlesRiver5075. Labeled metabolites elevated on the “Protective” diet: glutamine, 3’-sialyllactose, 1-stearoyl-2-oleoyl-GPS (18:0/18:1), cysteinylglycine, and malonylgenisten. Labeled metabolites elevated on the “Detrimental” diet: 5-(2-hydroxyethyl)-4-methylthiazole, 1-docosahexaenoyl-GPC (22:6), 1-eicosapentaenoyl-GPC (20:5), 1-docosahexaenoyl-GPE (22:6), and docosahexanoate (DHA; 22: 6n3). D. Kaplan-Meyer survival curve at CHOP for *sl/sl* fed the CHOP “Protective” diet (LabDiet5015 (Green)) or the UQAM “Detrimental” diet (CharlesRiver5075 (Magenta)). Mice at CHOP been maintained on the CHOP diet for at least 2 generations and on the UQAM diet for exactly 2 generations (i.e., grandparents were weaned to the designated diet, F2). Median age for *sl/sl* death when fed LabDiet5015 was 143 days (n=34). Median age at death for *sl/sl* fed CharlesRiver5075 was 42 days (n=25). Log-rank (Mantel-Cox) test p<0.0041. E. Kaplan-Meyer survival curve at UQAM for *sl/sl* fed the CHOP “Protective” diet (LabDiet5015 (Green)) or the UQAM “Detrimental” diet (CharlesRiver5075 (Magenta)). Mice at UQAM had been maintained on the UQAM diet for at least 2 generations and on the CHOP diet for exactly 2 generations (i.e., grandparents were weaned to the designated diet, F2). Median age for *sl/sl* death when fed LabDiet5015 was 80 days (n=20). Median age for *sl/sl* death when fed CharlesRiver5075 was 30 days (n=38). Log-rank (Mantel-Cox) test p<0.0002. F. Box plots of age at death for *sl/sl* based on location and diet. Mice had been maintained on each diet for at least 2 generations. Green outlines = CHOP location. Magenta outlines = UQAM location. Green circle = “Protective” diet (LabDiet5015). Magenta circle = “Detrimental” diet (CharlesRiver5075). Each dot represents one mouse. Difference in survival at CHOP when fed different diets: p=0.0013 (Mann-Whitney U-test). Difference in survival at UQAM when fed different diets: p=0.0024 (Mann-Whitney U-test).

We hypothesized that differences in *sl/sl* survival were mediated by non-genetic factors in our mouse colonies. In particular, we noted substantial differences in composition of diets routinely used in our vivaria (UQAM diet = Charles River Rodent Diet 5075 or CR-5075; CHOP diet = LabDiet Mouse Diet 5015 or LD-5015). Both diets are “standard rodent chow” but contain differences in the abundance of vitamins and minerals, fat source (pork fat and soybean oil versus poultry fat), protein source (casein versus fish meal), and other ingredients (brewer’s yeast and wheat germ versus wheat middlings) (**Table 1**). In addition, the LD-5015 diet used at CHOP is higher in fat (25.3%) and lower in fiber (2.2%) compared to the CR-5075 diet used at UQAM (12.1% fat, 4.1% fiber). To more broadly define how dietary metabolites differ, we pursued untargeted metabolomic analyses by ultra-high performance liquid chromatography-tandem mass spectrometry (UPLC-MS/MS). This approach showed 581 of 935 quantified molecules differ significantly in abundance between the diets (**Figure 1C, D** q-value < 0.012). For example, the LD-5015 diet used at CHOP has 83-fold more glutamine than the CR-5075 diet used at UQAM, but lower levels of glycerophosphocholines (GPCs) and glycerophosphoethanolamines (GPEs). **Supplemental Tables 1 and 2** list the most differentially abundant dietary metabolites by chow. All dietary metabolite abundance data are in **Supplemental Table 3**. These observations support the hypothesis that differences in the abundance of one or more dietary metabolites could impact survival of the inbred *sl/sl* HSCR mouse model.

**Table 1:** Table lists individual food components that differ by at least 1.5-fold based on food package labeling.

|  | CHOP Institutional Diet | UQAM Institutional Diet |  |
| --- | --- | --- | --- |
|  | Mouse Diet 5015 (LabDiet) | Charles River Rodent Diet 5075 (Cargill Animal Nutrition) | CHOP/UQAM ratio |
|  | “Protective diet” | “Detrimental diet” |  |
| Percent Fat Calories | 25.3% | 12.1% | 2.09 |
| Crude Fiber | 2.2% | 4.1% | 0.54 |
| Vitamin K | 3.0 mg/kg (L) | 8.8 mg/kg | 0.34 |
| Vitamin A | 18 IU/gram (L) | 51 IU/gram | 0.35 |
| Vitamin E | 66 IU/kg (L) | 110 IU/kg | 0.60 |
| Biotin | 0.3mg/kg (L) | 0.45 mg/kg | 0.67 |
| Vitamin D3 | 3.3 IU/gram (H) | 2.2 IU/gram | 1.50 |
| Vitamin B12 | 51 mcg/kg (H) | 26 mcg/kg | 1.96 |
| Copper | 20 mg/kg (H) | 12 mg/kg | 1.67 |
| Cobalt | 0.63 mg/kg (H) | 0.3 mg/kg | 2.10 |
| Iodine | 1.5 mg/kg | 0.67 mg/kg | 2.24 |
|  | Wheat germ | Wheat middlings |  |
|  | Brewer's yeast | Fish meal |  |
|  | Porcine animal fat | Poultry Fat |  |
|  | Soybean oil |  |  |
|  | Casein |  |  |

To test the hypothesis that diet differences could underlie the 5.7-fold difference in median survival between institutions, we fed each diet to new cohorts of *sl/sl* mice at each institution and tracked survival for 2 generations. At CHOP, feeding the UQAM diet CR-5075 reduced median survival of *sl/sl* mice to 42 days for the “second generation”, whereas the *sl/sl* cohort housed in the same room but fed the usual CHOP diet LD-5015 had a median survival of 143 days (**Figure 1E**, 3.4-fold difference, p<0.0001). Conversely, *sl/sl* mice at UQAM fed the CHOP diet LD-5015 for 2 generations had a median survival of 80 days, while *sl/sl* mice simultaneously housed in the same room but fed the usual UQAM diet CR-5075 had median survival of only 30 days (**Figure 1F**, 2.7-fold difference, p<0.005). These data show that diet potently impacts survival of *sl/sl* mice at both CHOP and UQAM (**Figure 1G**). To simplify nomenclature, we now begin to refer to CHOP diet LD-5015 as the “Protective” diet and UQAM diet CR-5075 as the “Detrimental” diet to reflect effects on *sl/sl* survival.

Because we wished to identify factors that predispose to or protect from HAEC instead of the consequences of HAEC, all subsequent studies used animal at P20±1 day, before any *sl/sl* mice died of HAEC complications. For all subsequent studies, *sl/sl* mice were fed the designated diet for at least two generations (i.e., grandparents or earlier generations had been weaned to the designated diet).

### Diet does not alter extent of aganglionosis, weight gain, or colon epithelial anatomy in P20 *sl/sl* mice

All *sl/sl* mice at CHOP had distal colon aganglionosis with similar lengths of aganglionic and “transition zone” (hypoganglionic) colon independent of diet measured at P20±1 day (**Figures 2A, B**). In addition, *sl/sl* mice at both institutions develop bowel inflammation as others reported^61–63^. Because diet-driven differences in survival were greatest at CHOP, we analyzed a cohort of CHOP housed P20 *sl/sl* mice fed different diets for at least 2 generations. Cross-sectional images from proximal and distal colon appeared similar in *sl/sl* mice fed each diet after staining with hematoxylin and eosin (H&E) and periodic acid Schiff (PAS) reagent (**Figure 2C**). There were no significant diet-dependent differences in colon length or mouse body weight (**Figure 2D, 2E**), crypt length (**Figure 2F, G**) or goblet cells per crypt (**Figure 2H, I**). Enterocolitis scores^64^ in proximal or distal colon at P20 were also similar for *sl/sl* mice fed each diet at CHOP (**Figure 2J, K)**. These data suggest, at least at P20, that *sl/sl* mice lack striking diet-dependent differences in nutrition (i.e., growth), ENS anatomy, or colon anatomy that could explain why 50% of *sl/sl* mice fed the Detrimental diet die between P20 and P38 at CHOP (**Figure 1E**), whereas only 11.4% of *sl/sl* mice fed the Protective diet die during this interval in the same vivarium room.

**Figure 2.**
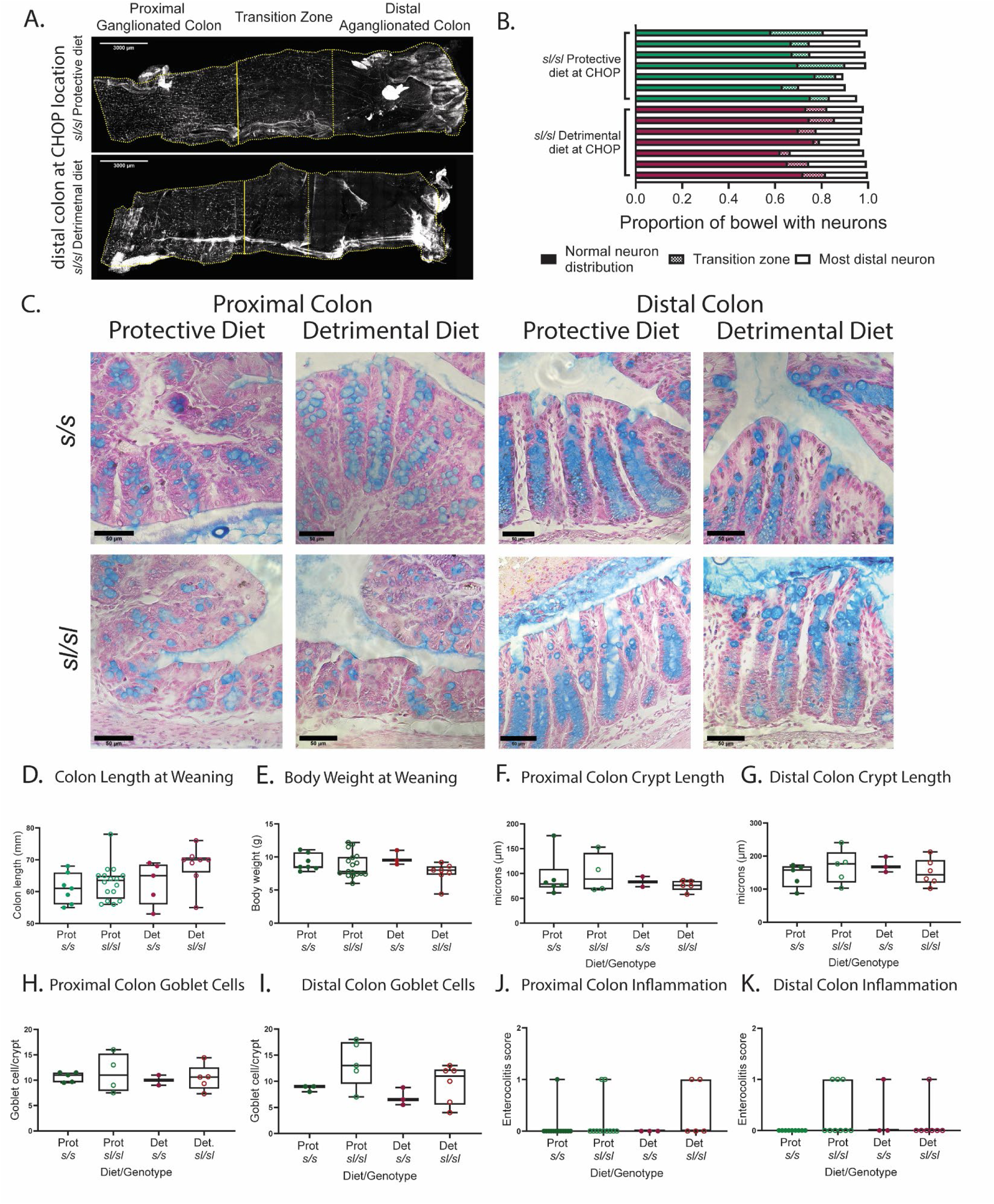
Alternate diets did not impact sl/sl colon length, weight or colon epithelial anatomy at P20 A. Enteric neurons visualized by whole mount ANNA1 (HuC/D) immunohistochemistry in P20 colon from *sl/sl* maintained at CHOP. Distal half of colons are shown. Top image: Protective diet (LD 5015) fed mouse. Bottom image: Detrimental diet (CR 5075) fed mouse. Detrimental diet-fed mice had grandparents weaned to the Detrimental diet and parents were fed the same diet. Protective diet-fed mice had been maintained on the Protective diet for many generations at CHOP. Enteric neurons identified with ANNA1 antibody had a similar distribution in *sl/sl* fed Protective or Detrimental diets with distal colon aganglionosis, a transition zone, and denser ENS in proximal colon. Small white dots are enteric neuron cell bodies. Vertical yellow lines show proximal (thicker line) and distal (thinner line) margins of the measured transition zones. Scale bars = 3 mm. B. Bar graphs show a similar extent of distal colon aganglionosis and similar length transition zone in P20±1 *sl/sl* mice maintained at CHOP on either the Protective or Detrimental diets. Individual Detrimental diet-fed *sl/sl* represented by each bar were from lineages with grandparents weaned to the designated diet. Protective diet-fed *sl/sl* were derived from mice maintained on the Protective diet for many generations. N = 7 for Protective diet and 7 for Detrimental diet. There was no statistical difference in the length of proximal colon with normal appearing ENS (p = 0.62, Mann-Whitney), length of the hypoganglionic transition zone (p = 0.80, Mann-Whitney), or location of most distal myenteric neuron (p = 0.38, Mann-Whitney) for *sl/sl* fed Protective versus Detrimental diets. C. Representative Alcian blue-stained images of proximal and distal colon in *s/s* and *sl/sl* mice show a similar distribution of goblet cells and PAS-stained mucins for mice fed each diet. Scale bar = 50 μm. D. Colon length at P20 did not differ based on diet (p=0.09 ANOVA). Prot = Protective diet fed. Det = Detrimental diet fed. E. Mouse weights at weaning did not differ based on diet at P20 (p=0.18 ANOVA) F-I. There were no diet-dependent differences in crypt length in proximal colon (p=0.73 ANOVA) (F) or distal colon (p=0.64 ANOVA) (G) and no diet-dependent differences in goblet cells per crypt in proximal (p=0.73 ANOVA) (H) or distal colon (p=0.092 ANOVA) (I) at P20. J, K. Proximal colon (p=0.35 ANOVA) (J) and distal colon (p=0.29 ANOVA) (K) enterocolitis scores at P20 (adapted from Cheng 2010^1^) did not differ based on diet (Protective versus Detrimental) or genotype (*s/s* versus *sl/sl*) at CHOP.

### The impact of diet on P20 *sl/sl* mouse stool metabolites

Given the marked differences in metabolite composition between the Protective and Detrimental diets (**Figures 1 C, D, Tables 1, Supplemental Table 1, 2, 3**), we next pursued untargeted metabolomics on stool from P20 *sl/sl* mice fed the Protective or Detrimental diets for at least 2 generations (**Figure 3**). Samples were collected at CHOP and at UQAM from mice fed each diet. When all stool samples (n = 40) were analyzed as a single dataset, only 10 fecal metabolites differed significantly (>4-fold different with q-value < 0.05) in abundance based on diet (**Figures 3A, B, C** and **Table 2**), but effects of diet on stool metabolites were stronger for *sl/sl* in the UQAM vivarium than in the CHOP vivarium (**Figures 3A, D, E** and **Table 2, Supplemental Tables 4, 5, 6**). These observations suggest that diet-dependent differences in fecal metabolites, which differ between vivaria, do not fully explain the consistent effects of each diet on *sl/sl* survival, independent of housing location (**Figures 1E, F**).

**Figure 3:**
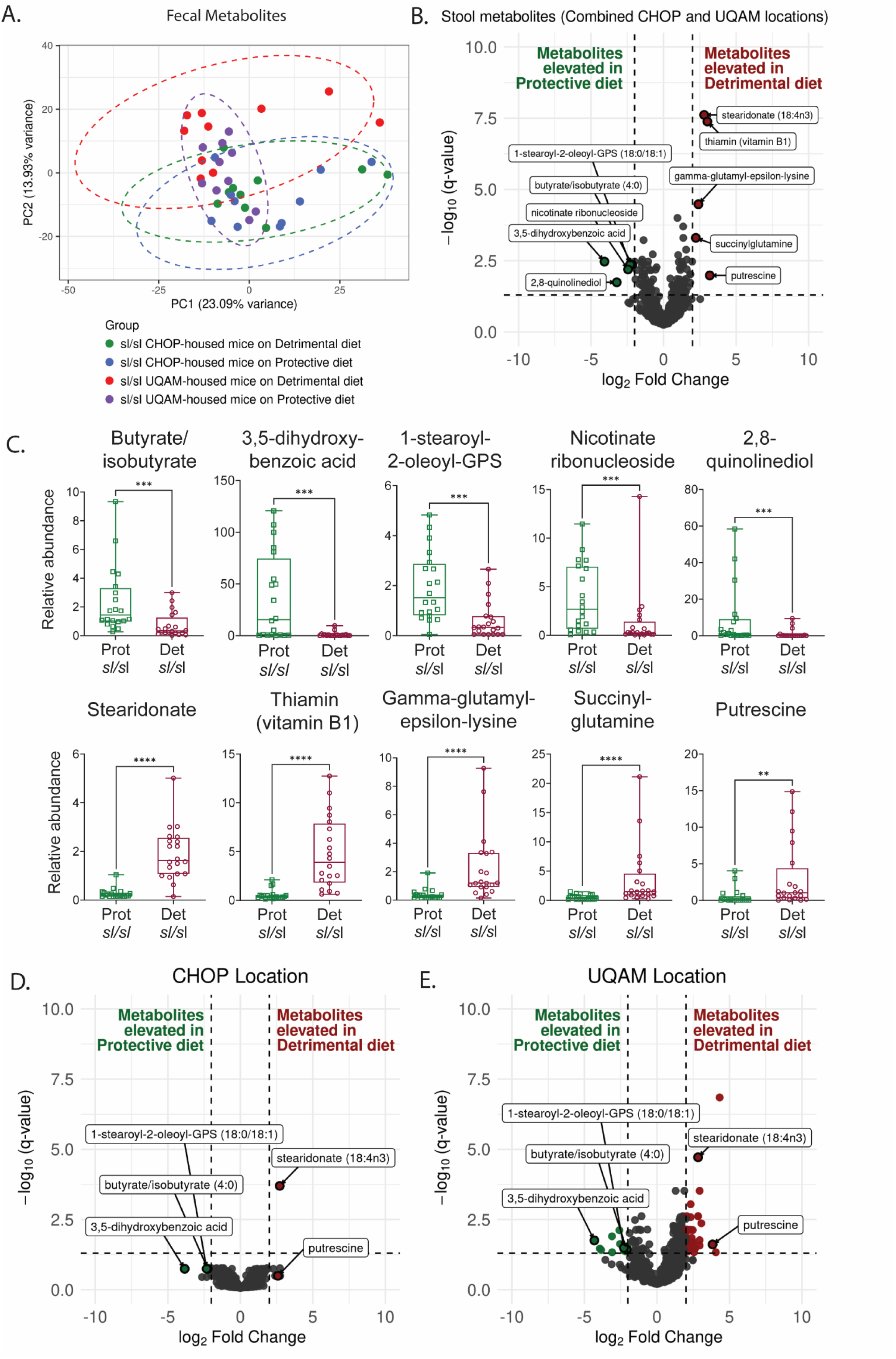
Untargeted mass spectrometry to quantify metabolites in P20 stool showed elevated butyrate/isobutyrate in sl/sl fed the Protective diet compared to sl/sl fed the Detrimental diet. B. Principal coordinate analysis of fecal metabolites from *sl/sl* mice housed at either CHOP or UQAM and fed either “Protective (LabDiet5015) or “Detrimental” (CharlesRiver5075) diets for at least 2 generations. N = 5 males and 5 females fed each diet at each location. Each group 95% confidence interval encircled by corresponding color. Dots represent fecal metabolites from individual mice. C. Volcano plot shows relative abundance of *sl/sl* fecal metabolites versus statistical significance of differences in metabolite abundance for mice fed different diets at both locations (n=40, combined CHOP and UQAM data). Dots above the horizontal dashed line indicates q value <0.05. Vertical dashed lines indicate log2 (Fold-difference) = 2. Labeled metabolites elevated on the “Protective” diet: 1-stearoyl-2-oleoyl-GPS (18:0/18:1), Butyrate/isobutyrate, nicotinate ribonucleoside, 3,5-dihydroxybenzoic acid and 2,8-quinolinediol. Labeled metabolites elevated on the “Detrimental” diet: stearidonate (18:4n3), thiamin (vitamin B1), gamma-glutamyl-epsilon-lysine, succinylglutamine and putrescine. D. Relative abundance of specified fecal metabolites in stool of *sl/sl* mice fed the “Protective” or “Detrimental” diets. Each circle represents metabolite abundance in stool from one mouse (n=40 *sl/sl* mice housed at both locations). Det = At least 2 generations fed the Detrimental diet. Prot = At least 2 generations fed the Protective diet fed. Boxes show interquartile range, error bars show min to max points. Lines within each box indicate median abundance. Metabolites selected are the same as metabolites labeled in the volcano plot (B): P-values (two-tailed Mann-Whitney): Butyrate/isobutyrate ***=0.0003, 3,5-dihydroxybenzoic acid ***=0.0003, 1-stearoyl-2-oleoyl-GPS (18:0/18:1) ***=0.0001, Nicotinate ribonucleoside ***=0.0007, 2,8-quinolinediol ***=0.0005, stearidonate (18:4n3) ****=<0.0001, thiamin (vitamin B1) ****=<0.0001, gamma-glutamyl-epsilon-lysine ****=<0.0001, succinylglutamine **** = <0.0001,and putrescine **=0.0011. D-E. (D) Volcano plot shows relative abundance of *sl/sl* fecal metabolites versus statistical significance of differences in metabolite abundance for mice fed different diets while housed at CHOP. Dots above the horizontal dashed line indicates q value < 0.05 (ANCOVA, FDR). Vertical dashed lines indicate log2 (Fold-difference) = 2. N=20 *sl/sl* mice housed at CHOP (with 10 *sl/sl* fed each diet). (E) Similar to the volcano plot (D) except using data for N=20 *sl/sl* mice housed at UQAM (with 10 *sl/sl* fed each diet). Each dot represents one metabolite. Labeled metabolites: Butyrate/isobutyrate, 3,5-dihydroxybenzoic acid, 1-stearoyl-2-oleoyl-GPS (18:0/18:1), stearidonate (18:4n3), and putrescine.

**Table 2:** Fecal metabolites in *sl/sl* mice that statistically differ more than 4-fold (q < 0.05, ANCOVA) based on diet.

| Fecal metabolites >4-fold different in abundance depending on *sl/sl* diet at both locations |  |
| --- | --- |
| More abundant in stool of *sl/sl* fed the Protective diet | More abundant in stool of *sl/sl* fed the Detrimental diet |
| Butyrate/Isobutyrate (4:0), 4.47-fold higher | Stearidonate (18:4n3)*, 6.95-fold higher |
| 3,5-dihydroxybenzoic acid*, 16.84-fold higher | Thiamin (vitamin B1), 8.10fold higher |
| 1-stearoyl-2-oleoyl-GPS (18:0/18:1), 4.94-fold higher | gamma-glutamyl-epsilon-lysine, 5.29-fold higher |
| nicotinate ribonucleoside, 5.46-fold higher | Succinylglutamine, 4.67-fold higher |
| 2,8-quinolinediol, 9.36-fold higher | Putrescine, 9.10-fold higher |
| *The metabolites 1-stearoyl-2-oleoyl-GPS (18:0/18:1) (13.9-fold), nicotinate ribonucleoside (2.44-fold)stearidonate(15.29-fold), thiamin (4.37 fold), and putrescine (1.6-fold) are more abundant in Protective or Detrimental diets (i.e., in food pellets) respectively (fold change in parenthesis) and could be diet-derived, whereas other metabolites listed here do not differ significantly or were not detected in mouse food. |  |

### Diet exerts strong effects on the proximal colon epithelial transcriptome of P20 *sl/sl* mice

Given that fecal metabolites were poor indicators of survival in contrast to the strongly distinct dietary metabolite profiles, we focused on possible impacts of the two different diets on intestinal epithelial cells. Colonic epithelial biology is a critical component of the barrier that prevents inflammation and sepsis in HSCR. To test the hypothesis that Protective and Detrimental diets alter *sl/sl* colon epithelium gene expression, we performed bulk RNA sequencing from proximal and distal colon epithelium of P20 *sl/sl* mice housed at CHOP (**Figure 4A**). All mice had been fed the designated diet for at least 2 generations. Principal coordinate analysis shows major differences in mRNA abundance patterns between epithelial cells in proximal versus distal colon along the PC1 axis **(Figure 4B**). Significant diet-dependent differences in proximal colon epithelial mRNA abundance patterns were also evident along the PC1 axis when principal coordinate analyses were pursued for proximal colon alone, excluding distal colon data (**Figure 4C**).

**Figure 4:**
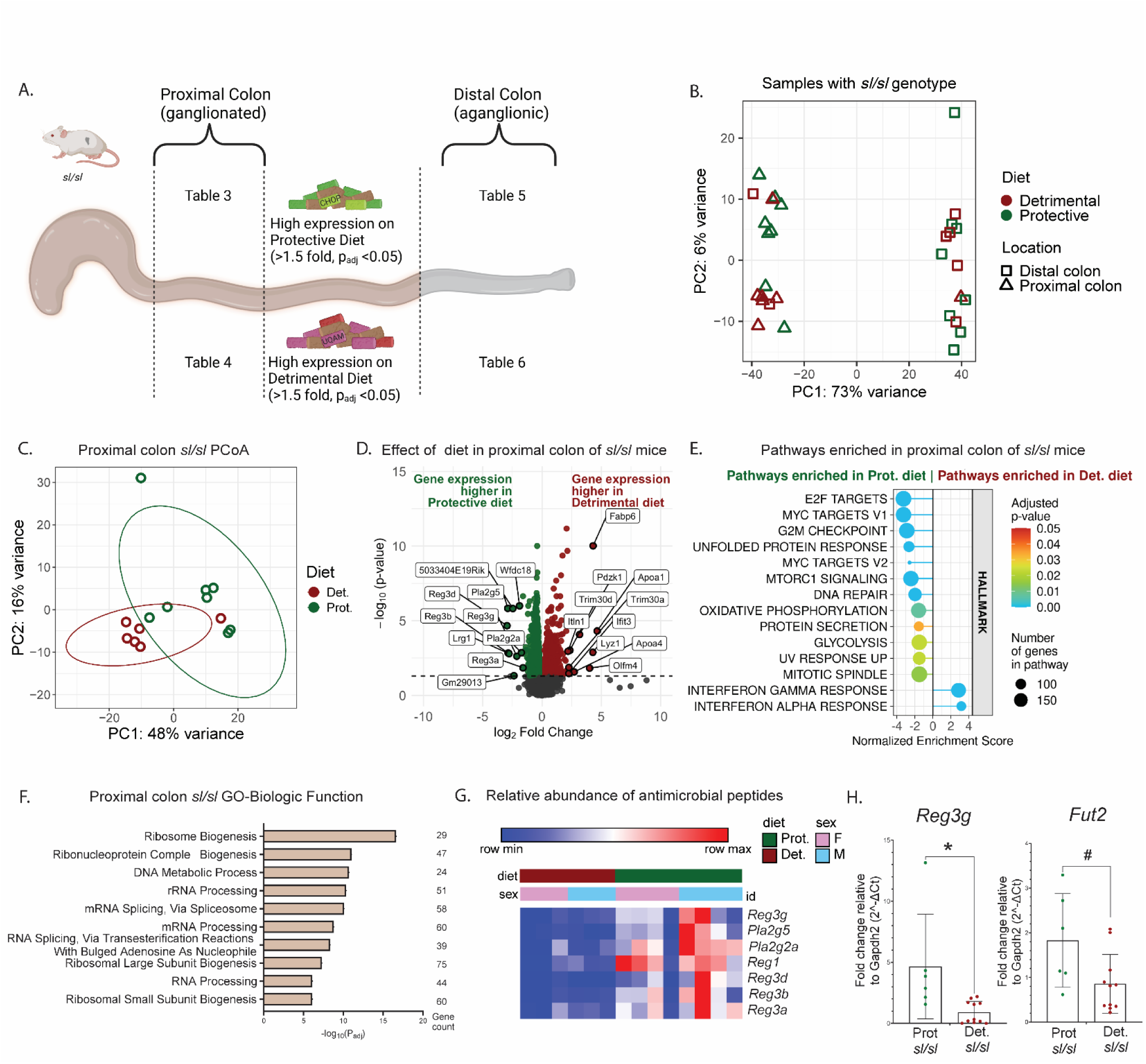
Colon epithelial transcriptome varies depending on diet (Protective versus Detrimental) and colon region (Proximal versus Distal). A. Experimental plan. Bulk RNA sequencing of colon epithelial cells from 1.5 cm of ENS-containing proximal colon or 1.5 cm of aganglionic distal colon. Groups: P20 *sl/sl* mice fed either Protective or Detrimental diet for at least 2 generations. N = 32 RNA pools (8 mice/group (4 males, 4 females) x 2 colon regions x 2 diets =32 pools). Top 10 differentially expressed genes in each group are in Tables 3-6 as indicated. B. Principal coordinate analysis shows colon epithelial RNA abundance patterns from all *sl/sl* mice. Each symbol represents one mouse. C. Principal coordinate analysis shows epithelial RNA abundance patterns from only the proximal colon of *sl/sl* mice fed the Protective (Prot) or Detrimental (Det) diets. Each symbol represents one mouse. D. Volcano plot showing log_2_(Fold-change) versus-log_10_(adjusted p-value) for proximal colon mRNA of *sl/sl* fed the Protective (green) or Detrimental (magenta) diets. Dots above the horizontal dashed line have adjusted p value < 0.05 using Wald test and Benjamini and Hochberg correction in the DESeq2 package. E. Fgsea pathway analysis of diet-dependent differentially expressed genes (DEGs) (adjusted p <0.05) from *sl/sl* proximal colon epithelial cells. Negative normalized enrichment scores indicate pathways with mRNA that were more abundant in *sl/sl* proximal colon epithelial cells from mice fed the Protective diet. Positive enrichment scores indicate pathways with mRNA more abundant in proximal colon epithelial cells of *sl/sl* fed the Detrimental diet. F. Enriched gene ontology (GO) analysis for biological functions (BF) of genes differentially expressed in *sl/sl* proximal colon epithelial cells depending on diet. G. Heatmap shows relative abundance of mRNA for antimicrobial peptides differs based on diet in proximal colon epithelium of *sl/sl* mice. All antimicrobial peptide mRNA were more abundant in *sl/sl* fed the Protective diet compared to *sl/sl* fed the Detrimental diet. H. Relative abundance of mRNA based on qRT-PCR for the antimicrobial peptide *Reg3g* and for fucosyltransferase 2 (*Fut2*) in proximal colon epithelium of *sl/sl* fed the Protective (green dots) or Detrimental (magenta dots) diet for two generations. * = 0.031; # = 0.043 Paired Wilcoxon test.

We identified 2134 diet-dependent differences in *sl/sl* proximal colon epithelial cell mRNA abundance (**Figure 4D**, Benjamini-Hochberg (BH) corrected p < 0.05). GSEA pathway analysis showed that many differentially expressed transcripts more abundant in proximal colon epithelium of *sl/sl* mice fed the Protective diet were linked cell cycle or cellular stress response pathways (**Figure 4E**). Gene ontology (GO-BF (enrichr)) (**Figure 4F**) pathway analyses show that transcript differences in ribosome-and spliceosome-associated pathways are prominent features of diet-dependent differences in *sl/sl* proximal colon epithelial gene expression. However, 6 of the 10 most differentially expressed mRNA that were more abundant in proximal colon epithelium of *sl/sl* fed the Protective diet encode antimicrobial peptides (**Table 3**, **Figure 4G**): *Reg3g, Reg3d, Reg3b, Reg3a, Pla2g2a, Pla2g5*. Additionally, the fucosyltransferase *Fut2* is significantly increased in the proximal colon mRNA of Protective diet-fed compared to Detrimental diet-fed *sl/sl*. Fucosyltransferases add fucose to intestinal epithelial glycoproteins. The fucose supports attachment of specific bacteria and fungi to surface glycoproteins. When fucose is present, beneficial bacteria thrive and pathogenic bacteria are inhibited^65^. In contrast, the most differentially expressed transcripts that were more abundant in proximal colon epithelium of *sl/sl* mice fed the Detrimental diet include genes critical in lipid transport and metabolism (*Apoa1*, *Apoa4*, *Fabp6*) as well as mRNA encoding neutrophil proteins lysozyme (*Lyz1*) and olfactomedin 4 (*Olfm4*), and negative regulators of the NLRP3 inflammasome (Trim30a, Trim30d)^66^ (**Table 4**). *Reg3g* and the fucosyltransferase *Fut2* were validated by qRT-PCR (**Figure 4H**).

**Table 3:** Top 10 differentially expressed mRNA more abundant in proximal colon epithelial cells of sl/sl fed the Protective diet compared to sl/sl fed the Detrimental diet.

| Proximal colon most differentially expressed genes, Protective Diet |  |  |  |  |
| --- | --- | --- | --- | --- |
| Gene name | Mean counts | log<sub>2</sub> Fold-difference in mRNA abundance | log<sub>2</sub> Fold-difference std. error | p<sub>adj</sub> |
| *Reg3d* | 69.00 | 2.95 | 0.55 | 2.15E-05 |
| *Pla2g5* | 353.97 | 2.89 | 0.49 | 1.49E-06 |
| *Reg3b* | 117351.90 | 2.86 | 0.69 | 1.46E-03 |
| *Lrg1* | 123.34 | 2.78 | 0.68 | 1.61E-03 |
| *5033404E19Rik* | 1653.80 | 2.46 | 0.41 | 1.53E-06 |
| *Gm29013* | 217.27 | 2.37 | 0.86 | 4.83E-02 |
| *Reg3g* | 31466.48 | 2.13 | 0.54 | 2.38E-03 |
| *Wfdc18* | 278.63 | 1.90 | 0.31 | 1.01E-06 |
| *Pla2g2a* | 206.31 | 1.75 | 0.42 | 1.39E-03 |
| *Reg3a* | 82.64 | 1.61 | 0.49 | 1.42E-02 |

**Table 4:** Top 10 differentially expressed mRNA more abundant in proximal colon epithelial cells of sl/sl fed the Detrimental diet compared to sl/sl fed the Protective diet.

| Proximal colon most differentially expressed genes, Detrimental Diet |  |  |  |  |
| --- | --- | --- | --- | --- |
| Gene name | Mean counts | log<sub>2</sub> Fold-difference in mRNA abundance | log<sub>2</sub> Fold-difference std. error | p<sub>adj</sub> |
| *Apoa1* | 509.73 | 4.64 | 0.91 | 4.87E-05 |
| *Fabp6* | 1743.30 | 4.30 | 0.56 | 9.62E-11 |
| *Lyz1* | 101.16 | 4.28 | 1.02 | 1.30E-03 |
| *Olfm4* | 106.40 | 4.01 | 1.22 | 1.49E-02 |
| *Pdzk1* | 45.90 | 3.17 | 0.64 | 8.37E-05 |
| *Apoa4* | 75.12 | 2.71 | 0.90 | 2.68E-02 |
| *Trim30d* | 141.32 | 2.35 | 0.55 | 9.55E-04 |
| *Trim30a* | 745.56 | 2.26 | 0.68 | 1.34E-02 |
| *Ifit3* | 2015.24 | 2.26 | 0.78 | 3.41E-02 |
| *Itln1* | 369.80 | 2.19 | 0.52 | 1.11E-03 |

In contrast to proximal colon, distal colon mRNA abundance patterns did not clearly separate in principal coordinate analysis by diet (**Figure 5A**). However, diet exerted an effect on the distal colon epithelial transcriptomes of *sl/sl* mice, with 1191 differentially expressed genes identified (BH-corrected p < 0.05) (**Figure 5B**). The 10 mRNA most differentially abundant in distal colon epithelium of *sl/sl* mice based on diet are in **Table 5** (higher in Protective diet-fed) and **Table 6** (higher in Detrimental diet-fed). GSEA pathway analysis showed that many differentially expressed transcripts more abundant in distal colon epithelium of *sl/sl* mice fed the Protective diet were involved in oxidative phosphorylation (OXPHOS) or cell growth (**Figure 5C**). Gene ontology (GO-BF (enrichr)) analysis confirmed enrichment of transcripts associated with OXPHOS (e.g., terms like mitochondrial ATP synthesis coupled electron transport, cellular respiration, oxidative phosphorylation) in distal colon epithelium of *sl/sl* mice fed the Protective diet (**Figure 5D**) compared to Detrimental diet-fed *sl/sl* mice. Relative abundance of individual OXPHOS-related transcripts in *sl/sl* distal colon epithelium are shown in a heatmap (**Figure 5E**).

**Figure 5:**
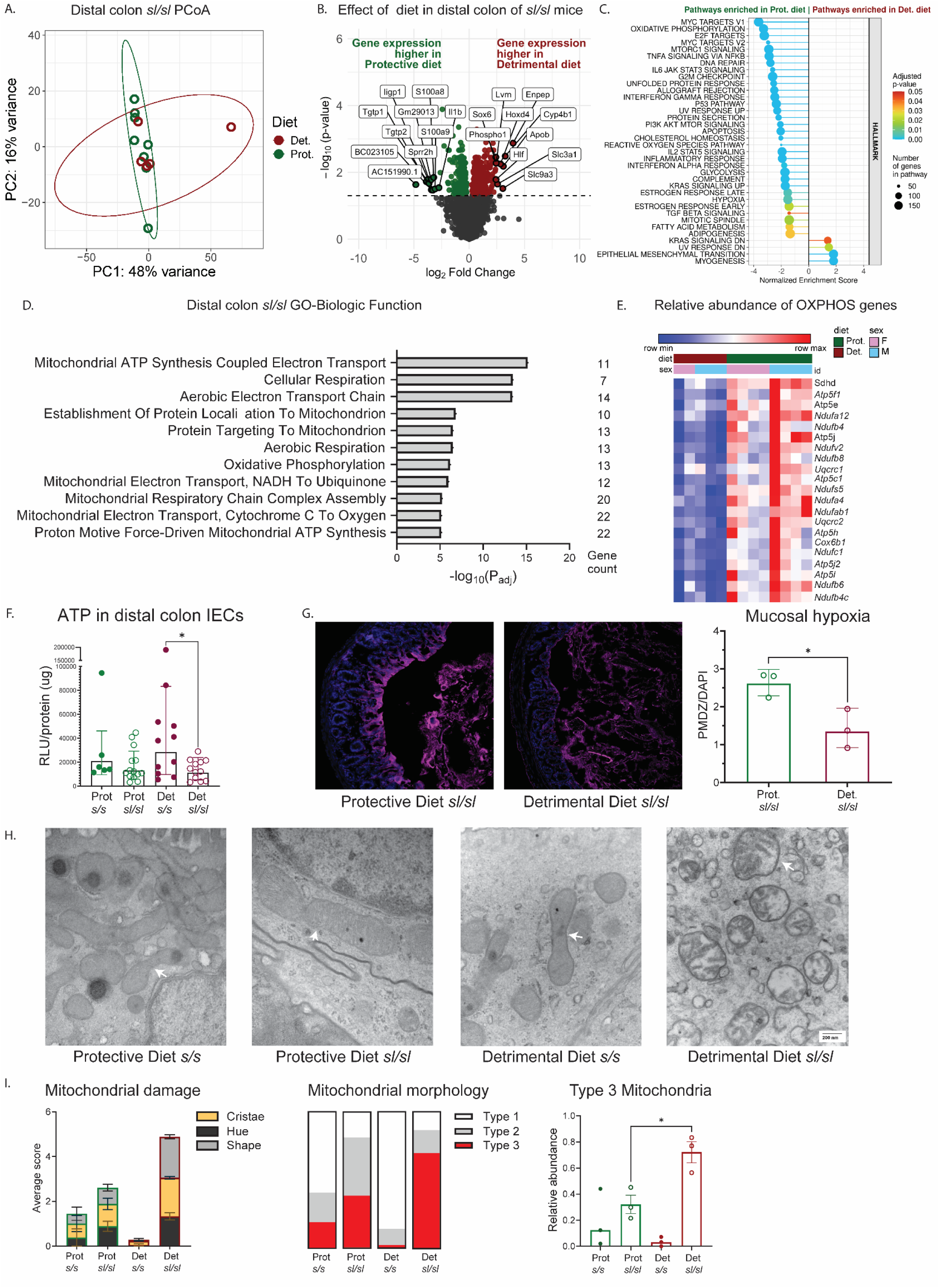
Distal colon epithelial transcriptome for sl/sl housed at CHOP revealed diet-dependent differences in mitochondria-associated mRNA that correlate with aberrant appearing mitochondria in epithelium of Detrimental diet-fed sl/sl. A. Principal coordinate analysis shows epithelial RNA abundance patterns from only the distal colon of *sl/sl* mice fed the Protective (Prot) or Detrimental (Det) diets. Each symbol represents one mouse. B. Volcano plot showing log_2_(Fold-change) versus-log_10_(adjusted p-value) for *sl/sl* distal colon epithelial mRNAs. Green dots represent mRNA significantly more abundant in *sl/sl* fed the Protective diet. Magenta dots represent mRNA significantly more abundant in *sl/sl* fed the Detrimental diet. Dots above the horizontal dashed line have adjusted p value < 0.05 using Wald test and Benjamini and Hochberg correction in the DESeq2 package. C. Fgsea pathway analysis of diet-dependent differentially expressed genes (adjusted p <0.05) in distal *sl/sl* colon epithelial cells. Negative normalized enrichment scores indicate pathways with mRNA that were more abundant in *sl/sl* distal colon epithelial cells from mice fed the Protective diet. Positive enrichment scores indicate pathways with mRNA more abundant in distal colon epithelial cells of *sl/sl* fed the Detrimental diet. D. Enriched gene ontology (GO) for biological functions (BF) of genes differentially expressed in *sl/sl* distal colon epithelial cells depending on diet. E. Heatmap shows relative abundance of mRNA in distal colon epithelium of *sl/sl* differs based on diet for many genes involved in oxidative phosphorylation. F. ATP levels in distal colon intestinal epithelial cells differs based on genotype (*s/s* versus *sl/sl*) for mice fed the Detrimental diet (p = 0.044) but did not differ based on genotype for mice fed the Protective diet (p = 0.86), Two-tailed Mann-Whitney. G. Distal colon epithelium of *sl/sl* fed the Protective diet had more Hypoxyprobe® (pimidazole hydrochloride, PMDZ) staining (indicating a more hypoxic environment) than distal colon epithelium in *sl/sl* fed the Detrimental diet. P-value: * =0.0264 (Unpaired t-test) H. Transmission electron microscopy (TEM) shows distal colon epithelial cell mitochondria at 75,000x magnification for *s/s* or *sl/sl* fed the Protective or Detrimental diets. Scale bar = 200nm. White arrows indicate mitochondria. I. Mitochondria from TEM images of distal colon epithelium were blinded, classified, and scored for damage (adapted from Matsuzawa-Ishimoto 2017^166^). Distal colon epithelium of *sl/sl* fed the Detrimental diet had increased mitochondrial damage (Supplemental Figure 1) compared to *sl/sl* fed the Protective diet. P-value: * =0.0257 (Unpaired t-test). Mitochondria were classified as type 1 mitochondria (dense with cristae structure maintained), type 2 mitochondria (mildly swollen, mild loss of density with a 50% decrease in the number of visible cristae), or type 3 mitochondria (severely aberrant morphology with disorganized cristae and over 70% of cristae are missing). Distal colon epithelium of *sl/sl* fed the Detrimental diet had more type 3 mitochondria than distal colon epithelium of *sl/sl* fed the Protective diet. P-value: * =0.0199 (Unpaired t-test). N=3 for each genotype and diet. 6 images that spanned 3-4 distinct epithelial cells across different regions of the colon with at least 5 mitochondria per image were quantified for each animal.

**Table 5:** Top 10 differentially expressed mRNA more abundant in distal colon epithelial cells of sl/sl fed the Protective diet compared to *sl/sl* fed the Detrimental diet.

| Distal colon most differentially expressed genes, Protective Diet |  |  |  |  |
| --- | --- | --- | --- | --- |
| Gene name | Mean counts | log<sub>2</sub> Fold-difference in mRNA abundance | log<sub>2</sub> Fold-difference std. error | p<sub>adj</sub> |
| **AC151990.1** | 373.09 | 4.84 | 1.43 | 2.33E-02 |
| **Tgtp1** | 3175.32 | 3.88 | 1.11 | 1.90E-02 |
| **ligp1** | 2544.16 | 3.73 | 1.04 | 1.62E-02 |
| **Tgtp2** | 440.55 | 3.64 | 1.06 | 2.15E-02 |
| **Gm29013** | 262.92 | 3.45 | 0.97 | 1.67E-02 |
| **BC023105** | 45.69 | 3.42 | 1.08 | 3.34E-02 |
| **Sprr2h** | 83.00 | 3.28 | 1.04 | 3.40E-02 |
| **S100a8** | 38.32 | 3.22 | 0.88 | 1.42E-02 |
| **S100a9** | 56.03 | 3.09 | 0.96 | 3.13E-02 |
| **Il1b** | 112.99 | 2.74 | 0.84 | 2.88E-02 |

**Table 6:**
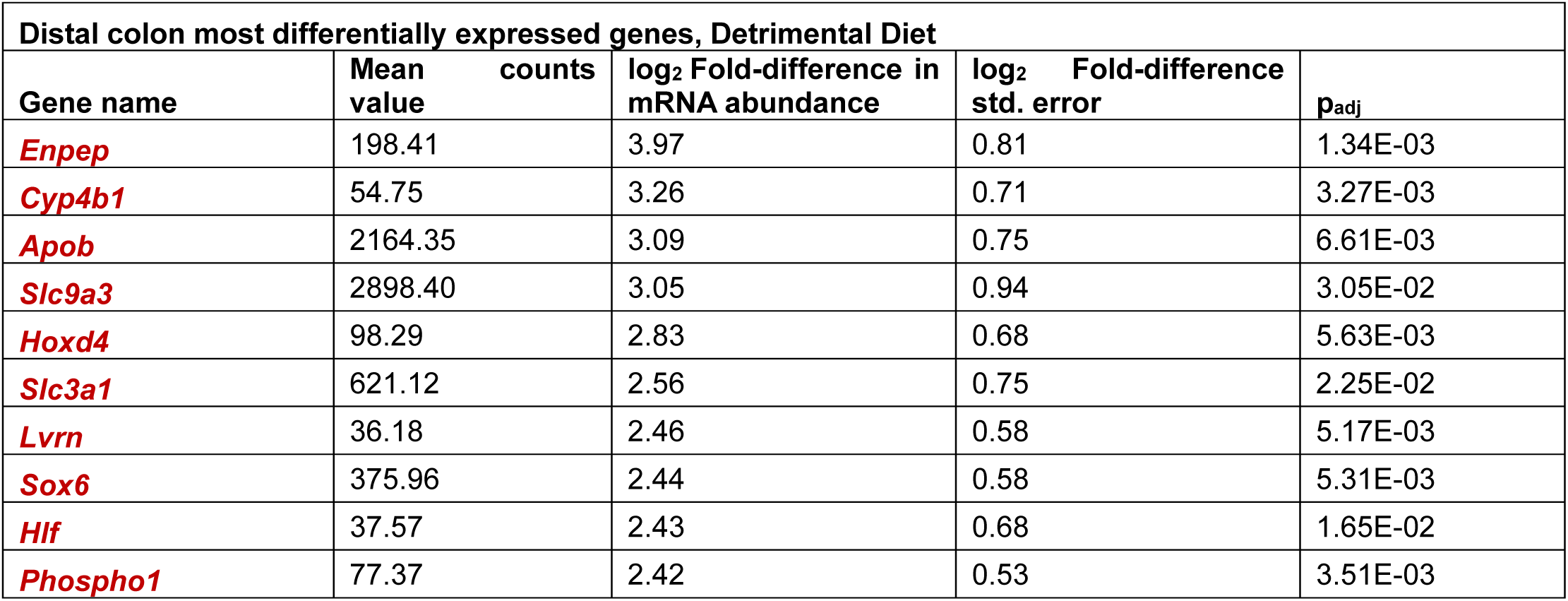
Top 10 differentially expressed mRNA more abundant in distal colon epithelial cells of *sl/sl* fed the Detrimental diet compared to *sl/sl* fed the Protective diet.

| Distal colon most differentially expressed genes, Detrimental Diet |  |  |  |  |
| --- | --- | --- | --- | --- |
| Gene name | Mean counts | log<sub>2</sub> Fold-difference in mRNA abundance | log<sub>2</sub> Fold-difference std. error | p<sub>adj</sub> |
| **Enpep** | 198.41 | 3.97 | 0.81 | 1.34E-03 |
| **Cyp4b1** | 54.75 | 3.26 | 0.71 | 3.27E-03 |
| **Apob** | 2164.35 | 3.09 | 0.75 | 6.61E-03 |
| **Slc9a3** | 2898.40 | 3.05 | 0.94 | 3.05E-02 |
| **Hoxd4** | 98.29 | 2.83 | 0.68 | 5.63E-03 |
| **Slc3a1** | 621.12 | 2.56 | 0.75 | 2.25E-02 |
| **Lvrn** | 36.18 | 2.46 | 0.58 | 5.17E-03 |
| **Sox6** | 375.96 | 2.44 | 0.58 | 5.31E-03 |
| **Hlf** | 37.57 | 2.43 | 0.68 | 1.65E-02 |
| **Phospho1** | 77.37 | 2.42 | 0.53 | 3.51E-03 |

Because these data suggested altered mitochondrial function in distal colon epithelium of *sl/sl* mice fed Protective versus Detrimental diets, we first compared relative epithelial ATP levels but found no diet-dependent difference (**Figure 5F**). There was, however, slightly less ATP in distal colon epithelium of *sl/sl* compared to *s/s* mice fed the Detrimental diet, of uncertain significance (**Figure 5F**). When mitochondria generate ATP by oxidative phosphorylation, oxygen is reduced and incorporated into water, leaving healthy colon epithelium relatively hypoxic. To evaluate distal colon epithelial oxygen levels, we used Hypoxyprobe® (pimonidazole (PMDZ)), which is reductively activated in hypoxic cells. Hypoxyprobe® staining showed colon epithelium was relatively oxygen-rich in Detrimental diet-fed *sl/sl* mice compared to Protective diet-fed *sl/sl* mice (**Figure 5G**). Furthermore, transmission electron microscopy revealed many morphologically aberrant and damaged mitochondria in distal colon epithelium of *sl/sl* mice fed the Detrimental diet (**Figure 5H, I**). In contrast, the distal colon epithelium of *sl/sl* mice fed the Protective diet had lower proportions of dysmorphic mitochondria and almost no mitochondrial damage was observed in *s/s* mice fed either diet (**Figure 5H, I**).

Collectively these data show potent effects of Protective versus Detrimental diets on proximal and distal colon epithelial transcriptome with diet-dependent differences in the abundance of mRNA encoding antimicrobial peptides, fucosyltransferases, lipid metabolism proteins, and mitochondrial oxidative phosphorylation associated proteins that might alter HAEC risk. Increased mRNA for some neutrophil and inflammasome associated proteins was also noted in Detrimental diet-fed *sl/sl* “epithelium”, but this could reflect increased numbers of inflammatory cells in our P20 bowel “epithelial cell” preparations, since we did not purify epithelial cells by flow sorting. Many diet-dependent differences in epithelial mRNA abundance differed depending on colon region (proximal versus distal colon). The elevated distal colon epithelial oxygen levels in *sl/sl* mice fed the Detrimental diet was particularly interesting since elevated oxygen levels could foster growth of inflammation-promoting facultative anaerobes like *Enterobacteriaceae*^67–69^.

### Detrimental diet promotes growth of *Enterobacteriaceae* in *sl/sl* mice while selectively depleting *Enterobacteriaceae* improves *sl/sl* survival

To evaluate diet and genotype-dependent effect on stool microbes, we pursued 16S rDNA sequencing using P20 stool collected at CHOP from mice fed the designated diet for at least 2 generations. Diet and genotype contributed to separations in microbial communities (p=0.023 by PERMANOVA) (**Figure 6A,6B**). This suggests that diet-dependent differences in stool microbial populations could impact *sl/sl* survival, a hypothesis supported by enhanced survival of Detrimental diet-fed *sl/sl* mice treated with broad spectrum antibiotics given from weaning (P21) till death (**Figure 6C**).

**Figure 6:**
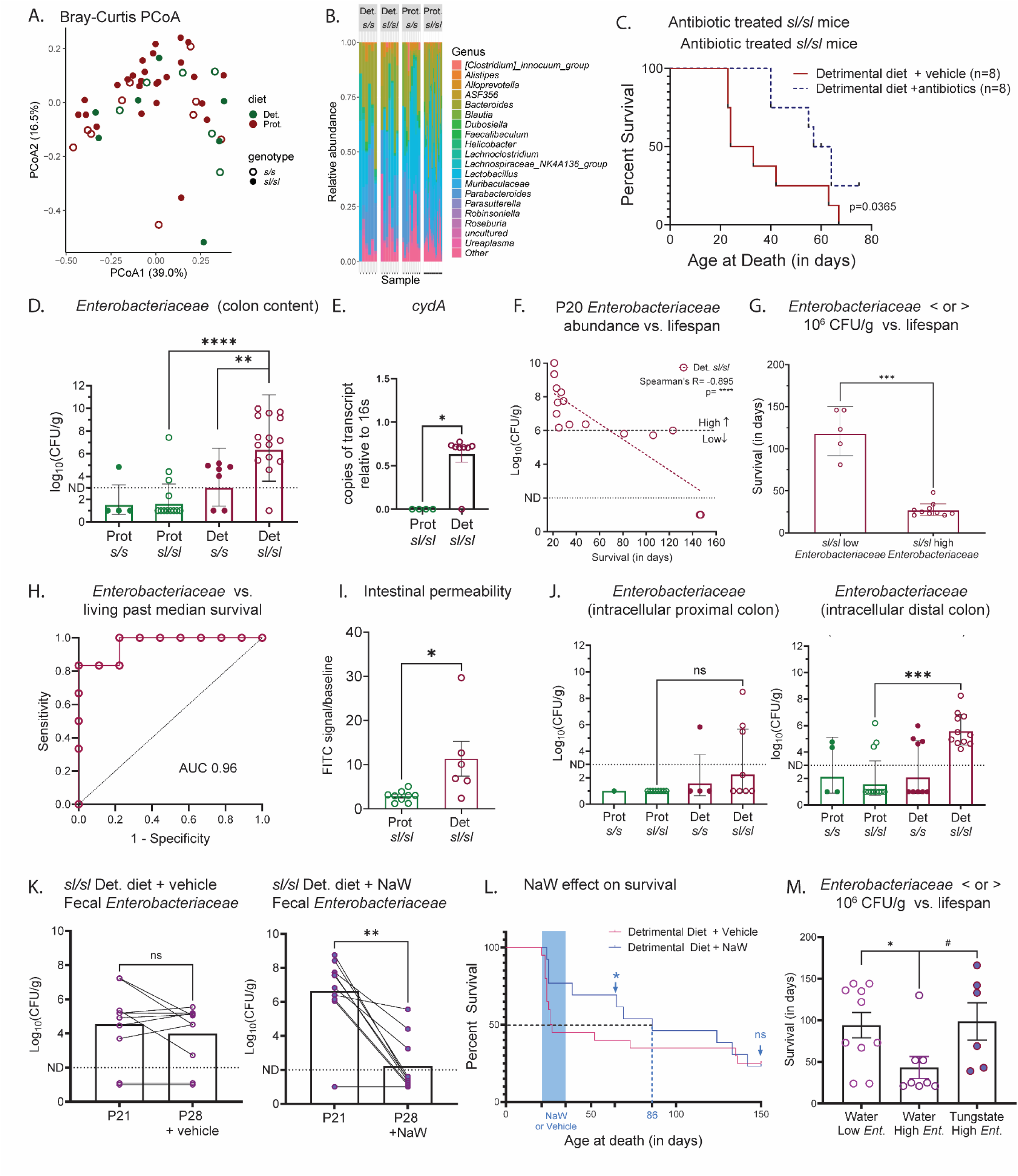
**Detrimental diet increases Enterobacteriaceae in sl/sl stool but reducing these pathobionts prolongs sl/sl survival**. A. Bray-Curtis principal coordinate analysis based on 16S rDNA sequencing results in microbial separation between genotypes, s/s and sl/sl, and diets, Protective and Detrimental, of stool microbial populations at CHOP. p=0.023 PERMANOVA, N = 49 mice. B. Relative taxonomic composition at the genus level between groups. C. Kaplan-Meier survival curve for *sl*/sl housed at CHOP and fed the Detrimental diet. N=8 sl/sl were given regular drinking water or while n=8 sl/sl received water with antibiotics from weaning at P21 until time of death. Antibiotic cocktail included neomycin, ampicillin, vancomycin, metronidazole, imipenem and ciprofloxacin. Gehan-Breslow-Wilcoxon test p<0.037. D. *Enterobacteriaceae* (quantified by culture on McConkey agar) in colon stool at P20 were more abundant in *sl/sl* mice fed the Detrimental diet compared to *sl/sl* fed the Protective diet. *Enterobacteriaceae* were also more abundant in stool of *sl/sl* compared to *s/s* fed the Detrimental diet. ** = 0.0015. **** = <0.0001, Two-tailed Mann-Whitney. Samples below the limit of detection (100 CFU/g) were plotted at 1. E. Relative abundance of bacterial mRNA for cytochrome oxidase d subunit I (*cydA*) in stool from distal colon of *sl/sl* fed the Protective (Prot) or Detrimental (Det) diet (normalized to *Enterobacteriaceae*-specific 16S rDNA copy number) based on qRT-PCR. P-value * = 0.0182, Two-tailed Mann-Whitney. F. *Enterobacteriaceae* abundance in stool at P20 (quantified by culture on McConkey agar) versus lifespan in days for *sl/sl* mice fed the Detrimental diet. All *sl/sl* with > 10^6^ *Enterobacteriaceae*/gram of stool at P20 died before 48 days of age. G. Survival (in days) for mice with more (high) or less (low) than 10^6^ *Enterobacteriaceae* per gram of colonic stool at P20. P-value ***= 0.0007, Two-tailed Mann-Whitney. H. Low *Enterobacteriaceae* abundance in colon stool at P20 (< 10^6^/gram) predicts that an *sl/sl* mouse will survive beyond the median age of death for *sl/sl* fed the Detrimental diet (P42) as shown in receiver operator characteristic (ROC) curve. I. Intestinal permeability is higher in *sl/sl* fed the Detrimental diet compared to *sl/sl* fed the Protective diet based on absorption of 4 kDa FITC dextran from bowel into the blood. Y-axis shows the ratio of FITC in serum 3 hours after gavage divided by fluorescence signal in serum before FITC dextran was administered. Serum was isolated from blood collected by cheek bleed. P-value * = 0.012, Two-tailed Mann-Whitney. J. Intracellular *Enterobacteriaceae* were more abundant in distal colon of *sl/sl* fed the Detrimental diet than in *sl/sl* fed the Protective diet based on resistance to gentamycin treatment of tissue *in vitro*. P-value *** = 0.004, Two-tailed Mann-Whitney. In contrast, in proximal colon there were very few intracellular *Enterobacteriaceae* in *s/s* or *sl/sl* independent of diet (ns). K. *Enterobacteriaceae* abundance decreases when *sl/sl* fed the Detrimental diet are supplemented with NaW in drinking water from P21 to P28 (right). P value **=0.0039, Two-tailed Mann-Whitney. In contrast, *Enterobacteriaceae* abundance does not change in *sl/sl* mice between P21 and P28 in the absence of NaW treatment (left). L. A brief period of sodium tungstate (NaW) treatment from P21 to P35 prolonged median *sl/sl* survival in mice fed the Detrimental diet from 26 days (without NaW, n=20) to 86 days (after NaW treatment, n=13). Survival benefits were transient, and equivalent between groups by P135. The wide blue vertical bar indicates the interval during which some mice received NaW treatment. The blue vertical dashed line is at median age of survival for the NaW treated group. Black horizontal dashed line indicates median survival. Log-rank (Mantel-Cox) test comparing NaW treated to control water treated *sl/sl* at P64 (p = 0.032) and P150 (p = 0.56). M. Early death that occurs in *sl/sl* with high (> 10^6^/gram) *Enterobacteriaceae* abundance at P20 could be prevented by NaW treatment. All *sl/sl* mice were fed the Detrimental diet. Low Ent. = stool *Enterobacteriaceae* abundance at P20 < 10^6^/gram. High Ent. = *Enterobacteriaceae* abundance at P20 > 10^6^/gram. P-values *=0.026, # = 0.0188, Two-tailed Mann-Whitney.

To identify relevant diet-dependent differences in microbial populations that could alter HAEC risk or predispose to early death, we directly cultured *Enterobacteriaceae* in stool using McConkey agar that selects for this family of microbes. *Enterobacteriaceae* are minority pathobionts in healthy microbiomes but commonly expand in other colitis models and predispose to bowel inflammation. For P20 *sl/sl* mice fed the Detrimental diet we cultured an average of 10^6^ colony forming units of *Enterobacteriaceae* per gram (CFU/g) of stool, which was below the limit of detection by conventional microbiota 16S rDNA profiling (**Figure 6D**)^70^. In contrast, stool from P20 *s/s* mice cohoused and fed the Detrimental diet had an average of 10^3^ stool *Enterobacteriaceae* (CFU/g). Strikingly, many *s/s* and *sl/sl* mice fed the Protective diet had almost undetectable fecal *Enterobacteriaceae* counts at P20 based on MacConkey agar culture. The dysbiotic expansion of facultative anaerobes like *Enterobacteriaceae* that occurs in an oxygen-rich colon environment is facilitated by bacterial utilization of cytochrome bd-I oxidase (cydA) (encoded by the CydABX operon) in the inflamed intestine^71–73^. Consistently, we detected increased transcripts of *cydA* mRNA in distal colon stool from *sl/sl* mice fed the Detrimental diet using *Enterobacteriaceae*-specific primers, but could not detect *cydA* mRNA in stool from *sl/sl* mice fed the Protective diet (**Figure 6E**). By plotting survival as a function of P20 stool *Enterobacteriaceae* counts (**Figure 6F**) we noticed that all Detrimental diet-fed *sl/sl* mice with > 10^6^ *Enterobacteriaceae* CFU/g at P20 died by 48 days of age (median survival 28 days). In contrast, Detrimental diet-fed *sl/sl* mice with < 10^6^ CFU/g *Enterobacteriaceae* in stool at P20 had a median survival of 120 days, and two Detrimental diet-fed *sl/sl* mice survived 146 days. The impact of P20 stool *Enterobacteriaceae* abundance is highlighted by a binary classification (**Figure 6G**) and a receiver operator characteristic curve that predicts survival past 42 days using 10^6^ *Enterobacteriaceae* CFU/g of stool as a threshold (42 days = median survival for all Detrimental diet-fed *sl/sl* mice) (**Figure 6H**).

We next hypothesized that the abundant *Enterobacteriaceae* promote inflammation and cause early death by penetrating the *sl/sl* bowel epithelial barrier, which is more permeable to small molecules (e.g., 4 kDa FITC-dextran) in Detrimental diet-fed *sl/sl* than in *sl/sl* fed the Protective diet (**Figure 6I**). To test this hypothesis, we treated colon *in vitro* with gentamycin to kill *Enterobacteriaceae*, recognizing this antibiotic does not penetrate mammalian cells efficiently. McConkey agar culture demonstrated high levels of gentamycin-resistant (i.e., intracellular) *Enterobacteriaceae* in distal colon of Detrimental diet-fed *sl/sl* (mean = 475,335 ± 3.89 CFU/g colon), compared to *sl/sl* fed the Protective diet (mean = 131 ± 2.38 CFU/g colon) (**Figure 6J**). Taken together, these results suggest the Detrimental diet promotes gut microbiome dysbiosis in *sl/sl* mice, specifically *Enterobacteriaceae* overgrowth and that the Detrimental diet enhances epithelial translocation of potentially pathogenic bacteria.

We next asked whether the bloom of *Enterobacteriaceae* drove early death in Detrimental diet-fed *sl/sl* mice, or if elevated *Enterobacteriaceae* at P20 simply allow us to identify *sl/sl* predisposed to early death for other reasons. To answer this question, we supplemented *sl/sl* fed the Detrimental diet with sodium tungstate (NaW) from P21 to P35 to selectively decrease *Enterobacteriaceae* abundance during this critical period of susceptibility by counteracting molybdoenzyme-dependent respiration^74, 75^. McConkey agar plating confirmed that NaW treatment reduced *Enterobacteriaceae* counts from a median of 15.5 x 10^6^ CFU at P21 to undetectable median levels by P28 (**Figure 6K**). In this cohort, 55% of untreated Detrimental diet-fed *sl/sl* mice died between P21 and P27, while only 26% of NaW-treated *sl/sl* mice died during the same interval (**Figure 6L**). The effects of transient NaW treatment were still observable at P70 (p=0.034) but no longer apparent by P142 (**Figure 6L**), possibly due to diet-induced regrowth of *Enterobacteriaceae*. Of note, effects of NaW on *sl/sl* survival were particularly evident for Detrimental diet-fed *sl/sl* mice with greater than 10^6^ *Enterobacteriaceae* CFU/g counts at P20 (**Figure 6M**)^56^, highlighting the potential to use this family of microbes as a biomarker for treatment response. Collectively, these data suggest that the Detrimental diet fosters the growth of *Enterobacteriaceae* and that high levels of *Enterobacteriaceae* reduce survival for *sl/sl* mice.

## Discussion

Hirschsprung disease is a first trimester birth defect defined by the absence of enteric nervous system cells in distal bowel^2–6, 32^. For reasons that remain incompletely understood, most children with HSCR (∼65%) develop bowel inflammation before and after pull-through surgery (i.e., HAEC)^9, 10, 12, 21, 76^. HAEC predisposes to sepsis^77–80^, the most common cause of HSCR-associated death in childhood. HAEC also causes significant morbidity, repeated hospitalizations, and recurrent HAEC is often treated by “re-do” pull-through surgery or ostomy^14, 16, 81–83^. Furthermore, 1-2% of children with HSCR are eventually diagnosed with inflammatory bowel disease (IBD)^84^. Half of these IBD/HSCR diagnoses occur before 5 years of age (defined as “very early onset (VEO) IBD”). In contrast, for the non-HSCR population, only ∼1-2 per 1000 children are diagnosed with IBD and only 4% of these children (4/100,000) have VEO IBD^84, 85^. These observations suggest that the absence of the ENS profoundly alters intestinal barriers that reduce inflammation and prevent sepsis, but how this occurs is not known.

To learn more about HAEC, we took advantage of our serendipitous observation that *Piebald lethal* (*sl/sl*) and *Holstein* (*Hol^Tg/Tg^*) HSCR-like mouse models live longer in Philadelphia than in Montreal (see **Supplemental Figure 10C and 10D** is our previous manuscript^60^). Effects on survival were evident within 2 generations after we mutually exchanged *Piebald letha*l and *Holstein* mouse lines, but were more dramatic for *sl/sl* than for *Hol^Tg/Tg^*, so we studied *sl/sl*. While many aspects of mouse vivaria alter phenotypes^86–92^, we hypothesized that different mouse diets at our institutions might be a key factor driving survival differences. By feeding the standard CHOP diet to *sl/sl* mice in Montreal and the standard UQAM diet to *sl/sl* mice at CHOP, we discovered for the first time (to the best of our knowledge) that diet significantly alters survival in a HSCR model. Survival curves also differed by location independent of diet, suggesting other colony factors may affect survival. We suspect this location-dependent survival difference is due at least in part to differences in colony-specific gut microbes, but we have not yet tested this hypothesis. Remarkably, there is almost nothing known about the impact of diet on HSCR and no formal recommendations about diet for children with HSCR or HAEC, although parents of children with HSCR report diet impacts their children’s symptoms^46, 47^.

The diets we studied are “standard mouse chow” used at many institutions. Although we refer to LabDiet Mouse Diet 5015 (LD-5015) as “Protective” and Charles River Rodent Diet 5075 (CR-5075) as “Detrimental”, these judgmental terms are apt descriptors (as far as we know) only for mice with distal colon aganglionosis mimicking human HSCR. In our colonies, *sl/sl* mice fed either diet had similar lengths of distal colon aganglionosis, but at P20 differed in fecal metabolites, stool microbes (very selectively), epithelial permeability, epithelial gene expression, distal colon epithelial oxygen levels, and epithelial mitochondrial morphology. Many of the Detrimental diet-driven changes in mammalian biology may promote fecal *Enterobacteriaceae* overgrowth in *sl/sl* colon and could increase intracellular *Enterobacteriaceae* as we found based on resistance to gentamycin treatment of colon *in vitro*. *Enterobacteriaceae* overgrowth is a hallmark of colitis in other model systems^93, 94^, and selective depletion of *Enterobacteriaceae* prolonged survival of *sl/sl*, consistent with the hypotheses that *Enterobacteriaceae* promote HAEC and cause early death when ENS is absent. These murine observations fit with limited human data showing increased *Proteobacteria* (a phylum that includes *Enterobacteriaceae*) in children with HAEC^23, 95, 96^.

### Dietary and fecal metabolites that might be protective or detrimental

Fecal metabolites derived from diet or microbial metabolism are well known to impact epithelial and immune cell biology in the bowel^97–102^. Among the metabolites most markedly increased in stool of *sl/sl* mice fed the Protective diet (compared to Detrimental diet-fed *sl/sl* mice) were 3,5-dihydroxybenzoic acid (16.84-fold higher), butyrate/isobutyrate (4.47-fold higher), valerate (3.83-fold higher), and 1-stearoyl-2-oleoyl-GPS (4.94-fold higher). 3,5-dihydroxybenzoic acid is an agonist for GPR81^103^, the lactate receptor that protects against colitis^104^. Butyrate and valerate^105^ are short chain fatty acids that reduce colitis severity, but butyrate is most well studied. Butyrate is a primary energy source for colon epithelial cells^106, 107^, alters gene expression by inhibiting histone deacetylases (HDACs)^106, 108, 109^, reduces NFκB signaling^110–112^, blocks interferon gamma signaling^109, 113^, increases epithelial production of anti-microbial peptides^114, 115^, acts as a growth factor^116^, strengthens the colon epithelial barrier^97, 106, 117^, activates peroxisome proliferator-activated receptor gamma^118–120^, and increases IL-22 receptor expression in colon epithelium^121, 122^. Butyrate also increases *IL22* mRNA in innate lymphoid cells type 3 (ILC3) and CD4+ T cells in a HIF1α and hypoxia-dependent manner^106, 122^. In addition, butyrate supports regulatory T cells^100, 123^ and polarizes macrophages toward an anti-inflammatory “M2” phenotype^124–126^. Finally, because butyrate metabolism in mitochondria requires oxygen, epithelial butyrate utilization should reduce epithelial oxygen levels. The low oxygen enhances HIF1α dependent gene expression^127, 128^ and facilitates the growth of beneficial obligate anaerobes near colon epithelium, including microbes that produce more butyrate^106, 129, 130^. Thus, many of the beneficial effects of the Protective diet might be due to elevated colon fecal butyrate, valerate and 3,5-dihydroxybenzoic acid, although these metabolites were only statistically elevated in stool of Protective diet-fed *sl/sl* at UQAM (and not at CHOP) once we corrected metabolomic data for multiple comparisons inherent in untargeted metabolomics. We note that mean diet-dependent difference in abundance was similar for many metabolites at both locations, but p-values differed by location, suggesting increased variability in the CHOP dataset (**Figures 3B, D, E, Table 2**). In contrast to Protective diet-fed stool, the stool of *sl/sl* fed the Detrimental diet was relatively enriched in putrescine, a metabolite reported by Grosheva *et al.* to damage colon epithelial tight junctions, increase gut permeability, increase *C. rodentium* attachment to colon epithelium, increase colon inflammatory cytokines, and increase epithelial mRNAs that regulate metal binding, cytoskeletal organization and oxidative stress^99, 131^. In contrast, Nakamura *et al*. reported putrescine reduces the severity of colitis in the dextran sodium sulfate model, increases anti-inflammatory macrophages in colon, facilitates colon epithelial cell proliferation, and enhances autophagy and oxidative phosphorylation in colon epithelial cells^132^. This suggests Protective versus Detrimental effects of putrescine in colitis models might be context or dose dependent. Furthermore, when CHOP data were analyzed in isolation, relative abundance of putrescine in Protective versus Detrimental diet-fed *sl/sl* were not statistically different after multiple comparison testing.

The reasons that Protective diet-fed *sl/sl* have elevated butyrate in stool at P20 (at least at UQAM) are not known. Butyrate/isobutyrate were not detected by untargeted mass spectrometry in either diet but are usually generated within the colon by microbial metabolism of fiber or some amino acids. Because fiber content of the Detrimental diet is twice as high as in the Protective diet (**Table 1**), it is unlikely that fiber is the primary source of elevated butyrate in Protective diet-fed *sl/sl* mice. Consistent with this hypothesis, we did not detect diet-dependent differences in fecal abundance of common fiber metabolizing butyrate-producing microbes (e.g., *Lachnospiraceae* like *Roseburia intestinalis* and *Eubacterium*, or *Faecalibacterium prausnitzii*,)^98^, suggesting butyrate is not primarily fiber-derived at P20 in the *sl/sl* model. However, butyrate can also be produced from glutamine by *Clostridium*, *Fusobacteriales*, and *Peptococcus* species^133,134^ (and perhaps other organisms) and glutamine is 83-fold more abundant in the Protective diet than in the Detrimental diet suggesting increased substrate (glutamine) availability might drive butyrate synthesis. An alternative hypothesis to be explored in future studies is that differences in *sl/sl* fecal butyrate at P20 are driven by diet-induced differences in breastmilk composition. Butyrate is present in breastmilk (at least in humans^135^ and goats^136^) and butyrate can be produced from breast milk oligosaccharides that also foster growth of butyrogenic microbes^137^, but we do not know how diets might alter breastmilk composition. Since mice typically start consuming chow at P14 and were weaned at P21, the fecal metabolites we measured at P20 might be altered by chow the pup consumes and/or changes in breastmilk composition that depend on maternal diet. We also do not yet know why relative butyrate abundance differs at CHOP compared to UQAM but suspect this reflects differences in gut microbes at the different vivaria, a topic for future study.

### Diet-dependent differences in mammalian biology that might be protective or detrimental

In the proximal colon, *sl/sl* mice fed the Protective diet had higher levels of epithelial mRNA for antimicrobial peptides (*Reg3b*, *Reg3g*) and the fucosyltransferase *Fut2*, compared to Detrimental diet-fed *sl/sl* mice. These genes, which prevent dysbiosis-induced colon inflammation^138^, and several other diet-induced differences in epithelial gene expression (*Lrg1*, *Pl2g2a*, *Socs3*, and *Tifa*) could be explained by differences in interleukin-22 (IL-22), which regulated expression or these mRNA ^139^. IL-22 synthesis in turn is regulated by butyrate and hypoxia in ILC3 and CD4 T cells as noted above. Future work involving modulation of IL-22 levels in the *sl/sl* model should help to address this hypothesis. Another striking observation was that Detrimental diet-fed *sl/sl* mice had abnormal appearing mitochondria compared to Protective diet-fed *sl/sl* mice or cohoused *s/s* mice fed either diet. Consistent with these findings, distal colon epithelial RNAseq showed marked reductions in many transcripts associated with oxidative phosphorylation in Detrimental diet-fed *sl/sl* mice compared to Protective diet-fed *sl/sl* mice. Mitochondrial bioenergetics is critical for intestinal homeostasis^140^, including maintaining barrier function^106, 141, 142^ and to maintain physiological hypoxia^106^ that suppresses aerobic expansion of facultative anaerobes like *Enterobacteriaceae*. These data raise fascinating and largely unexplored questions about how enteric neurons and glia impact colon epithelial intermediary metabolism, another topic for future study.

### Future directions

Hirschsprung disease-associated enterocolitis is an under-investigated form of bowel inflammation that is common and can be fatal in children with distal bowel aganglionosis. We (and others)^2, 7, 9, 10, 143–147^ hypothesized that enterocolitis occurs in people with HSCR because enteric neurons and glia critically control colon epithelial repair and proliferation^148–150^, mucin production and quality^151–154^, epithelial ion transport, bowel motility, local blood flow, and mucosal immune cell function among other ENS-specific effects on bowel biology^7, 155, 156^. We were therefore surprised that many diet-induced differences in mammalian and microbial biology that seem likely to prevent (or increase the risk of) early death in *sl/sl* mice recapitulate well-established disease mechanisms that impact the severity of inflammatory bowel disease, infectious colitis, and necrotizing enterocolitis. This may explain why HSCR increases very early onset IBD diagnosis by about 2000-fold, although what distinguishes HAEC from IBD in children with HSCR is not well defined. Interestingly, once children with HSCR are diagnosed with IBD, they are treated with a wide range of IBD-focused therapies^84^ that have never been evaluated in “non-IBD HAEC” including corticosteroids, aminosalicylates, immunomodulators, and biologic therapy (e.g., anti-TNF antibodies or others). If these medicines were effective to treat “non-IBD HAEC”, it might reduce the need for “re-do” pull-through surgery. The impact of diet on human HSCR is also almost entirely unexplored, but these data suggest diet-based therapy could markedly impact HAEC incidence or severity if critical dietary factors could be identified. Furthermore, since almost all our mechanistic studies focus on P20, before any *sl/sl* mice begin to die from complications of aganglionosis, it will be critical to understand what occurs after P20 that causes death in the context of aganglionosis. In addition, the impact of maternal diet in HSCR is completely unexplored, and protective effects of breastfeeding in HSCR deserve more study, since exclusive breastfeeding appears to markedly reduce HAEC incidence^21^ and diet could affect breastmilk composition. Finally, the impact of ENS function on epithelial metabolism, mitochondrial function, and epithelial energetics deserves more investigation, since these links could be manipulated to reduce colitis severity in HSCR and other diseases. The emerging fields of mucosal neuroimmune and neuroepithelial biology may therefore provide new answers not only about HSCR and HAEC, but also relevant for IBD, infectious colitis, cancer biology, and necrotizing enterocolitis.

## Abbreviations

AMPs: Antimicrobial peptides
ANOVA: Analysis of variance
BEH: Waters ethylene hybrid bridged particle technology
BH: Benjamini-Hochberg
CHOP: The Children’s Hospital of Philadelphia Research Institute
DAPI: 4’,6-Diamidino-2-phenylindole
DMSO: Dimethyl sulfoxide
DPBS: Dulbecco’s phosphate-buffered saline
EDTA: Ethylenediaminetetraacetic acid
ENS: Enteric nervous system
ENCDC: Enteric neural crest derived precursors
F2: Generation of mice fed a specific diet for exactly 2 generations (i.e. grandparents were weaned to the designated diet)
FA: Formic acid
FBS: Fetal bovine serum
FITC: Fluorescein isothiocyanate
g: Guage, referring to the size of a needle
HAEC: Hirschsprung disease associated enterocolitis
HBSS: Hank’s balanced salt solution
H&E: Hematoxylin and eosin
HILIC: Hydrophobic interacting chromatography
HSCR: Hirschsprung disease
IACUC: Institutional Animal Care and Use Committee
min: Minutes
LAN: Local area network
LIMS: Laboratory information management system
MS: Mass spectrometry
MS/MS: Tandem mass spectrometry
m/z: Mass to charge ratio
p: probability
P (followed by a number): Postnatal day
PBS: Phosphate-buffered saline
PBST: PBS with Triton X-100 at 0.5%
PCR: Polymerase chain reaction
PFA: Paraformaldehyde
PFPA: Perfluoropentanoic acid
QA: Quality assurance
QC: Quality control
RI: Retention time/index
RSD: Relative standard deviation
RT: Room temperature
s: seconds
SEM: Standard error of the mean
SSL/LeJ: Mouse strain name
UPLC: Ultrahigh performance liquid chromatography
UQAM: University of Quebec at Montreal

## Acknowledgements

Figures created with BioRender.com. We would like to acknowledge Briana Christophers for her early contributions to this project.

## Funding

This research was funded by NIH RO1: DK129691-04 (ROH), American Neurogastroenterology and Motility Society (ANMS) Discovery Grants Program (NT), NIH/NIDDK T32 DK101371 (NT), NIH L40DK134022 (NT), North American Society for Pediatric Gastroenterology, Hepatology and Nutrition (NASPGHAN) Reckitt Mead Johnson Nutrition Research Young Investigator Development Award (NT), the Irma and Norman Braman Endowment, and the Suzi and Scott Lustgarten Center Endowment (ROH), the Children’s Hospital of Philadelphia Frontier Program Center for Precision Diagnosis and Therapy for Pediatric Motility Disorders. Hartwell Foundation (ML) and NIH F32: DK139723 (ML).

## Author contributions

Conceptualization, NBT, MJL, NL, SSch, JE, RR, CT, NP, ROH; Methodology, NBT, MJL, NL, SEA, JR, SSax, SSch, JE, CT, NP, ROH; Investigation, NBT, MJL, NL, SSax, SSch, JE, ML, EL, YZ, LD, EF, DJ, SEA, JR, GY, ZT; Formal analysis, MJL, KB, LL, LD; Writing Original draft, NBT, ML; Writing, Reviewing & Editing ROH, NP, CT; Visualization, NBT, MJL; Funding Acquisition, ROH, CT, NP, NBT, MJL; Supervision, ROH, NP, CT, RR

## Methods

### Key resources table

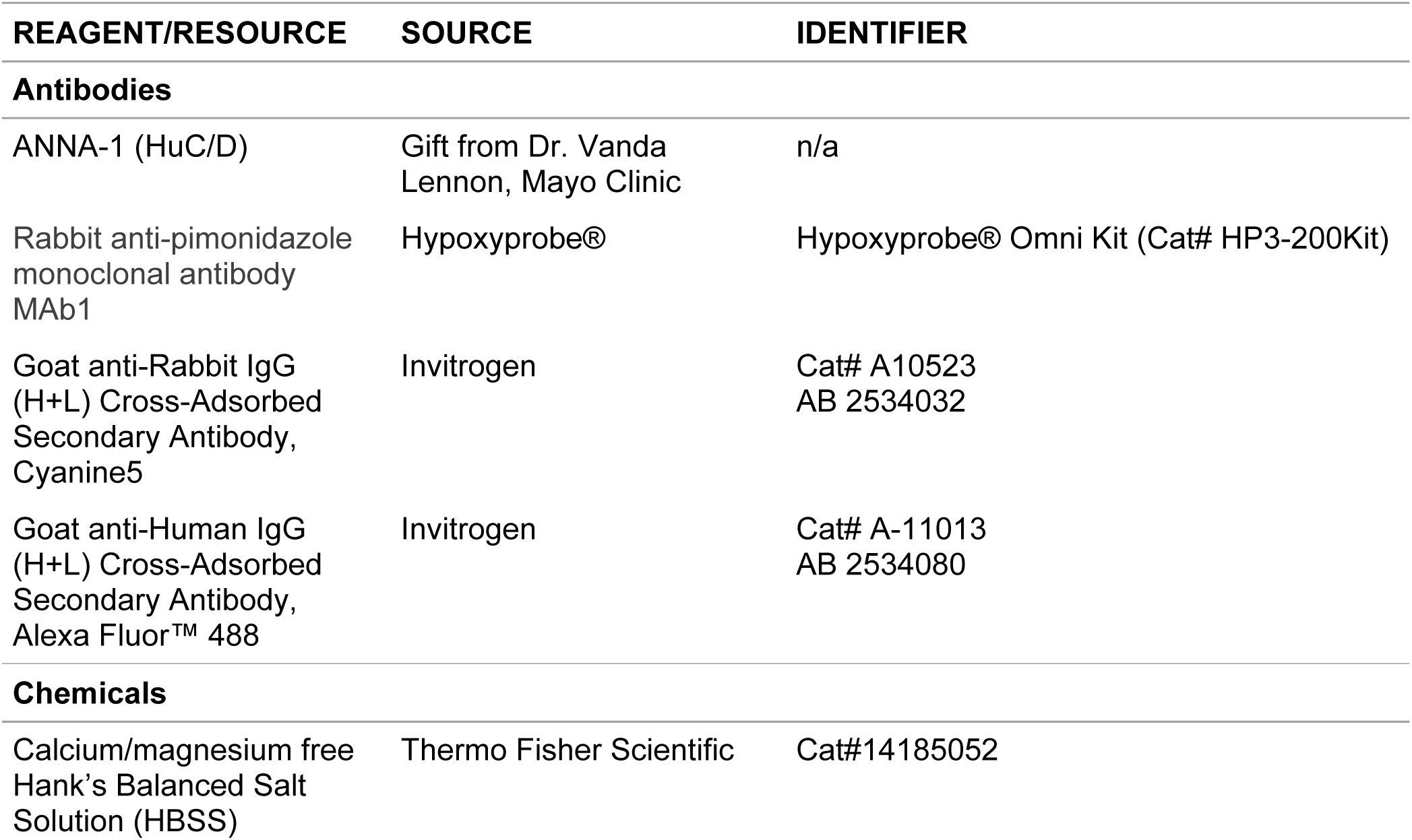
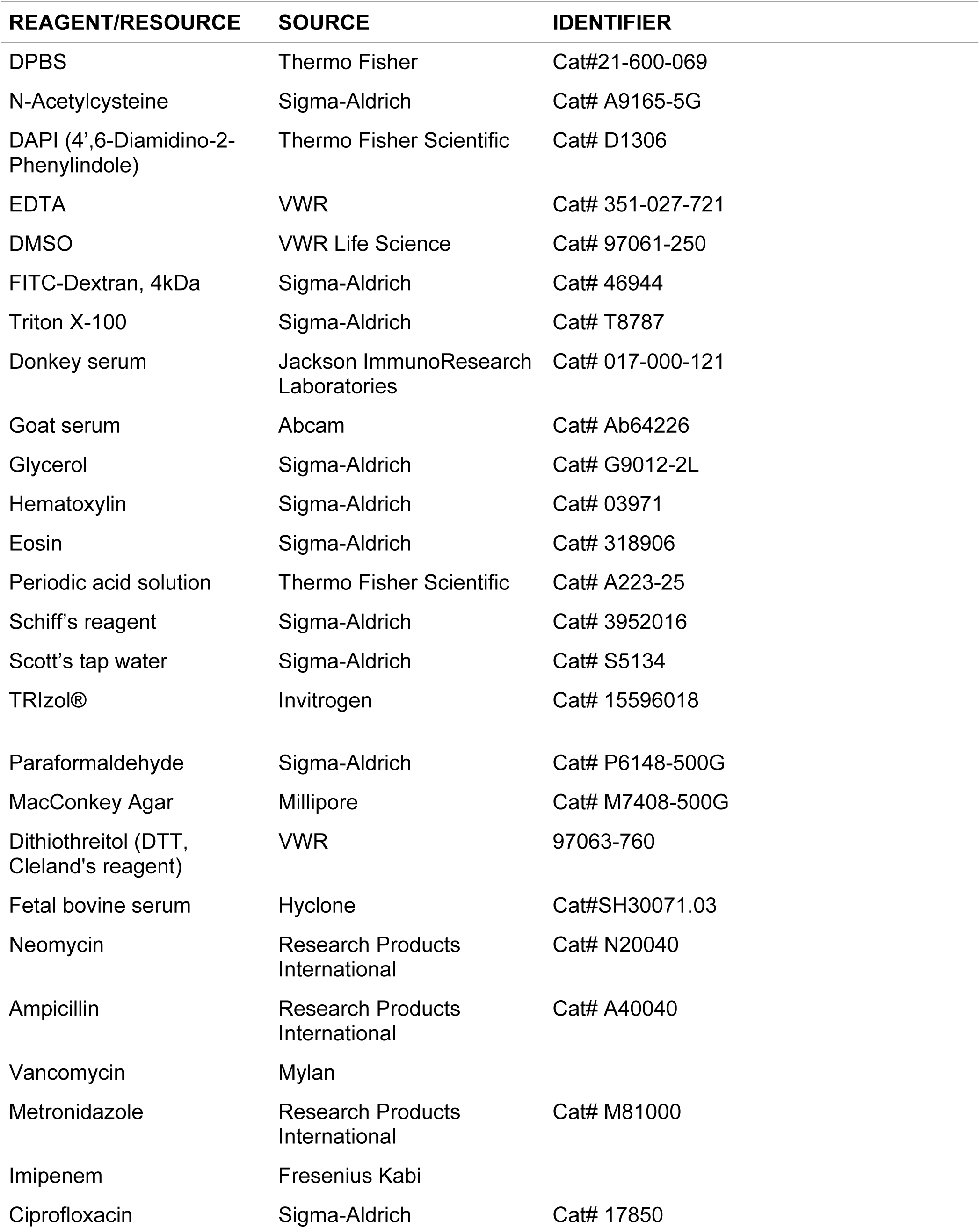
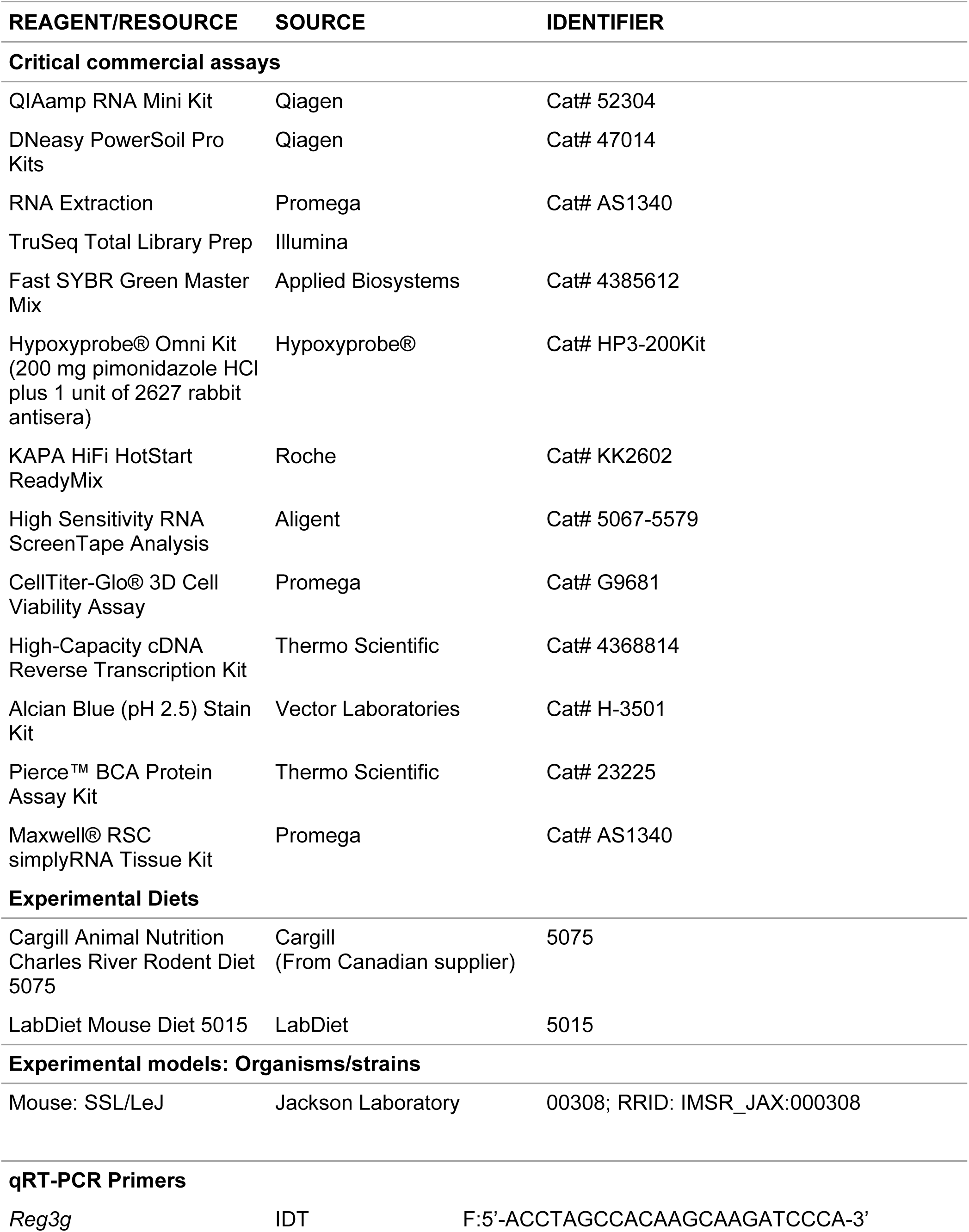
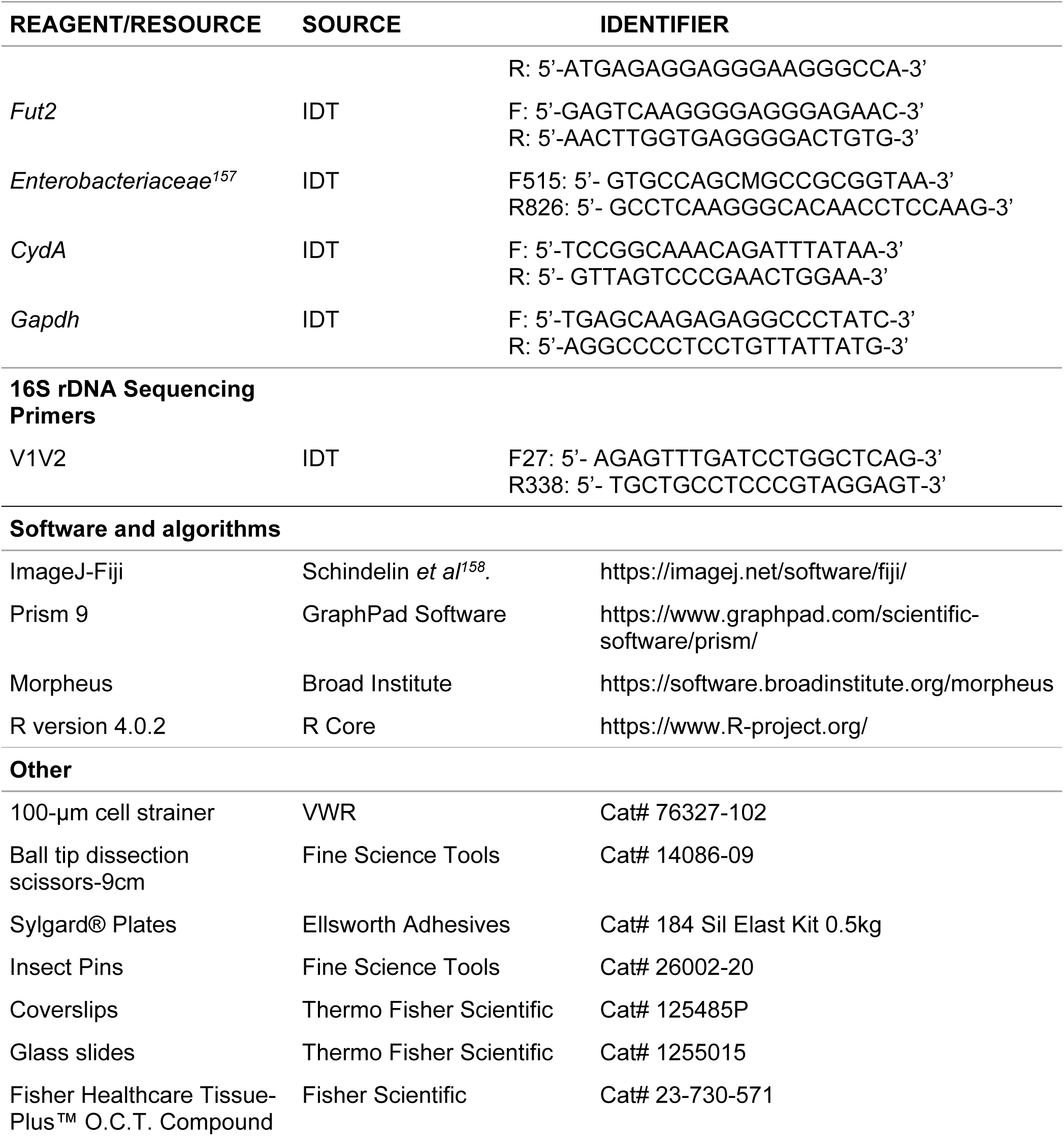

## Resource availability

### Lead contact

Further information and requests for resources and reagents should be directed to and will be fulfilled by the lead contacts, Naomi Tjaden, Megan Liou and Robert O. Heuckeroth.

### Materials availability

This study did not generate new unique reagents.

### Data and code availability

- The raw and processed RNA-seq data will be deposited in the NCBI Gene Expression Omnibus at time of submission to a peer-reviewed journal.
- Other data will be shared by the lead contacts upon request.
- R scripts are available from the lead contacts upon request.
- Additional information required to reanalyze data is available from lead contacts upon request.

### Mice

All procedures were conducted with the approval of the Institutional Animal Care and Use Committee (IACUC) at The Children’s Hospital of Philadelphia Research Institute (IAC 22-001041), The University of Pennsylvania (IAC 806361), and University of Quebec at Montreal (CIPA-959). New weanling mice (P19-21) of either sex were used for most studies. All mice were maintained in a standard 12h light: 12h dark cycle at 72°F ± 2 degrees with *ad libitum* access to standard chow (Cargill Animal Nutrition Charles River Rodent Diet 5075 or LabDiet Mouse Diet 5015) and water. Cage enrichment at both locations: House/dome (Bioserve, S3174) and nestlet (Ancare). *sl/sl* and *s/s* mice used in all studies were maintained on the specified diet for at least two generations (i.e., their grandparents (or earlier generations) were weaned to the designated diet. *SSL/LeJ* (*Piebald lethal*) mice were from The Jackson Laboratory (*Ednrb^s-l^*; JAX stock #000308; C3H/HeJ-C57BL/6 mixed background). Mice were maintained by breeding *s/sl* x *s/sl* genotypes. Bedding at CHOP: ¼ inch corn cob (The Andersons, Product 4B) (2018-2020); Shepherd’s cob blend (50:50 corncob + ALPHA-dri, Shepherd Specialty Papers) (2021-present). Bedding at UQAM: Beta Chip, heat treated hardwood lab bedding (NEPCO).

### Euthanasia

Mice were euthanized according to IACUC protocols. Mice at CHOP were euthanized by terminal inhalation of carbon dioxide from a compressed gas cylinder followed by the secondary physical method of cervical dislocation or cardiac puncture, or alternatively primary physical method of cervical dislocation followed by exsanguination of a vital organ (liver or heart). Mice at UQAM were euthanized after isoflurane anesthesia, followed by terminal inhalation of carbon dioxide followed by a secondary physical method of cervical dislocation or cardiac puncture.

### Necropsy and histopathology

*Piebald lethal* mice that died during survival studies at CHOP and UQAM were evaluated by necropsy. Moribund (hunched, dehydrated, hypothermic, with ruffled fur) mice were euthanized and subjected to gross necropsy, since mice that met moribund criteria died within 24 hours without intervention. Histopathology studies were performed at necropsy to evaluate proximal and distal colon. *Piebald lethal* mice aged P19-P21 were euthanized for RNA sequencing, histologic analysis, and qRT-PCR. Immunohistochemical, histochemical, and electron microscopy images were reviewed blinded.

### Hematoxylin and eosin (H&E) staining

Proximal and distal colon (intact without attempts to clear stool) were fixed in 4% paraformaldehyde (PFA) overnight and paraffin embedded. 5 μm sections cut on an HM 355S Microm microtome were stained using standard a H&E protocol. Briefly, sections were deparaffinized and rehydrated, before nuclei were stained by hematoxylin for 3 min, differentiated by acid alcohol for 30 s and then eosin stained was for 10 s. Stained sections were dehydrated and sealed with a neutral resin. This staining was performed at the Center for Molecular Studies in Digestive and Liver Diseases (P30DK050306, RRID: SCR_022420).

### Alcian blue staining for goblet cell, mucus layer, and crypt length quantification

1 cm colon containing a fecal pellet was fixed in Carnoy’s fixative (60% 200 proof ethanol, 30% chloroform, 10% glacial acetic acid) overnight and transferred to 100% ethanol before paraffin embedding at the Molecular Pathology and Imaging Core at Penn (P30DK050306, RRID: SCR_022420). 7μm sections were deparaffinized and stained with Alcian Blue (pH 2.5) Stain Kit (H-3501) according to manufacturer’s guidelines. Stained sections were visualized using a Nikon Ti2 microscope. Goblet cells were enumerated from full crypts and averaged per mouse. Crypt length from full crypts was measured using NIS-Elements Advanced Research software and averaged per mouse.

### Enterocolitis Scores

Enterocolitis scores from 0-3 were adapted from Cheng et al^64^. H&E-stained sections from proximal ENS-containing colon and distal aganglionic colon were scored by a pathologist who did not know the origin of the specimens (*i.e*., blinded). Scale: 0: no inflammation, rare neutrophil. 1: mild inflammatory infiltrates, no necrosis. 2: moderate to marked inflammatory infiltrates and mucosal necrosis. 3: transmural necrosis.

### Fecal pellet collection

Fecal pellets for metabolomic analysis and sequencing were collected at each timepoint, flash frozen on dry ice and stored at-80C until analysis. Pellets were either spontaneously passed just prior to sacrifice or taken from the distal colon upon dissection.

### 16S rDNA Library Preparation and Microbiome Analysis

DNA was amplified using the KAPA Hifi HotStart ReadyMix (Roche, KK2602) and the F27/R338 primer-pair targeting the V1V2 region of the 16S gene (F27: 5’-AGAGTTTGATCCTGGCTCAG-3’, R338: 5’- TGCTGCCTCCCGTAGGAGT-3’). After PCR amplification, the pooled library underwent beadbased size selection using AMPure XP beads (Beckman Coulter, A63881) and was sequenced using 500 bp paired-end sequencing (Illumina MiSeq). QIIME 2 (version 2021.2.0^159^) was used to process 16S sequencing reads. Briefly, reads were demultiplexed^160^, quality-filtered^161^, and denoised using deblur with ‘trim-length 200’^162^. For taxonomic classification, reads were extracted from the Silva 138 database^163, 164^ using the following parameters: ‘--p-f-primer AGAGTTTGATCCTGGCTCAG--p-r-primer TGCTGCCTCCCGTAGGAGT--p-trunc-len 200’. These reads were used to train a naïve Bayes classifier that was then applied to our dataset. Downstream analysis was performed in R version 4.1.0 using tidyverse, phyloseq, and vegan packages^165^.

### Wholemount colon immunohistochemical staining of enteric neurons

Mouse full-length colons from postnatal day 19-21 mice were cut along the mesentery with ball tip scissors in ice cold PBS. The colon was pinned flat luminal side down on a Sylgard® 184 Elastomer (Ellsworth Adhesives, catalog 184 SIL ELAST KIT 0.5KG) using insect pins (Fine Science Tools, catalog 26002-20), fixed while pinned in 4% PFA (Sigma-Aldrich, catalog P6148-500G) (30 minutes, room temperature (RT)), washed 3 times with PBS (5 minutes), bisected into “proximal” and “distal” pieces, then stored in 50% glycerol/50% PBS with 0.05% sodium azide (Sigma-Aldrich, catalog G9012-2L) at room temperature (30 minutes) before-20°C storage. Colon was split into proximal and distal pieces for improved antibody penetration and imaging. Muscle layer with myenteric plexus was separated from the mucosa and submucosa after fixation. Prior to staining of the distal muscle layer piece, frozen tissue was thawed (RT), rinsed in 1x PBS (30 minutes, RT), blocked (1 hour, RT; 5% normal donkey serum and 0.5% Triton-X100 in PBS (0.5% PBST) and then incubated with HuC/D antibody (ANNA1) diluted 1:50 in 1% normal donkey serum in 1x PBS either 2 hours (RT) or overnight (4°C). Tissue was then washed 3 times in 1x PBS (5 minutes/wash, RT) before incubating in secondary antibody diluted 1:400 in 5% normal donkey serum in 0.5% PBST (RT, 60–90 minutes), washing 3 times (RT, 5 minutes, 1× PBS), and mounting in 50% glycerol/50% 1× PBS on slides using coverslips. Incubation and wash steps were performed on a rocker.

### Quantification of enteric neurons in HuC/D stained colon

Whole-mount immunofluorescence images of the entire P19-21 age colon were processed using ImageJ (NIH) to add scale bars and generate the maximum intensity projections used to measure length of aganglionic colon, length of transition zone and region with normal appearing ENS anatomy. Transition zone was defined as region with lower than expected myenteric neuron density. Distal transition zone margin was defined by complete absence of the ENS or the presence of only single isolated enteric neurons.

### Hypoxia staining

To evaluate the degree of epithelial hypoxia, mice were treated with pimonidazole HCl (60 mg/kg, intraperitoneal) (Hypoxyprobe® Omni Kit, Hypoxyprobe, HP3-200Kit) one hour prior to euthanasia. Colon samples were fixed in 4% PFA for (24 hours, 4°C), equilibrated with 40% sucrose (24 hours, 4°C), frozen in Optimal Cutting Temperature (OCT) compound, and sectioned at 7 microns. Sections were probed according to kit instructions (1:50 rabbit anti-pimonidazole antibody) and stained with Cy-5 conjugated goat anti-rabbit antibody (Invitrogen). Samples were counterstained with DAPI.

### ATP measurements

Primary colonocytes from the most distal 1.5 cm of colon were isolated by opening colon longitudinally in 5 mL ice-cold Dulbecco’s Phosphate Buffered Saline (DPBS) in a 15 mL conical. After gently inverting once, colon was moved to a new tube of DPBS and swirled until clean of feces. Cleaned colon was then incubated in fresh DPBS with 0.03 M EDTA and 1.5 mM DTT on ice for 20 mins, before transferring to DPBS with 0.03 M EDTA for 10 mins at 37°C. Samples were shaken vigorously until colonocytes were released. Remaining tissues were removed and DPBS tubes containing colonocytes were centrifuges (800g, 5 min., 4°C). After supernatant was removed, colonocyte pellet was transferred to a cryovial and stored at-80°C. ATP measurement in colonocyte lysates was performed by using CellTiter-Glo® 3D Cell Viability Assay (Promega G9681), according to manufacturer’s instructions. ATP measurements were normalized to input protein quantified with Pierce™ BCA Protein Assay Kits (Thermo Scientific, 23225).

### Transmission electron microscopy (TEM)

1cm of P20 distal colon tissue from 3 mice of each diet and genotype (12 total) was collected and was fixed with 2.5% glutaraldehyde, 2% paraformaldehyde in 0.1 M sodium cacodylate buffer, pH 7.4 for 48h. Samples were submitted to Penn Electron Microscopy Resource Lab (RRID:SCR_022375) for processing, embedding, and staining. Mitochondria from TEM images taken at 40,000x magnification of distal colon epithelium were blinded, classified, and scored for damage (adapted from Matsuzawa-Ishimoto^166^) 6 images that spanned 3-4 distinct epithelial cells across different regions of the colon with at least 5 mitochondria per image were observed for each animal. Samples were scored for mitochondrial damage according to the rubric in **Supplemental Figure 1**. Mitochondria in each image were classified as type 1 mitochondria (dense with cristae structure maintained), type 2 mitochondria (mildly swollen, mild loss of density with a 50% decrease in the number of visible cristae), or type 3 mitochondria (severely aberrant morphology with disorganized cristae and over 70% of cristae are missing).

### Microscopy

*Confocal microscopy:* Whole-mount immunofluorescence images were acquired with a Zeiss LSM 710 confocal microscope or Zeiss LSM 980 with a 10x or 20x objective and Zeiss Zen 2.3 SP1 FP3 (Black) LSM710 Release Version 14.0.0.0.

*Light microscopy and fluorescent microscopy*: Images were acquired using a Nikon Ti2 epifluorescence microscope with 10x and 20x objectives and NIS-Elements Advanced Research software (Nikon). H&E-or Alcian blue–stained tissue images were acquired with the included attached camera. Fluorescent images were acquired with a Nikon Ti2 with an attached Hamamatsu C13440 Orca-Flash 4.0 Camera. Image brightness and contrast were uniformly adjusted in ImageJ, NIS-Elements AR, and Adobe Photoshop CS2.

### Administration of oral drugs

Mice were given a combination of neomycin (1 g/L, Research Products International N20040), ampicillin (1 g/L, Research Products International A40040), vancomycin (0.5 g/L, Mylan), metronidazole (0.5 g/L, Research Products International M81000), imipenem (0.5 g/L, Fresenius Kabi), and ciprofloxacin (0.2 g/L, Sigma-Aldrich 17850) in their drinking water from P20 to time of death or time of analysis. Control mice received water without antibiotics.

### Measuring colonic epithelial barrier permeability using FITC-dextran

On the day of the assay, 4 kDa fluorescein isothiocyanate (FITC)-dextran was dissolved in phosphate buffered saline (PBS) at 80 mg/mL. Blood was collected via submandibular (facial) vein from mice before gavaging with 150 μL FITC-dextran solution. 3 hours following gavage, blood was collected again from submandibular vein. Blood samples were centrifuged (1,000 x g, 12 min,4°C) to separate serum from cells. Fluorescence was quantified at an excitation wavelength of 485 nm and emission wavelength of 535 nm using Perkins-Elmer Envision fluorescent plate reader.

### Colon epithelial cell isolation for RNAseq

Mouse colon from distal cecum to rectum was placed in cold DPBS (Thermo Fisher, Cat#21-600-069). Protocol was adapted from Danan *et al.*^167^. Intraluminal contents were flushed with DPBS using a 19g needle attached to a 3 mL syringe. Colon was then opened longitudinally along the mesentery using ball tip dissecting scissors and mucus was removed from the epithelial lining by lightly scraping with a glass cover slip. After rinsing again in DPBS, distal colon was placed in epithelial dissociation buffer (1x Calcium/Magnesium free HBSS, 1mM N-acetylcysteine, 10mM EDTA) and vortexed for 30 seconds to release epithelial cells from submucosa. Proximal colon required an additional rinse in 1x Calcium/Magnesium free HBSS with 1mM N-acetyl cysteine for 1 minute (vortexing for 15 s, storing on ice 15 s, vortexing 15 s, storing on ice 15 s). Distal and proximal colon segments in epithelial dissociation buffer were then rotated (4°C, 1 hour) before scraping off epithelium with a glass cover slip into dissociation buffer and vortexing 30 seconds until crypts dissociated from each other to make a single cell suspension. The solution with crypts was then filtered through a 100uM cell strainer (VWR. Cat# 76327-102) into a 50mL conical tube. The filter was rinsed with 2mL of 4% FBS in HBSS. The crypt containing eluate that passed through the filter was centrifuged at 400 x g (5 min, 4°C). The supernatant was removed from the pellet and discarded. Pellet was resuspended in 500uL TRIzol® and stored at-80°C until RNA extraction.

### RNA extraction and bulk epithelial RNA sequencing

Total RNA was extracted from epithelial preparations using the Promega RNA Extraction Kit (AS1340) following the manufacturer’s instructions. RNA quality was assessed with the RNA ScreenTape Assay (Agilent). NanoDrop8000 was used to quantify RNA. Libraries were prepared (200ng RNA input) using Illumina TruSeq library prep and sequenced on a NovaSeq 6000 (v1.5 S2 200 cycle kit) with mean sequencing depth of 124.0 M clusters per sample. Reads were pseudoaligned and transcripts were quantified using *kallisto*^168^ (v0.44.0) in paired-end mode with mm10 as reference genome (http://igenomes.illumina.com.s3-website-us-east-1.amazonaws.com/Mus_musculus/UCSC/mm10/Mus_musculus_UCSC_mm10.tar.gz). On average, 59.3% of reads per sample were aligned. The R package *tximport* (v1.16.1)^169^ was used to export “lengthScaledTPM” gene counts. Genes with fewer than 200 counts across all 64 samples were omitted. Samples were separated by colon location for analyses. *DESeq2* (v4.4)^170^ was used to analyze differential gene expression as we compared groups of different diets (Protective versus Detrimental), different genotypes, and different colon regions (proximal versus distal). Reported adjusted p-values are FDR/BH-corrected p-values generated with Wald test, which are the default *DESeq2* parameters. Gene set enrichment analysis (GSEA) was performed using the R package *fgsea*^171, 172^. For GSEA with *fgsea*, the Molecular Signatures Database Mouse Collection “MH” (M: mouse, H: hallmark gene sets) were used to test the gene set enrichment of differentially expressed genes from DESeq2^173–175^. In R, fgsea was run using a minimum pathway size of 30, maximum pathway size of 500, and eps=0.0 as the boundary for estimating p-values, and p-values were adjusted in *fgsea* using the Benjamini-Hochberg procedure. Results from *fgsea* were filtered to keep gene sets with adjusted p-values less than 0.05. Gene Ontology analysis was performed using the online tool David Bioinformatics Resources (https://david.ncifcrf.gov/)^176^.

### qRT-PCR

Total RNA was extracted using TRIzol®. First-strand cDNA was synthesized using a High-Capacity cDNA Reverse Transcription Kit (Thermo Fisher Scientific, Cat# 4368814). Real-time PCR was performed using Fast SybrGreen Master Mix (Thermo Scientific, Cat#385612) and an QuantStudio 3 real-time PCR system. The relative amounts of transcripts were calculated using the 2^−ΔΔCt^ formula, normalized to *Gapdh* as an internal control. PCR primers are provided in Key Resources Table.

### Quantification and statistical analysis

Data are presented as mean ± SEM unless otherwise indicated. Statistical analyzes used GraphPad Prism 10 (GraphPad Software 10.1.0 (316)). To compare two groups, we used Student’s t-test when data were approximately normally distributed and of equal variance. When these criteria were not met, we used Mann-Whitney U tests to compare two groups. To compare more than two groups when used one-way ANOVA to evaluate effects of one independent variable on a dependent variable and two-way ANOVA when there were two independent variables. To analyze survival curves we employed log-rank tests. Figures were prepared with GraphPad Prism 9 or R v4.0.2. Statistical analyses are indicated in figure legends.

### Global untargeted metabolomic analysis performed by Metabolon (description of methods provided by Metabolon)

#### Sample Handling

Prior to shipping samples to Metabolon, food and stool were saved at-80°C. Following receipt by Metabolon, samples were inventoried and immediately stored at-80°C until processing.

#### Sample Preparation

Samples were prepared using the automated MicroLab STAR® system from Hamilton Company. Several recovery standards were added prior to the first step in the extraction process for QC purposes. To remove protein, dissociate small molecules bound to protein or trapped in the precipitated protein matrix, and to recover chemically diverse metabolites, proteins were precipitated with methanol under vigorous shaking for 2 min (Glen Mills GenoGrinder 2000) followed by centrifugation. The resulting extract was divided into multiple fractions: two for analysis by two separate reverse phase (RP)/UPLC-MS/MS methods with positive ion mode electrospray ionization (ESI), one for analysis by RP/UPLC-MS/MS with negative ion mode ESI, one for analysis by HILIC/UPLC-MS/MS with negative ion mode ESI, while the remaining fractions were reserved for backup. Samples were placed briefly on a TurboVap® (Zymark) to remove the organic solvent. The sample extracts were stored overnight under nitrogen before preparation for analysis.

#### QA/QC

Several types of controls were analyzed in concert with the experimental samples: a pooled matrix sample generated by taking a small volume of each experimental sample (or alternatively, use of a pool of well-characterized human plasma) served as a technical replicate throughout the data set; extracted water samples served as process blanks; and a cocktail of QC standards that were carefully chosen not to interfere with the measurement of endogenous compounds were spiked into every analyzed sample, allowed instrument performance monitoring and aided chromatographic alignment. Instrument variability was determined by calculating the median relative standard deviation (RSD) for the standards that were added to each sample prior to injection into the mass spectrometers. Overall process variability was determined by calculating the median RSD for all endogenous metabolites (*i.e.,* non-instrument standards) present in 100% of the pooled matrix samples. Experimental samples were randomized across the platform run with QC samples spaced evenly among the injections.

#### Ultrahigh Performance Liquid Chromatography-Tandem Mass Spectroscopy (UPLC-MS/MS)

All methods utilized a Waters ACQUITY ultra-performance liquid chromatography (UPLC) and a Thermo Scientific Q-Exactive high resolution/accurate mass spectrometer interfaced with a heated electrospray ionization (HESI-II) source and Orbitrap mass analyzer operated at 35,000 mass resolution (PMID: 32445384). The dried sample extracts were then reconstituted in solvents compatible to each of the four methods. Each reconstitution solvent contained a series of standards at fixed concentrations to ensure injection and chromatographic consistency. One aliquot was analyzed using acidic positive ion conditions, chromatographically optimized for more hydrophilic compounds (PosEarly). In this method, the extract was gradient eluted from a C18 column (Waters UPLC BEH C18-2.1×100 mm, 1.7 µm) using water and methanol, containing 0.05% perfluoropentanoic acid (PFPA) and 0.1% formic acid (FA). Another aliquot was also analyzed using acidic positive ion conditions, however it was chromatographically optimized for more hydrophobic compounds (PosLate). In this method, the extract was gradient eluted from the same aforementioned C18 column using methanol, acetonitrile, water, 0.05% PFPA and 0.01% FA and was operated at an overall higher organic content. Another aliquot was analyzed using basic negative ion optimized conditions using a separate dedicated C18 column (Neg). The basic extracts were gradient eluted from the column using methanol and water, however with 6.5mM Ammonium Bicarbonate at pH 8. The fourth aliquot was analyzed via negative ionization following elution from a HILIC column (Waters UPLC BEH Amide 2.1×150 mm, 1.7 µm) using a gradient consisting of water and acetonitrile with 10mM Ammonium Formate, pH 10.8 (HILIC). The MS analysis alternated between MS and data-dependent MS^n^ scans using dynamic exclusion. The scan range varied slightly between methods but covered 70-1000 m/z. Raw data files are archived and extracted as described below.

#### Bioinformatics

The informatics system consisted of four major components, the Laboratory Information Management System (LIMS), the data extraction and peak-identification software, data processing tools for QC and compound identification, and a collection of information interpretation and visualization tools for use by data analysts. The hardware and software foundations for these informatics components were the LAN backbone, and a database server running Oracle 10.2.0.1 Enterprise Edition.

#### LIMS

The purpose of the Metabolon LIMS system was to enable fully auditable laboratory automation through a secure, easy to use, and highly specialized system. The scope of the Metabolon LIMS system encompasses sample accessioning, sample preparation and instrumental analysis and reporting and advanced data analysis. All of the subsequent software systems are grounded in the LIMS data structures. It has been modified to leverage and interface with the in-house information extraction and data visualization systems, as well as third party instrumentation and data analysis software.

#### Data Extraction and Compound Identification

Raw data was extracted, peak-identified and QC processed using a combination of Metabolon developed software services (applications). Each of these services perform a specific task independently, and they communicate/coordinate with each other using industry-standard protocols. Compounds were identified by comparison to library entries of purified standards or recurrent unknown entities. Metabolon maintains a library based on authenticated standards that contains the retention time/index (RI), mass to charge ratio (*m/z)*, and fragmentation data on all molecules present in the library. Furthermore, biochemical identifications are based on three criteria: retention index within a narrow RI window of the proposed identification, accurate mass match to the library +/-10 ppm, and the MS/MS forward and reverse scores between the experimental data and authentic standards. The MS/MS scores are based on a comparison of the ions present in the experimental spectrum to the ions present in the library spectrum. While there may be similarities between molecules based on one of these factors, the use of all three data points is utilized to distinguish and differentiate biochemicals. More than 5,400 commercially available purified or in-house synthesized standard compounds have been acquired and analyzed on all platforms for determination of their analytical characteristics. An additional 7000 mass spectral entries have been created for structurally unnamed biochemicals, which have been identified by virtue of their recurrent nature (both chromatographic and mass spectral). These compounds have the potential to be identified by future acquisition of a matching purified standard or by classical structural analysis. Metabolon continuously adds biologically-relevant compounds to its chemical library to further enhance its level of Tier 1 metabolite identifications.

#### Compound Quality Control

A variety of curation procedures were carried out to ensure that a high-quality data set was made available for statistical analysis and data interpretation. The QC and curation processes were designed to ensure accurate and consistent identification of true chemical entities, and to remove or correct those representing system artifacts, mis-assignments, mis-integration and background noise. Metabolon data analysts use proprietary visualization and interpretation software to confirm the consistency of peak identification and integration among the various samples.

#### Metabolite Quantification and Data Normalization

Peaks were quantified using area-under-the-curve. For studies spanning multiple days, a data normalization step was performed to correct variation resulting from instrument inter-day tuning differences. Essentially, each compound was corrected in run-day blocks by registering the medians to equal one (1.00) and normalizing each data point proportionately (termed the “block correction”). For studies that did not require more than one day of analysis, no normalization is necessary, other than for purposes of data visualization. In certain instances, biochemical data may have been normalized to an additional factor (*e.g.,* cell counts, total protein as determined by Bradford assay, osmolality, etc.) to account for differences in metabolite levels due to differences in the amount of material present in each sample.

#### Statistical Calculations

Standard statistical analyses for metabolomics were performed in ArrayStudio/Jupyter Notebook on log transformed data. For those analyses not standard in ArrayStudio/Jupyter Notebook, the programs R (http://cran.r-project.org/) or JMP Statistical Software were used.

## Notes

### Competing Interest Statement

The authors have declared no competing interest.

## References

1. Heuckeroth RO. Hirschsprung disease. In: Christophe Faure NT, Carlo Di Lorenzo, ed. Pediatric Neurogastroenterology, Gastrointestinal Motility Disorders and Disorders of Gut Brain Interaction in Children. 3 ed. Switzerland: Springer Cham, 2023:355-370.

2. Heuckeroth RO. Hirschsprung disease - integrating basic science and clinical medicine to improve outcomes. Nat Rev Gastroenterol Hepatol 2018;15:152–167.

3. Mueller JL, Goldstein AM. The science of Hirschsprung disease: What we know and where we are headed. Semin Pediatr Surg 2022;31:151157.

4. Hei Ha JL, Hang Lui VC, Hang Tam PK. Embryology and anatomy of Hirschsprung disease. Semin Pediatr Surg 2022;31:151227.

5. Skinner MA. Hirschsprung’s disease. Curr Probl Surg 1996;33:389–460.

6. Swenson O, Rheinlander HF, Diamond I. Hirschsprung’s disease; a new concept of the etiology; operative results in 34 patients. N Engl J Med 1949;241:551–6.

7. Schneider S, Wright CM, Heuckeroth RO. Unexpected Roles for the Second Brain: Enteric Nervous System as Master Regulator of Bowel Function. Annu Rev Physiol 2019;81:235–259.

8. Furness JB. Integrated Neural and Endocrine Control of Gastrointestinal Function. Adv Exp Med Biol 2016;891:159–73.

9. Li S, Zhang Y, Li K, et al. Update on the Pathogenesis of the Hirschsprung-Associated Enterocolitis. Int J Mol Sci 2023;24.

10. Lewit RA, Kuruvilla KP, Fu M, et al. Current understanding of Hirschsprung-associated enterocolitis: Pathogenesis, diagnosis and treatment. Semin Pediatr Surg 2022;31:151162.

11. Swenson O, Bill AH, Jr. Resection of rectum and rectosigmoid with preservation of the sphincter for benign spastic lesions producing megacolon; an experimental study. Surgery 1948;24:212–20.

12. Zhang X, Sun D, Xu Q, et al. Risk factors for Hirschsprung disease-associated enterocolitis: a systematic review and meta-analysis. Int J Surg 2023;109:2509–2524.

13. Ahmad H, Levitt MA, Yacob D, et al. Evaluation and Management of Persistent Problems After Surgery for Hirschsprung Disease in a Child. Curr Gastroenterol Rep 2021;23:18.

14. Han JW, Youn JK, Oh C, et al. Why Do the Patients with Hirschsprung Disease Get Redo Pull-Through Operation? Eur J Pediatr Surg 2018.

15. Demehri FR, Halaweish IF, Coran AG, et al. Hirschsprung-associated enterocolitis: pathogenesis, treatment and prevention. Pediatr Surg Int 2013;29:873–81.

16. Coe A, Collins MH, Lawal T, et al. Reoperation for Hirschsprung disease: pathology of the resected problematic distal pull-through. Pediatr Dev Pathol 2012;15:30–8.

17. Neuman H, Forsythe P, Uzan A, et al. Antibiotics in early life: dysbiosis and the damage done. FEMS Microbiology Reviews 2018;42:489–499.

18. Pini Prato A, Rossi V, Avanzini S, et al. Hirschsprung’s disease: what about mortality? Pediatric Surgery International 2011;27:473–478.

19. Xie C, Yan J, Zhang Z, et al. Risk factors for Hirschsprung-associated enterocolitis following Soave: a retrospective study over a decade. BMC Pediatr 2022;22:654.

20. Chantakhow S, Khorana J, Tepmalai K, et al. Alterations of Gut Bacteria in Hirschsprung Disease and Hirschsprung-Associated Enterocolitis. Microorganisms 2021;9.

21. Tang W, Su Y, Yuan C, et al. Prospective study reveals a microbiome signature that predicts the occurrence of post-operative enterocolitis in Hirschsprung disease (HSCR) patients. Gut Microbes 2020:1–13.

22. Dariel A, Grynberg L, Auger M, et al. Analysis of enteric nervous system and intestinal epithelial barrier to predict complications in Hirschsprung’s disease. Sci Rep 2020;10:21725.

23. Neuvonen MI, Korpela K, Kyrklund K, et al. Intestinal Microbiota in Hirschsprung Disease. J Pediatr Gastroenterol Nutr 2018.

24. Gosain A, Frykman PK, Cowles RA, et al. Guidelines for the diagnosis and management of Hirschsprung-associated enterocolitis. Pediatr Surg Int 2017;33:517–521.

25. Keck S, Galati-Fournier V, Kym U, et al. Lack of Mucosal Cholinergic Innervation Is Associated With Increased Risk of Enterocolitis in Hirschsprung’s Disease. Cell Mol Gastroenterol Hepatol 2021;12:507–545.

26. Gosain A, Barlow-Anacker AJ, Erickson CS, et al. Impaired Cellular Immunity in the Murine Neural Crest Conditional Deletion of Endothelin Receptor-B Model of Hirschsprung’s Disease. PLoS One 2015;10:e0128822.

27. Frykman PK, Cheng Z, Wang X, et al. Enterocolitis causes profound lymphoid depletion in endothelin receptor B-and endothelin 3-null mouse models of Hirschsprung-associated enterocolitis. Eur J Immunol 2015;45:807–17.

28. Aboagye J, Goldstein SD, Salazar JH, et al. Age at presentation of common pediatric surgical conditions: Reexamining dogma. J Pediatr Surg 2014;49:995–9.

29. Nakagawa H, Miyata Y. Refractory Constipation in a 53-Year-Old Man. Gastroenterology 2021;161:429–430.

30. Pruitt LCC, Skarda DE, Rollins MD, et al. Hirschsprung-associated enterocolitis in children treated at US children’s hospitals. J Pediatr Surg 2020;55:535–540.

31. Tang CS, Li P, Lai FP, et al. Identification of Genes Associated with Hirschsprung Disease, Based on Whole-genome Sequence Analysis, and Potential Effects on Enteric Nervous System Development. Gastroenterology 2018.

32. Diposarosa R, Bustam NA, Sahiratmadja E, et al. Literature review: enteric nervous system development, genetic and epigenetic regulation in the etiology of Hirschsprung’s disease. Heliyon 2021;7:e07308.

33. Amiel J, Sproat-Emison E, Garcia-Barcelo M, et al. Hirschsprung disease, associated syndromes and genetics: a review. J Med Genet 2008;45:1–14.

34. Tilghman JM, Ling AY, Turner TN, et al. Molecular Genetic Anatomy and Risk Profile of Hirschsprung’s Disease. N Engl J Med 2019;380:1421–1432.

35. Moore SW, Johnson AG. Hirschsprung’s disease: genetic and functional associations of Down’s and Waardenburg syndromes. Semin Pediatr Surg 1998;7:156–61.

36. Quinn FM, Surana R, Puri P. The influence of trisomy 21 on outcome in children with Hirschsprung’s disease. J Pediatr Surg 1994;29:781–3.

37. Teitelbaum DH, Qualman SJ, Caniano DA. Hirschsprung’s disease. Identification of risk factors for enterocolitis. Ann Surg 1988;207:240–4.

38. Menezes M, Puri P. Long-term clinical outcome in patients with Hirschsprung’s disease and associated Down’s syndrome. J Pediatr Surg 2005;40:810–2.

39. Hom B, Boyd NK, Vogel BN, et al. Down Syndrome and Autoimmune Disease. Clin Rev Allergy Immunol 2024;66:261–273.

40. Zheng D, Liwinski T, Elinav E. Interaction between microbiota and immunity in health and disease. Cell Res 2020;30:492–506.

41. Salvi PS, Cowles RA. Butyrate and the Intestinal Epithelium: Modulation of Proliferation and Inflammation in Homeostasis and Disease. Cells 2021;10:1775.

42. Gehrig JL, Venkatesh S, Chang HW, et al. Effects of microbiota-directed foods in gnotobiotic animals and undernourished children. Science 2019;365.

43. Neustaeter A, Leibovitzh H, Turpin W, et al. Understanding Predictors of Crohn’s Disease: Determinants of Altered Barrier Function in Pre-Disease Phase of Crohn’s Disease. J Can Assoc Gastroenterol 2024;7:68–77.

44. Levine A, Wine E, Assa A, et al. Crohn’s Disease Exclusion Diet Plus Partial Enteral Nutrition Induces Sustained Remission in a Randomized Controlled Trial. Gastroenterology 2019;157:440–450 e8.

45. Telborn L, Tofft L, Kristensson Hallström I, et al. Diet plays a central role in parental self-treatment of children with Hirschsprung’s disease-a qualitative study. Acta Paediatr 2021;110:2610–2617.

46. Telborn L, Graneli C, Axelsson I, et al. Children with Hirschsprung’s Disease Report Dietary Effects on Gastrointestinal Complaints More Frequently than Controls. Children (Basel) 2023;10.

47. Telborn L, Kumlien C, Graneli C, et al. Diet and bowel function in children with Hirschsprung’s disease: development and content validation of a patient-reported questionnaire. BMC Nutr 2023;9:78.

48. Rogier EW, Frantz AL, Bruno ME, et al. Secretory antibodies in breast milk promote long-term intestinal homeostasis by regulating the gut microbiota and host gene expression. Proc Natl Acad Sci U S A 2014;111:3074–9.

49. Fichter M, Klotz M, Hirschberg DL, et al. Breast milk contains relevant neurotrophic factors and cytokines for enteric nervous system development. Mol Nutr Food Res 2011;55:1592–6.

50. Zhu Y, Zhang J, Zhang W, et al. Recent progress on health effects and biosynthesis of two key sialylated human milk oligosaccharides, 3’-sialyllactose and 6’-sialyllactose. Biotechnol Adv 2023;62:108058.

51. Kim YJ. Immunomodulatory Effects of Human Colostrum and Milk. Pediatr Gastroenterol Hepatol Nutr 2021;24:337–345.

52. Ali AS, Hasan SS, Kow CS, et al. Lactoferrin reduces the risk of respiratory tract infections: A meta-analysis of randomized controlled trials. Clin Nutr ESPEN 2021;45:26–32.

53. Lane PW. Association of megacolon with two recessive spotting genes in the mouse. J Hered 1966;57:29–31.

54. Hosoda K, Hammer RE, Richardson JA, et al. Targeted and Natural (Piebald-Lethal) Mutations of Endothelin-B receptor gene produce megacolon associated with spotted coat color in mice. Cell 1994;79:1267–1276.

55. Puffenberger EG, Hosoda K, Washington SS, et al. A missense mutation of the endothelin-B receptor gene in multigenic Hirschsprung’s disease. Cell 1994;79:1257–66.

56. Amiel J, Attie T, Jan D, et al. Heterozygous endothelin receptor B (EDNRB) mutations in isolated Hirschsprung disease. Hum Mol Genet 1996;5:355–7.

57. Gariepy CE, Cass DT, Yanagisawa M. Null mutation of endothelin receptor type B gene in spotting lethal rats causes aganglionic megacolon and white coat color. Proc Natl Acad Sci U S A 1996;93:867–72.

58. McCallion AS, Stames E, Conlon RA, et al. Phenotype variation in two-locus mouse models of Hirschsprung disease: tissue-specific interaction between Ret and Ednrb. Proc Natl Acad Sci U S A 2003;100:1826–31.

59. Ro S, Hwang SJ, Muto M, et al. Anatomic modifications in the enteric nervous system of piebald mice and physiological consequences to colonic motor activity. Am J Physiol Gastrointest Liver Physiol 2006;290:G710–8.

60. Soret R, Schneider S, Bernas G, et al. Glial Cell-Derived Neurotrophic Factor Induces Enteric Neurogenesis and Improves Colon Structure and Function in Mouse Models of Hirschsprung Disease. Gastroenterology 2020;159:1824–1838 e17.

61. Caniano DA, Teitelbaum DH, Qualman SJ, et al. The piebald-lethal murine strain: investigation of the cause of early death. J Pediatr Surg 1989;24:906–10.

62. Fujimoto T. Natural history and pathophysiology of enterocolitis in the piebald lethal mouse model of Hirschsprung’s disease. J Pediatr Surg 1988;23:237–42.

63. Fujimoto T, Reen DJ, Puri P. Inflammatory response in enterocolitis in the piebald lethal mouse model of Hirschsprung’s disease. Pediatr Res 1988;24:152–5.

64. Cheng Z, Dhall D, Zhao L, et al. Murine model of Hirschsprung-associated enterocolitis. I: phenotypic characterization with development of a histopathologic grading system. J Pediatr Surg 2010;45:475–82.

65. Goto Y, Uematsu S, Kiyono H. Epithelial glycosylation in gut homeostasis and inflammation. Nat Immunol 2016;17:1244–1251.

66. Hu Y, Mao K, Zeng Y, et al. Tripartite-motif protein 30 negatively regulates NLRP3 inflammasome activation by modulating reactive oxygen species production. J Immunol 2010;185:7699–705.

67. Rigottier-Gois L. Dysbiosis in inflammatory bowel diseases: the oxygen hypothesis. Isme j 2013;7:1256–61.

68. Litvak Y, Byndloss MX, Tsolis RM, et al. Dysbiotic Proteobacteria expansion: a microbial signature of epithelial dysfunction. Curr Opin Microbiol 2017;39:1–6.

69. Shelton CD, Byndloss MX. Gut Epithelial Metabolism as a Key Driver of Intestinal Dysbiosis Associated with Noncommunicable Diseases. Infect Immun 2020;88.

70. Velazquez EM, Nguyen H, Heasley KT, et al. Endogenous Enterobacteriaceae underlie variation in susceptibility to Salmonella infection. Nat Microbiol 2019;4:1057–1064.

71. Byndloss MX, Olsan EE, Rivera-Chávez F, et al. Microbiota-activated PPAR-γ signaling inhibits dysbiotic Enterobacteriaceae expansion. Science 2017;357:570–575.

72. Cevallos Stephanie A, Lee J-Y, Velazquez Eric M, et al. 5-Aminosalicylic Acid Ameliorates Colitis and Checks Dysbiotic Escherichia coli Expansion by Activating PPAR-γ Signaling in the Intestinal Epithelium. mBio 2021;12:10.1128/mbio.03227-20.

73. Lee JY, Cevallos SA, Byndloss MX, et al. High-Fat Diet and Antibiotics Cooperatively Impair Mitochondrial Bioenergetics to Trigger Dysbiosis that Exacerbates Pre-inflammatory Bowel Disease. Cell Host Microbe 2020;28:273–284.e6.

74. Zhu W, Winter MG, Byndloss MX, et al. Precision editing of the gut microbiota ameliorates colitis. Nature 2018;553:208–211.

75. Zhu W, Miyata N, Winter MG, et al. Editing of the gut microbiota reduces carcinogenesis in mouse models of colitis-associated colorectal cancer. J Exp Med 2019;216:2378–2393.

76. Lewit RA, Veras LV, Cowles RA, et al. Reducing Underdiagnosis of Hirschsprung-Associated Enterocolitis: A Novel Scoring System. J Surg Res 2021;261:253–260.

77. Obata S, Ieiri S, Akiyama T, et al. Nationwide survey of outcome in patients with extensive aganglionosis in Japan. Pediatr Surg Int 2019;35:547–550.

78. Melendez E, Goldstein AM, Sagar P, et al. Case records of the Massachusetts General Hospital. Case 3-2012. A newborn boy with vomiting, diarrhea, and abdominal distention. N Engl J Med 2012;366:361-72.

79. Kouranloo J, Sadeghian N, Monfared MK. Treatment and postoperative complication of 420 patients with congenital megacolon. Saudi Med J 2003;24 Suppl:S25-8.

80. Rescorla FJ, Morrison AM, Engles D, et al. Hirschsprung’s disease. Evaluation of mortality and long-term function in 260 cases. Arch Surg 1992;127:934–41; discussion 941-2.

81. Ralls MW, Freeman JJ, Rabah R, et al. Redo pullthrough for Hirschsprung disease: a single surgical group’s experience. J Pediatr Surg 2014;49:1394–9.

82. Levitt MA, Dickie B, Pena A. Evaluation and treatment of the patient with Hirschsprung disease who is not doing well after a pull-through procedure. Semin Pediatr Surg 2010;19:146–53.

83. Pini-Prato A, Mattioli G, Giunta C, et al. Redo surgery in Hirschsprung disease: what did we learn? Unicentric experience on 70 patients. J Pediatr Surg 2010;45:747-54.

84. Sutthatarn P, Lapidus-Krol E, Smith C, et al. Hirschsprung-associated inflammatory bowel disease: A multicenter study from the APSA Hirschsprung disease interest group. J Pediatr Surg 2023;58:856–861.

85. Rosen MJ, Dhawan A, Saeed SA. Inflammatory Bowel Disease in Children and Adolescents. JAMA Pediatr 2015;169:1053–60.

86. Unger CA, Hope MC, 3rd, Aladhami AK, et al. How stable is your vivarium’s temperature? Fluctuations in vivarium temperature significantly impact metabolism and behavior impeding scientific reproducibility. Physiol Behav 2023;258:114029.

87. Melendez-Fernandez OH, Walton JC, DeVries AC, et al. The role of daylight exposure on body mass in male mice. Physiol Behav 2023;266:114186.

88. Zierath DK, Davidson S, Manoukian J, et al. Diet composition and sterilization modifies intestinal microbiome diversity and burden of Theiler’s virus infection-induced acute seizures. bioRxiv 2023.

89. Armstrong JL, Saraf TS, Bhatavdekar O, et al. Spontaneous seizures in adult Fmr1 knockout mice: FVB.129P2-Pde6b(+)Tyr(c-ch)Fmr1(tm1Cgr)/J. Epilepsy Res 2022;182:106891.

90. Long LL, Svenson KL, Mourino AJ, et al. Shared and distinctive features of the gut microbiome of C57BL/6 mice from different vendors and production sites, and in response to a new vivarium. Lab Anim (NY) 2021;50:185–195.

91. Blacher E, Bashiardes S, Shapiro H, et al. Potential roles of gut microbiome and metabolites in modulating ALS in mice. Nature 2019;572:474–480.

92. Clayton ZS, McCurdy CE. Short-term thermoneutral housing alters glucose metabolism and markers of adipose tissue browning in response to a high-fat diet in lean mice. Am J Physiol Regul Integr Comp Physiol 2018;315:R627–R637.

93. Zeng MY, Inohara N, Nuñez G. Mechanisms of inflammation-driven bacterial dysbiosis in the gut. Mucosal Immunol 2017;10:18–26.

94. Lupp C, Robertson ML, Wickham ME, et al. Host-mediated inflammation disrupts the intestinal microbiota and promotes the overgrowth of Enterobacteriaceae. Cell Host Microbe 2007;2:204.

95. Li Y, Poroyko V, Yan Z, et al. Characterization of Intestinal Microbiomes of Hirschsprung’s Disease Patients with or without Enterocolitis Using Illumina-MiSeq High-Throughput Sequencing. PLoS One 2016;11:e0162079.

96. Frykman PK, Nordenskjold A, Kawaguchi A, et al. Characterization of Bacterial and Fungal Microbiome in Children with Hirschsprung Disease with and without a History of Enterocolitis: A Multicenter Study. PLoS One 2015;10:e0124172.

97. Recharla N, Geesala R, Shi XZ. Gut Microbial Metabolite Butyrate and Its Therapeutic Role in Inflammatory Bowel Disease: A Literature Review. Nutrients 2023;15.

98. Singh V, Lee G, Son H, et al. Butyrate producers, “The Sentinel of Gut”: Their intestinal significance with and beyond butyrate, and prospective use as microbial therapeutics. Front Microbiol 2022;13:1103836.

99. Grosheva I, Zheng D, Levy M, et al. High-Throughput Screen Identifies Host and Microbiota Regulators of Intestinal Barrier Function. Gastroenterology 2020.

100. Wang H, Wang Y, Zhao J, et al. Dietary Nondigestible Polysaccharides Ameliorate Colitis by Improving Gut Microbiota and CD4(+) Differentiation, as Well as Facilitating M2 Macrophage Polarization. JPEN J Parenter Enteral Nutr 2019;43:401–411.

101. Ye J, Lv L, Wu W, et al. Butyrate Protects Mice Against Methionine-Choline-Deficient Diet-Induced Non-alcoholic Steatohepatitis by Improving Gut Barrier Function, Attenuating Inflammation and Reducing Endotoxin Levels. Front Microbiol 2018;9:1967.

102. Machiels K, Joossens M, Sabino J, et al. A decrease of the butyrate-producing species Roseburia hominis and Faecalibacterium prausnitzii defines dysbiosis in patients with ulcerative colitis. Gut 2014;63:1275–83.

103. Liu C, Kuei C, Zhu J, et al. 3,5-Dihydroxybenzoic acid, a specific agonist for hydroxycarboxylic acid 1, inhibits lipolysis in adipocytes. J Pharmacol Exp Ther 2012;341:794–801.

104. Li X, Yao Z, Qian J, et al. Lactate Protects Intestinal Epithelial Barrier Function from Dextran Sulfate Sodium-Induced Damage by GPR81 Signaling. Nutrients 2024;16.

105. Xu HM, Zhao HL, Guo GJ, et al. Characterization of short-chain fatty acids in patients with ulcerative colitis: a meta-analysis. BMC Gastroenterol 2022;22:117.

106. Gasaly N, Hermoso MA, Gotteland M. Butyrate and the Fine-Tuning of Colonic Homeostasis: Implication for Inflammatory Bowel Diseases. Int J Mol Sci 2021;22.

107. Roediger WE. Utilization of nutrients by isolated epithelial cells of the rat colon. Gastroenterology 1982;83:424–9.

108. Steliou K, Boosalis MS, Perrine SP, et al. Butyrate histone deacetylase inhibitors. Biores Open Access 2012;1:192–8.

109. Zimmerman MA, Singh N, Martin PM, et al. Butyrate suppresses colonic inflammation through HDAC1-dependent Fas upregulation and Fas-mediated apoptosis of T cells. Am J Physiol Gastrointest Liver Physiol 2012;302:G1405–15.

110. Zhou D, Pan Q, Xin FZ, et al. Sodium butyrate attenuates high-fat diet-induced steatohepatitis in mice by improving gut microbiota and gastrointestinal barrier. World J Gastroenterol 2017;23:60–75.

111. Pedersen SS, Prause M, Williams K, et al. Butyrate inhibits IL-1beta-induced inflammatory gene expression by suppression of NF-kappaB activity in pancreatic beta cells. J Biol Chem 2022;298:102312.

112. Liu Y, Upadhyaya B, Fardin-Kia AR, et al. Dietary resistant starch type 4-derived butyrate attenuates nuclear factor-kappa-B1 through modulation of histone H3 trimethylation at lysine 27. Food Funct 2016;7:3772–3781.

113. Huang X, Oshima T, Tomita T, et al. Butyrate Alleviates Cytokine-Induced Barrier Dysfunction by Modifying Claudin-2 Levels. Biology (Basel) 2021;10.

114. Cobo ER, Kissoon-Singh V, Moreau F, et al. MUC2 Mucin and Butyrate Contribute to the Synthesis of the Antimicrobial Peptide Cathelicidin in Response to Entamoeba histolytica-and Dextran Sodium Sulfate-Induced Colitis. Infect Immun 2017;85.

115. Campbell Y, Fantacone ML, Gombart AF. Regulation of antimicrobial peptide gene expression by nutrients and by-products of microbial metabolism. Eur J Nutr 2012;51:899–907.

116. Nepelska M, Cultrone A, Beguet-Crespel F, et al. Butyrate produced by commensal bacteria potentiates phorbol esters induced AP-1 response in human intestinal epithelial cells. PLoS One 2012;7:e52869.

117. Zheng L, Kelly CJ, Battista KD, et al. Microbial-Derived Butyrate Promotes Epithelial Barrier Function through IL-10 Receptor-Dependent Repression of Claudin-2. J Immunol 2017;199:2976–2984.

118. Byndloss MX, Olsan EE, Rivera-Chavez F, et al. Microbiota-activated PPAR-gamma signaling inhibits dysbiotic Enterobacteriaceae expansion. Science 2017;357:570–575.

119. Kinoshita M, Suzuki Y, Saito Y. Butyrate reduces colonic paracellular permeability by enhancing PPARgamma activation. Biochem Biophys Res Commun 2002;293:827–31.

120. Wachtershauser A, Loitsch SM, Stein J. PPAR-gamma is selectively upregulated in Caco-2 cells by butyrate. Biochem Biophys Res Commun 2000;272:380–5.

121. Bachmann M, Meissner C, Pfeilschifter J, et al. Cooperation between the bacterial-derived short-chain fatty acid butyrate and interleukin-22 detected in human Caco2 colon epithelial/carcinoma cells. Biofactors 2017;43:283–292.

122. Yang W, Yu T, Huang X, et al. Intestinal microbiota-derived short-chain fatty acids regulation of immune cell IL-22 production and gut immunity. Nat Commun 2020;11:4457.

123. Furusawa Y, Obata Y, Fukuda S, et al. Commensal microbe-derived butyrate induces the differentiation of colonic regulatory T cells. Nature 2013;504:446–50.

124. Shao X, Liu L, Zhou Y, et al. High-fat diet promotes colitis-associated tumorigenesis by altering gut microbial butyrate metabolism. Int J Biol Sci 2023;19:5004–5019.

125. Chang PV, Hao L, Offermanns S, et al. The microbial metabolite butyrate regulates intestinal macrophage function via histone deacetylase inhibition. Proc Natl Acad Sci U S A 2014;111:2247–52.

126. Ji J, Shu D, Zheng M, et al. Microbial metabolite butyrate facilitates M2 macrophage polarization and function. Sci Rep 2016;6:24838.

127. Yin CL, Ma YJ. The Regulatory Mechanism of Hypoxia-inducible Factor 1 and its Clinical Significance. Curr Mol Pharmacol 2024;17:e18761429266116.

128. Karhausen J, Furuta GT, Tomaszewski JE, et al. Epithelial hypoxia-inducible factor-1 is protective in murine experimental colitis. J Clin Invest 2004;114:1098–106.

129. Wang T, Wang RX, Colgan SP. Physiologic hypoxia in the intestinal mucosa: a central role for short-chain fatty acids. Am J Physiol Cell Physiol 2024;327:C1087–C1093.

130. Byndloss MX, Pernitzsch SR, Baumler AJ. Healthy hosts rule within: ecological forces shaping the gut microbiota. Mucosal Immunol 2018.

131. Grosheva I, Zheng D, Levy M, et al. High-Throughput Screen Identifies Host and Microbiota Regulators of Intestinal Barrier Function. Gastroenterology 2020;159:1807–1823.

132. Nakamura A, Kurihara S, Takahashi D, et al. Symbiotic polyamine metabolism regulates epithelial proliferation and macrophage differentiation in the colon. Nat Commun 2021;12:2105.

133. Buckel W. Unusual enzymes involved in five pathways of glutamate fermentation. Appl Microbiol Biotechnol 2001;57:263–73.

134. Buckel W, Barker HA. Two pathways of glutamate fermentation by anaerobic bacteria. J Bacteriol 1974;117:1248–60.

135. Paparo L, Nocerino R, Ciaglia E, et al. Butyrate as a bioactive human milk protective component against food allergy. Allergy 2021;76:1398–1415.

136. Liao GQ, Han HL, Wang TC, et al. Comparative analysis of the fatty acid profiles in goat milk during different lactation periods and their interactions with volatile compounds and metabolites. Food Chem 2024;460:140427.

137. Pichler MJ, Yamada C, Shuoker B, et al. Butyrate producing colonic Clostridiales metabolise human milk oligosaccharides and cross feed on mucin via conserved pathways. Nat Commun 2020;11:3285.

138. Okumura R, Takeda K. Maintenance of intestinal homeostasis by mucosal barriers. Inflamm Regen 2018;38:5.

139. Pickert G, Neufert C, Leppkes M, et al. STAT3 links IL-22 signaling in intestinal epithelial cells to mucosal wound healing. J Exp Med 2009;206:1465–72.

140. Haque PS, Kapur N, Barrett TA, et al. Mitochondrial function and gastrointestinal diseases. Nat Rev Gastroenterol Hepatol 2024;21:537–555.

141. Zhou B, Te B, Wang L, et al. Combination of sodium butyrate and probiotics ameliorates severe burn-induced intestinal injury by inhibiting oxidative stress and inflammatory response. Burns 2022;48:1213–1220.

142. Blachier F, Beaumont M, Andriamihaja M, et al. Changes in the Luminal Environment of the Colonic Epithelial Cells and Physiopathological Consequences. Am J Pathol 2017;187:476–486.

143. Gosain A. Established and emerging concepts in Hirschsprung’s-associated enterocolitis. Pediatr Surg Int 2016.

144. Schneider KM, Blank N, Alvarez Y, et al. The enteric nervous system relays psychological stress to intestinal inflammation. Cell 2023;186:2823–2838 e20.

145. Wang H, Foong JPP, Harris NL, et al. Enteric neuroimmune interactions coordinate intestinal responses in health and disease. Mucosal Immunol 2022;15:27–39.

146. Schill EM, Floyd AN, Newberry RD. Neonatal development of intestinal neuroimmune interactions. Trends Neurosci 2022;45:928–941.

147. Schneider KM, Kim J, Bahnsen K, et al. Environmental perception and control of gastrointestinal immunity by the enteric nervous system. Trends Mol Med 2022;28:989–1005.

148. Avetisyan M, Wang H, Schill EM, et al. Hepatocyte Growth Factor and MET Support Mouse Enteric Nervous System Development, the Peristaltic Response, and Intestinal Epithelial Proliferation in Response to Injury. J Neurosci 2015;35:11543–58.

149. Hitch MC, Leinicke JA, Wakeman D, et al. Ret heterozygous mice have enhanced intestinal adaptation after massive small bowel resection. Am J Physiol Gastrointest Liver Physiol 2012;302:G1143–50.

150. Bjerknes M, Cheng H. Modulation of specific intestinal epithelial progenitors by enteric neurons. Proc Natl Acad Sci U S A 2001;98:12497–502.

151. Yildiz HM, Carlson TL, Goldstein AM, et al. Mucus Barriers to Microparticles and Microbes are Altered in Hirschsprung’s Disease. Macromol Biosci 2015;15:712–8.

152. Thiagarajah JR, Yildiz H, Carlson T, et al. Altered goblet cell differentiation and surface mucus properties in Hirschsprung disease. PLoS One 2014;9:e99944.

153. Aslam A, Spicer RD, Corfield AP. Histochemical and genetic analysis of colonic mucin glycoproteins in Hirschsprung’s disease. J Pediatr Surg 1999;34:330–3.

154. Aslam A, Spicer RD, Corfield AP. Turnover of radioactive mucin precursors in the colon of patients with Hirschsprung’s disease correlates with the development of enterocolitis. J Pediatr Surg 1998;33:103–5.

155. Gonzales J, Gulbransen BD. The Physiology of Enteric Glia. Annu Rev Physiol 2024.

156. Rao M, Gulbransen BD. Enteric Glia. Cold Spring Harb Perspect Biol 2024.

157. Croswell A, Amir E, Teggatz P, et al. Prolonged Impact of Antibiotics on Intestinal Microbial Ecology and Susceptibility to Enteric Salmonella Infection. Infection and Immunity 2009;77:2741–2753.

158. Schindelin J, Arganda-Carreras I, Frise E, et al. Fiji: an open-source platform for biological-image analysis. Nat Methods 2012;9:676-82.

159. Bolyen E, Rideout JR, Dillon MR, et al. Reproducible, interactive, scalable and extensible microbiome data science using QIIME 2. Nat Biotechnol 2019;37:852–857.

160. Hamady M, Walker JJ, Harris JK, et al. Error-correcting barcoded primers for pyrosequencing hundreds of samples in multiplex. Nat Methods 2008;5:235–7.

161. Bokulich NA, Subramanian S, Faith JJ, et al. Quality-filtering vastly improves diversity estimates from Illumina amplicon sequencing. Nature Methods 2013;10:57–59.

162. Amir A, McDonald D, Navas-Molina Jose A, et al. Deblur Rapidly Resolves Single-Nucleotide Community Sequence Patterns. mSystems 2017;2:10.1128/msystems.00191-16.

163. Robeson MS, II, O’Rourke DR, Kaehler BD, et al. RESCRIPt: Reproducible sequence taxonomy reference database management. PLOS Computational Biology 2021;17:e1009581.

164. Quast C, Pruesse E, Yilmaz P, et al. The SILVA ribosomal RNA gene database project: improved data processing and web-based tools. Nucleic Acids Res 2013;41:D590–6.

165. McMurdie PJ, Holmes S. phyloseq: An R Package for Reproducible Interactive Analysis and Graphics of Microbiome Census Data. PLOS ONE 2013;8:e61217.

166. Matsuzawa-Ishimoto Y, Shono Y, Gomez LE, et al. Autophagy protein ATG16L1 prevents necroptosis in the intestinal epithelium. Journal of Experimental Medicine 2017;214:3687–3705.

167. Danan CH, Naughton KE, Hayer KE, et al. Intestinal transit-amplifying cells require METTL3 for growth factor signaling and cell survival. JCI Insight 2023;8.

168. Bray NL, Pimentel H, Melsted P, et al. Near-optimal probabilistic RNA-seq quantification. Nature Biotechnology 2016;34:525–527.

169. Soneson C, Love MI, Robinson MD. Differential analyses for RNA-seq: transcript-level estimates improve gene-level inferences. F1000Res 2015;4:1521.

170. Love MI, Huber W, Anders S. Moderated estimation of fold change and dispersion for RNA-seq data with DESeq2. Genome Biol 2014;15:550.

171. Korotkevich G, Sukhov V, Budin N, et al. Fast gene set enrichment analysis. bioRxiv 2021:060012.

172. Subramanian A, Tamayo P, Mootha VK, et al. Gene set enrichment analysis: A knowledge-based approach for interpreting genome-wide expression profiles. Proceedings of the National Academy of Sciences 2005;102:15545–15550.

173. Liberzon A, Birger C, Thorvaldsdóttir H, et al. The Molecular Signatures Database (MSigDB) hallmark gene set collection. Cell Syst 2015;1:417–425.

174. Castanza AS, Recla JM, Eby D, et al. Extending support for mouse data in the Molecular Signatures Database (MSigDB). Nat Methods 2023;20:1619–1620.

175. Liberzon A, Subramanian A, Pinchback R, et al. Molecular signatures database (MSigDB) 3.0. Bioinformatics 2011;27:1739–1740.

176. Huang da W, Sherman BT, Lempicki RA. Systematic and integrative analysis of large gene lists using DAVID bioinformatics resources. Nat Protoc 2009;4:44–57.

