## Supplemental Figures and Tables for "Dietary manipulation of intestinal microbes prolongs survival in a mouse model of Hirschsprung disease"

**Supplemental Figures & Legends**

*Supplemental Figure 1: Mitochondria rubric used for mitochondrial damage and mitochondrial morphology scoring.*

A. Rubric for scoring mitochondrial damage adapted from Matsuzawa-Ishimoto 2017[166]. Score range is from 0-6 (normal to damaged) and consists of shape (0, 1, 2), cristae quality (0, 1, 2) and hue (0, 1, 2). 6 images that spanned 3-4 distinct epithelial cells across different regions of the colon with at least 5 mitochondria per image were scored for each animal. N=12 total (3 of each diet and genotype).
B. Mitochondria in each image were classified as type 1, 2 or 3 based on the following criterion:
    - Representative Type 1 mitochondria that appear dense with cristae structure maintained and long, sneaker shape.
    - Representative Type 2 mitochondria appear mildly swollen, with a mild loss of density with a 50% decrease in the number of visible cristae.
    - Representative Type 3 mitochondria have severely aberrant morphology with disorganized cristae and over 70% of cristae are missing.

### A. Mitochondria scoring criteria

| Criteria | **0** | **1** | **2** |
|---|---|---|---|
| **Shape** | Mix of oblong (sneaker) shape and rounded | Rounded without any oblong shape | Swollen, deformed, irregular shape |
| **Cristae** | Dense | Missing less than 75% cristae | Missing more than 75% cristae |
| **Hue** | Opaque | Semi-transparent | Transparent |

### B. Mitochondria morphology criteria

Type 1 examples:

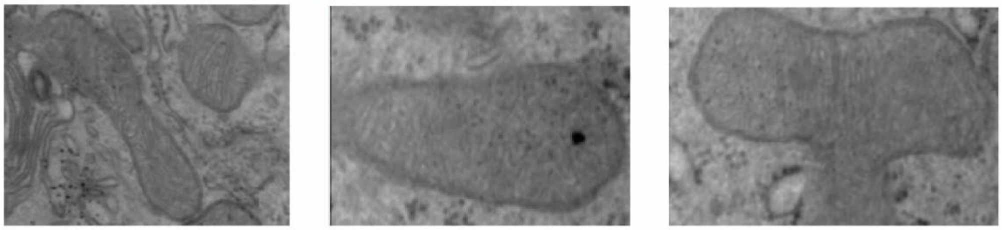

Type 2 examples:

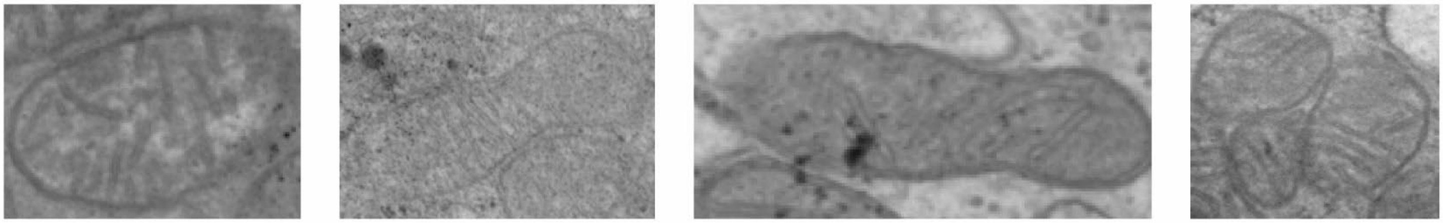

Type 3 examples:

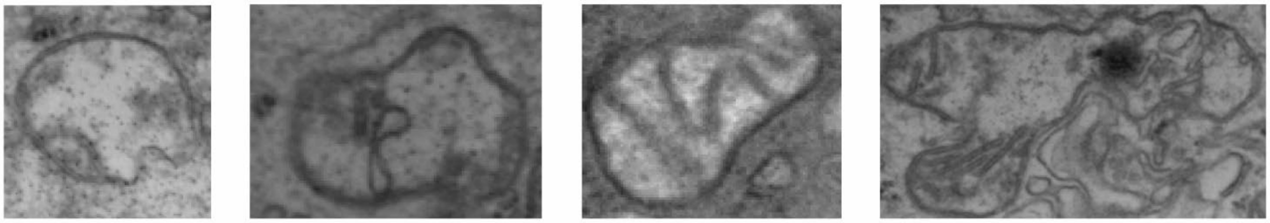

*Supplemental Table 1:* Metabolites more abundant (>4-fold difference, $p<0.05$) in Protective than in Detrimental diet food pellets

| **Metabolite** | **Abundance ratio in diets Protective/Detrimental** | **Charles-River 5075/Detrimental Diet Mean** | **LabDiet 5015/Protective Diet Mean** |
|---|---|---|---|
| glutamine | 82.65 | 0.03 | 2.29 |
| 3'-sialyllactose | 19.81 | 0.06 | 1.15 |
| 1-stearoyl-2-oleoyl-GPS (18:0/18:1) | 13.89 | 0.14 | 1.95 |
| cysteinylglycine | 13.4 | 0.32 | 4.22 |
| malonylgenistin | 12.86 | 0.14 | 1.86 |
| glucuronate | 12.84 | 0.18 | 2.33 |
| tricosanoyl sphingomyelin (d18:1/23:0)* | 12.25 | 0.16 | 1.94 |
| spermidine | 11.26 | 0.21 | 2.36 |
| flavone derivative C26H28O14 (2)* | 11.23 | 0.19 | 2.08 |
| lignoceroyl sphingomyelin (d18:1/24:0) | 10.75 | 0.18 | 1.92 |
| sphingomyelin (d18:0/20:0, d16:0/22:0)* | 9.86 | 0.18 | 1.8 |
| lanthionine | 9.59 | 0.12 | 1.13 |
| sphingomyelin (d18:1/17:0, d17:1/18:0, d19:1/16:0) | 9.19 | 0.12 | 1.06 |
| flavone derivative C26H28O14 (1)* | 8.76 | 0.22 | 1.97 |
| methylphosphate | 8.71 | 0.26 | 2.3 |
| ectoine | 8.55 | 0.24 | 2.08 |
| stearoyl ethanolamide | 8.45 | 0.21 | 1.8 |
| flavone derivative C26H28O14 (3)* | 8.21 | 0.24 | 1.97 |
| sphingomyelin (d18:1/21:0, d17:1/22:0, d16:1/23:0)* | 8.13 | 0.23 | 1.89 |
| 2-palmitoylglycerol (16:0) | 8.02 | 0.52 | 4.19 |
| piperine | 8 | 0.22 | 1.79 |
| benzoate | 7.87 | 0.27 | 2.12 |
| flavone derivative C26H28O14 (4)* | 7.84 | 0.25 | 1.94 |
| sphingomyelin (d18:1/20:0, d16:1/22:0)* | 7.74 | 0.23 | 1.77 |
| behenoyl sphingomyelin (d18:1/22:0)* | 7.66 | 0.23 | 1.75 |
| urea | 7.62 | 0.16 | 1.19 |
| enterolactone sulfate | 7.25 | 0.26 | 1.92 |
| propionylcarnitine (C3) | 6.66 | 0.27 | 1.77 |
| palmitoyl dihydrosphingomyelin (d18:0/16:0)* | 6.52 | 0.27 | 1.77 |
| arachidonoyl ethanolamide | 6.49 | 0.27 | 1.74 |
| sphingomyelin (d18:2/16:0, d18:1/16:1)* | 6.11 | 0.18 | 1.08 |
| 2-aminoadipate | 5.85 | 0.17 | 1.02 |
| tyrosol | 5.68 | 0.31 | 1.77 |
| 2,3-dihydroxyisovalerate | 5.66 | 0.42 | 2.35 |
| acetylcarnitine (C2) | 5.64 | 0.33 | 1.87 |
| propionylglycine | 5.49 | 0.21 | 1.16 |
| allantoic acid | 5.41 | 0.36 | 1.92 |
| salidroside | 5.29 | 0.34 | 1.8 |
| S-methylglutathione | 5.28 | 0.2 | 1.04 |
| isobutyrylcarnitine (C4) | 5.11 | 0.36 | 1.82 |

| | | | |
|---|---|---|---|
| palmitoyl sphingomyelin (d18:1/16:0) | 5.08 | 0.34 | 1.75 |
| sphingomyelin (d17:1/16:0, d18:1/15:0, d16:1/17:0)* | 5.06 | 0.34 | 1.72 |
| ethyl alpha-glucopyranoside | 5.05 | 0.48 | 2.43 |
| galactinol | 4.78 | 0.39 | 1.89 |
| flavin adenine dinucleotide (FAD) | 4.7 | 0.23 | 1.07 |
| trimethylamine N-oxide | 4.67 | 0.43 | 2.01 |
| phenyllactate (PLA) | 4.64 | 0.44 | 2.03 |
| 1-stearoyl-2-arachidonoyl-GPE (18:0/20:4) | 4.63 | 0.25 | 1.16 |
| glycosyl ceramide (d18:1/20:0, d16:1/22:0)* | 4.39 | 0.35 | 1.52 |
| 5-methylthioadenosine (MTA) | 4.38 | 0.42 | 1.83 |
| butyrylcarnitine (C4) | 4.33 | 0.37 | 1.61 |
| adenosine 3',5'-cyclic monophosphate (cAMP) | 4.31 | 0.4 | 1.7 |
| dihydroferulate | 4.27 | 0.46 | 1.96 |
| ophthalmate | 4.27 | 0.39 | 1.68 |
| UDP-N-acetylglucosamine/galactosamine | 4.16 | 0.42 | 1.75 |
| nicotianamine | 4.13 | 0.43 | 1.76 |
| N-acetyl-2-aminoadipate | 4.12 | 0.48 | 1.96 |
| ergothioneine | 4.1 | 0.47 | 1.91 |
| N,N,N-trimethyl-5-aminovalerate | 4.03 | 0.29 | 1.18 |

*Supplemental Table 2:* Metabolites more abundant (>4-fold difference, $p<0.05$) in Detrimental than in Protective diet food pellets

| Metabolite | Abundance ratio in diets<br><br>Detrimental/Protective | Charles-River 5075/Detrimental Diet Mean | LabDiet 5015/Protective Diet Mean |
|---|---|---|---|
| 1-eicosapentaenoyl-GPC (20:5)* | 426.93 | 1.33 | 0.003 |
| 1-docosahexaenoyl-GPC (22:6)* | 353.62 | 2.60 | 0.01 |
| 1-docosahexaenoyl-GPE (22:6)* | 146.12 | 1.17 | 0.01 |
| 5-(2-hydroxyethyl)-4-methylthiazole | 80.92 | 2.34 | 0.03 |
| docosahexaenoate (DHA; 22:6n3) | 78.44 | 2.14 | 0.03 |
| fructosyllysine | 67.76 | 2.21 | 0.03 |
| 1-pentadecanoyl-2-docosahexaenoyl-GPC (15:0/22:6)* | 65.69 | 1.06 | 0.02 |
| imidazole propionate | 65.31 | 2.26 | 0.03 |
| eicosapentaenoate (EPA; 20:5n3) | 51.96 | 2.16 | 0.04 |
| 1-stearoyl-2-docosahexaenoyl-GPC (18:0/22:6) | 47.66 | 1.16 | 0.02 |
| N6-carboxymethyllysine | 46.17 | 2.32 | 0.05 |
| 1-palmitoyl-2-docosahexaenoyl-GPC (16:0/22:6) | 43.95 | 1.18 | 0.03 |
| glycosyl ceramide (d16:1/24:1, d18:1/22:1)* | 41.64 | 1.11 | 0.03 |
| 5-HEPE | 33.74 | 1.12 | 0.03 |
| 1-erucoyl-GPC (22:1)* | 33.06 | 1.06 | 0.03 |
| taurocholate | 32.73 | 2.16 | 0.07 |
| 3-phosphoglycerate | 31.34 | 3.23 | 0.10 |
| trans-urocanate | 31.00 | 2.18 | 0.07 |
| 1-nervonoyl-GPC (24:1n9)* | 30.89 | 1.11 | 0.04 |
| pyrraline | 28.96 | 2.36 | 0.08 |

| | | | |
|---|---|---|---|
| cis-urocanate | 28.47 | 1.99 | 0.07 |
| hypoxanthine | 26.13 | 2.20 | 0.08 |
| cholate | 25.22 | 1.15 | 0.05 |
| N-methylhydantoin | 24.95 | 2.04 | 0.08 |
| taurine | 24.35 | 2.02 | 0.08 |
| 14-HDoHE/17-HDoHE | 23.87 | 2.33 | 0.10 |
| guanine | 22.37 | 2.38 | 0.11 |
| hydroxymethylpyrimidine | 21.93 | 2.07 | 0.09 |
| homocitrulline | 20.78 | 1.02 | 0.05 |
| hexadecatrienoate (16:3n3) | 19.03 | 2.18 | 0.11 |
| N2-acetyllysine | 18.76 | 1.09 | 0.06 |
| 1-palmitoyl-2-eicosapentaenoyl-GPC (16:0/20:5)* | 18.08 | 1.14 | 0.06 |
| sarcosine | 16.56 | 1.01 | 0.06 |
| vanillin | 16.02 | 2.00 | 0.12 |
| diosmin (diosmetin 7-rutinoside) | 15.83 | 3.51 | 0.22 |
| 1-docosahexaenoylglycerol (22:6) | 15.57 | 1.02 | 0.07 |
| stearidonate (18:4n3) | 15.29 | 1.97 | 0.13 |
| lysine | 14.85 | 1.93 | 0.13 |
| 1-methylguanidine | 14.52 | 2.07 | 0.14 |
| taurochenodeoxycholate | 13.59 | 1.88 | 0.14 |
| erucate (22:1n9) | 12.98 | 1.88 | 0.14 |
| harmane | 11.72 | 2.05 | 0.17 |
| (S)-a-amino-omega-caprolactam | 11.50 | 1.99 | 0.17 |
| thymine | 10.73 | 1.90 | 0.18 |
| palmitoloelycholine | 9.67 | 2.24 | 0.23 |
| citraconate/glutaconate | 9.63 | 2.07 | 0.21 |
| anserine | 9.30 | 1.73 | 0.19 |
| N6-carboxyethyllysine | 9.17 | 1.92 | 0.21 |
| maleate | 8.87 | 2.07 | 0.23 |
| linoleoyl-arachidonoyl-glycerol (18:2/20:4) [2]* | 8.36 | 1.89 | 0.23 |
| ceramide (d16:1/24:1, d18:1/22:1)* | 8.23 | 1.97 | 0.24 |
| ceramide (d18:1/14:0, d16:1/16:0)* | 8.06 | 1.86 | 0.23 |
| 4-methylthio-2-oxobutanoate | 7.85 | 1.94 | 0.25 |
| N-acetylhistidine | 7.83 | 1.87 | 0.24 |
| palmitoleoyl-linoleoyl-glycerol (16:1/18:2) [1]* | 7.28 | 1.97 | 0.27 |
| 3,4-dihydroxybutyrate | 6.94 | 1.94 | 0.28 |
| N-acetylmethionine sulfoxide | 6.74 | 1.98 | 0.29 |
| 2,6-dihydroxybenzoic acid | 6.57 | 1.84 | 0.28 |
| alpha-tocopherol acetate | 6.36 | 2.33 | 0.37 |
| 5-hydroxymethylfurfural | 6.34 | 1.54 | 0.24 |
| 5-hydroxylysine | 6.12 | 2.20 | 0.36 |
| N-formylmethionine | 5.97 | 1.88 | 0.31 |
| cyclo(gly-pro) | 5.47 | 1.85 | 0.34 |
| 3-formylindole | 5.39 | 1.93 | 0.36 |
| 1-palmityl-GPC (O-16:0) | 5.38 | 1.85 | 0.34 |
| mannose | 5.28 | 2.35 | 0.44 |

| | | | |
|---|---|---|---|
| 1-oleyl-GPC (O-18:1)* | 5.22 | 1.91 | 0.37 |
| 3-phenylpropionate (hydrocinnamate) | 5.22 | 1.79 | 0.34 |
| 1-palmitoleoyl-GPC (16:1)* | 5.14 | 1.99 | 0.39 |
| gamma-glutamyl-alpha-lysine | 5.13 | 1.75 | 0.34 |
| N-carbamoylalanine | 5.02 | 1.63 | 0.32 |
| N-acetylmethionine | 4.97 | 1.73 | 0.35 |
| oleoyl-linoleoyl-glycerol (18:1/18:2) [1] | 4.82 | 1.77 | 0.37 |
| feruloylquinate (2) | 4.55 | 1.79 | 0.39 |
| linoleoyl-linoleoyl-glycerol (18:2/18:2) [1]* | 4.51 | 1.69 | 0.37 |
| oxindolylalanine | 4.44 | 1.83 | 0.41 |
| 2-methylcitrate | 4.38 | 1.85 | 0.42 |
| thiamin (vitamin B1) | 4.37 | 1.75 | 0.40 |
| pantoate | 4.37 | 1.83 | 0.42 |
| coumaroylquinate (2) | 4.36 | 1.86 | 0.43 |
| 1-myristoyl-GPC (14:0) | 4.21 | 1.80 | 0.43 |
| linoleoyl-linolenoyl-glycerol (18:2/18:3) [1]* | 4.16 | 1.72 | 0.41 |
| hexadecadienoate (16:2n6) | 4.09 | 1.52 | 0.37 |
| N-acetylhistamine | 4.06 | 1.52 | 0.37 |

*Supplemental Table 3: Relative abundance of all metabolites in the Protective or Detrimental diets (chow)*

| Chemical name | Prot. food relative abundance (median) | Det. food relative abundance (median) | Prot. / Det. Fold-change |
|---|---|---|---|
| putrescine | 0.78 | 1.18 | 0.66 |
| spermidine | 1.98 | 0.21 | 9.46 |
| 1-methyladenine | 0.85 | 1.20 | 0.71 |
| 12,13-DiHOME | 0.92 | 1.00 | 0.92 |
| adenosine-2',3'-cyclic monophosphate | 0.52 | 1.51 | 0.34 |
| cytidine 2',3'-cyclic monophosphate | 0.46 | 1.15 | 0.40 |
| guanosine-2',3'-cyclic monophosphate | 0.43 | 1.55 | 0.28 |
| uridine-2',3'-cyclic monophosphate | 0.51 | 1.75 | 0.29 |
| 2-methylcitrate | 0.40 | 1.66 | 0.24 |
| alpha-ketoglutarate | 1.20 | 0.85 | 1.41 |
| kynurenate | 0.72 | 1.23 | 0.59 |
| uridine 3'-monophosphate (3'-UMP) | 0.48 | 1.30 | 0.37 |
| 3-hydroxyisobutyrate | 1.00 | 1.09 | 0.92 |
| 3-hydroxy-3-methylglutarate | 0.79 | 1.19 | 0.66 |
| 3-phosphoglycerate | 0.10 | 2.36 | 0.04 |
| cholate | 0.04 | 1.11 | 0.04 |
| 4-hydroxyphenylacetate | 1.31 | 0.68 | 1.93 |
| hypoxanthine | 0.09 | 2.14 | 0.04 |
| guanine | 0.10 | 2.49 | 0.04 |
| 9,10-DiHOME | 0.82 | 1.22 | 0.67 |
| linoleate (18:2n6) | 0.67 | 1.05 | 0.63 |
| laurate (12:0) | 1.42 | 0.69 | 2.04 |
| N6,N6,N6-trimethyllysine | 0.84 | 1.06 | 0.80 |
| N-acetylputrescine | 0.57 | 1.35 | 0.42 |
| N-formylmethionine | 0.32 | 1.85 | 0.17 |
| S-adenosylhomocysteine (SAH) | 1.22 | 0.84 | 1.46 |
| adenosine 3',5'-cyclic monophosphate (cAMP) | 1.74 | 0.42 | 4.17 |
| adenosine 5'-monophosphate (AMP) | 1.66 | 0.77 | 2.15 |
| 2'-deoxyadenosine | 1.18 | 1.00 | 1.18 |
| 5-methylthioadenosine (MTA) | 1.78 | 0.42 | 4.24 |
| N6-methyladenosine | 0.58 | 1.40 | 0.41 |
| allantoic acid | 1.92 | 0.36 | 5.30 |
| arachidonate (20:4n6) | 1.81 | 1.00 | 1.81 |
| arginine | 1.32 | 0.74 | 1.78 |
| argininosuccinate | 1.00 | 0.91 | 1.10 |
| ascorbate (vitamin C) | 0.98 | 1.00 | 0.98 |
| aspartate | 1.21 | 0.88 | 1.38 |
| 3-(4-hydroxyphenyl)lactate | 1.15 | 0.88 | 1.30 |
| phenylpyruvate | 0.96 | 1.00 | 0.96 |

| | | | |
|---|---|---|---|
| beta-alanine | 0.81 | 1.16 | 0.70 |
| carnosine | 1.29 | 0.98 | 1.32 |
| biotin | 0.87 | 1.11 | 0.79 |
| succinate | 0.85 | 1.13 | 0.75 |
| 3-hydroxybutyrate (BHBA) | 1.18 | 0.78 | 1.51 |
| cholesterol | 0.87 | 1.06 | 0.82 |
| choline phosphate | 1.38 | 0.68 | 2.03 |
| creatinine | 0.48 | 1.76 | 0.27 |
| cysteinylglycine | 4.18 | 0.29 | 14.35 |
| cytidine 5'-monophosphate (5'-CMP) | 1.30 | 0.63 | 2.08 |
| glucose 6-phosphate | 1.00 | 0.99 | 1.02 |
| sphingosine | 0.85 | 1.00 | 0.85 |
| dihydroxyacetone phosphate (DHAP) | 1.09 | 0.37 | 2.95 |
| cystathionine | 1.35 | 0.93 | 1.46 |
| sphinganine | 1.72 | 0.56 | 3.05 |
| behenate (22:0)* | 0.80 | 1.00 | 0.80 |
| flavin adenine dinucleotide (FAD) | 1.05 | 0.22 | 4.70 |
| folate | 0.39 | 1.72 | 0.23 |
| fumarate | 1.06 | 0.97 | 1.09 |
| gamma-glutamylglutamate | 1.24 | 0.84 | 1.47 |
| gluconate | 0.74 | 1.14 | 0.65 |
| glutarate (C5-DC) | 0.69 | 2.16 | 0.32 |
| glycine | 0.85 | 1.14 | 0.74 |
| guanidinoacetate | 0.57 | 1.48 | 0.39 |
| guanosine 5'-monophosphate (5'-GMP) | 1.18 | 0.96 | 1.23 |
| 2'-deoxyguanosine | 0.95 | 1.00 | 0.95 |
| histamine | 0.90 | 1.10 | 0.82 |
| histidine | 1.00 | 1.00 | 1.00 |
| hypotaurine | 1.00 | 0.61 | 1.64 |
| inosine | 0.39 | 1.55 | 0.25 |
| myo-inositol | 1.00 | 1.03 | 0.97 |
| inositol 1-phosphate (I1P) | 0.68 | 1.07 | 0.63 |
| isoleucine | 1.45 | 0.68 | 2.13 |
| 2-aminoadipate | 1.02 | 0.17 | 5.82 |
| 3-sulfo-alanine | 0.62 | 1.33 | 0.47 |
| citrulline | 1.33 | 0.74 | 1.80 |
| saccharopine | 1.55 | 0.60 | 2.59 |
| lactose | 1.83 | 0.46 | 4.00 |
| leucine | 1.11 | 0.98 | 1.14 |
| lithocholate | 1.00 | 0.62 | 1.60 |
| lysine | 0.13 | 1.93 | 0.07 |
| malate | 0.93 | 1.05 | 0.89 |
| methionine | 1.72 | 0.56 | 3.05 |

| | | | |
|---|---|---|---|
| palmitate (16:0) | 0.76 | 1.02 | 0.75 |
| nicotinamide adenine dinucleotide reduced (NADH) | 1.00 | 0.90 | 1.11 |
| nicotinamide | 0.90 | 1.01 | 0.89 |
| ornithine | 0.73 | 1.07 | 0.68 |
| orotate | 1.21 | 0.68 | 1.78 |
| glutathione, oxidized (GSSG) | 1.00 | 0.59 | 1.71 |
| palmitoleate (16:1n7) | 0.66 | 1.04 | 0.63 |
| phenylalanine | 0.97 | 1.00 | 0.97 |
| phosphate | 0.45 | 1.43 | 0.32 |
| phytosphingosine | 0.95 | 1.29 | 0.74 |
| proline | 0.81 | 1.02 | 0.79 |
| lactate | 1.37 | 0.53 | 2.58 |
| pyridoxal | 1.07 | 0.33 | 3.21 |
| glutathione, reduced (GSH) | 1.00 | 0.14 | 6.98 |
| retinol (vitamin A) | 0.86 | 1.89 | 0.45 |
| riboflavin (vitamin B2) | 0.56 | 2.33 | 0.24 |
| salicylate | 0.72 | 1.40 | 0.51 |
| serine | 1.03 | 0.98 | 1.05 |
| serotonin | 1.00 | 1.02 | 0.99 |
| taurine | 0.06 | 2.02 | 0.03 |
| myristate (14:0) | 1.11 | 1.00 | 1.11 |
| urea | 1.23 | 0.15 | 8.42 |
| uridine | 1.04 | 0.95 | 1.09 |
| 2'-deoxyuridine | 0.80 | 1.05 | 0.76 |
| trans-urocanate | 0.07 | 2.21 | 0.03 |
| glutamate | 1.24 | 0.75 | 1.64 |
| glutamine | 2.18 | 0.03 | 80.30 |
| threonine | 0.43 | 1.68 | 0.26 |
| tryptophan | 0.87 | 1.13 | 0.77 |
| valine | 0.43 | 1.06 | 0.41 |
| nicotinate | 0.15 | 1.44 | 0.11 |
| pyridoxamine | 1.31 | 0.77 | 1.70 |
| glucose | 0.77 | 1.28 | 0.60 |
| adenosine | 0.94 | 1.04 | 0.91 |
| betaine | 0.82 | 1.20 | 0.68 |
| cysteine | 1.00 | 0.35 | 2.89 |
| mannose | 0.37 | 2.26 | 0.16 |
| dimethylglycine | 0.91 | 1.19 | 0.77 |
| alanine | 0.95 | 1.04 | 0.92 |
| tyrosine | 0.76 | 1.38 | 0.55 |
| malonate | 0.98 | 1.03 | 0.95 |
| pseudouridine | 1.07 | 0.94 | 1.13 |
| pyruvate | 1.07 | 0.81 | 1.32 |

| | | | |
|---|---|---|---|
| uracil | 0.65 | 1.18 | 0.56 |
| cytidine | 0.95 | 1.11 | 0.85 |
| uridine 5'-monophosphate (UMP) | 1.20 | 0.86 | 1.39 |
| thymidine | 0.70 | 1.22 | 0.57 |
| thiamin (vitamin B1) | 0.39 | 1.72 | 0.23 |
| fructose | 0.40 | 1.42 | 0.28 |
| raffinose | 1.25 | 0.76 | 1.65 |
| adenine | 0.57 | 1.44 | 0.39 |
| cytosine | 0.83 | 1.10 | 0.76 |
| thymine | 0.18 | 1.89 | 0.10 |
| caprate (10:0) | 1.36 | 0.79 | 1.72 |
| margarate (17:0) | 1.04 | 1.00 | 1.04 |
| nonadecanoate (19:0) | 0.72 | 1.00 | 0.72 |
| arachidate (20:0) | 0.88 | 1.00 | 0.88 |
| maltose | 1.14 | 0.65 | 1.75 |
| asparagine | 1.00 | 1.01 | 0.99 |
| dihydroorotate | 1.61 | 0.39 | 4.09 |
| heptanoate (7:0) | 0.97 | 0.56 | 1.74 |
| caproate (6:0) | 1.44 | 0.80 | 1.80 |
| stachyose | 1.21 | 0.62 | 1.96 |
| sucrose | 1.14 | 0.80 | 1.42 |
| pyridoxine (vitamin B6) | 0.52 | 1.45 | 0.36 |
| trans-4-hydroxyproline | 0.87 | 1.14 | 0.76 |
| allantoin | 0.81 | 1.05 | 0.77 |
| galactitol (dulcitol) | 0.92 | 1.00 | 0.92 |
| xanthine | 0.75 | 1.05 | 0.72 |
| glucarate (saccharate) | 1.39 | 0.55 | 2.55 |
| 5-oxoproline | 0.58 | 1.45 | 0.40 |
| picolinate | 0.43 | 1.43 | 0.30 |
| sarcosine | 0.06 | 1.02 | 0.06 |
| pantothenate | 0.83 | 1.07 | 0.77 |
| pipecolate | 0.80 | 1.12 | 0.72 |
| phosphoethanolamine | 1.21 | 0.31 | 3.96 |
| gamma-glutamylcysteine | 1.00 | 0.19 | 5.16 |
| glycerate | 0.59 | 1.46 | 0.41 |
| 3-ureidopropionate | 0.99 | 1.16 | 0.85 |
| N-acetylleucine | 0.95 | 1.05 | 0.90 |
| N-acetylmethionine | 0.33 | 1.75 | 0.19 |
| N-acetylvaline | 0.83 | 1.23 | 0.67 |
| erucate (22:1n9) | 0.09 | 1.89 | 0.05 |
| tryptamine | 1.49 | 0.68 | 2.18 |
| guanosine | 0.77 | 1.19 | 0.64 |
| gamma-glutamyltyrosine | 1.62 | 0.65 | 2.48 |

| | | | |
|---|---|---|---|
| alpha-tocopherol | 0.90 | 1.10 | 0.82 |
| N-carbamoylaspartate | 0.90 | 1.04 | 0.86 |
| N-acetylalanine | 0.71 | 1.16 | 0.61 |
| 4-acetamidobutanoate | 0.62 | 1.41 | 0.44 |
| 3-aminoisobutyrate | 0.62 | 1.67 | 0.37 |
| chenodeoxycholate | 0.94 | 1.15 | 0.82 |
| citrate | 1.17 | 0.84 | 1.40 |
| tyramine | 1.26 | 0.87 | 1.45 |
| urate | 0.92 | 1.11 | 0.83 |
| oleoyl ethanolamide | 1.32 | 0.64 | 2.07 |
| gamma-glutamylglutamine | 1.24 | 0.69 | 1.81 |
| 4-hydroxyphenylpyruvate | 1.12 | 0.81 | 1.37 |
| N-acetylneuraminate | 0.97 | 1.15 | 0.85 |
| isocitrate | 1.18 | 0.65 | 1.83 |
| phenethylamine | 0.92 | 1.07 | 0.87 |
| N-acetyl-glucosamine 1-phosphate | 1.50 | 0.44 | 3.40 |
| N-acetylglucosaminylasparagine | 0.89 | 1.03 | 0.87 |
| cytidine 5'-diphosphocholine | 1.80 | 0.63 | 2.85 |
| creatine | 1.18 | 0.82 | 1.43 |
| dihomo-linoleate (20:2n6) | 2.13 | 1.00 | 2.13 |
| gamma-glutamylhistidine | 1.34 | 0.89 | 1.50 |
| 2-hydroxystearate | 1.08 | 1.00 | 1.08 |
| N1-methyladenosine | 0.91 | 0.81 | 1.12 |
| glycerol | 0.63 | 1.30 | 0.49 |
| choline | 1.00 | 0.99 | 1.00 |
| anthranilate | 0.75 | 1.25 | 0.60 |
| gamma-glutamylleucine | 1.00 | 1.00 | 1.00 |
| p-hydroxybenzaldehyde | 0.41 | 1.53 | 0.27 |
| nicotinamide adenine dinucleotide (NAD+) | 1.00 | 0.83 | 1.21 |
| 3-methoxytyrosine | 0.85 | 1.05 | 0.81 |
| beta-hydroxyisovalerate | 1.02 | 0.98 | 1.04 |
| arachidonoyl ethanolamide | 1.77 | 0.24 | 7.30 |
| palmitoyl ethanolamide | 1.30 | 0.63 | 2.06 |
| linoleamide (18:2n6) | 3.14 | 0.82 | 3.82 |
| N-linoleoylglycine | 1.57 | 1.00 | 1.57 |
| N-palmitoyl-sphingosine (d18:1/16:0) | 1.22 | 0.73 | 1.67 |
| 1-palmitoyl-2-oleoyl-GPE (16:0/18:1) | 1.32 | 0.73 | 1.80 |
| 1-palmitoyl-2-linoleoyl-GPI (16:0/18:2) | 1.17 | 0.86 | 1.36 |
| 1-palmitoyl-2-linoleoyl-GPC (16:0/18:2) | 1.15 | 0.87 | 1.33 |
| stearoyl sphingomyelin (d18:1/18:0) | 1.73 | 0.49 | 3.57 |
| 1-palmitoyl-2-oleoyl-GPC (16:0/18:1) | 0.97 | 1.03 | 0.94 |
| 1-palmitoyl-2-linoleoyl-GPG (16:0/18:2) | 1.39 | 0.70 | 1.97 |
| N-stearoyl-sphingosine (d18:1/18:0)* | 1.35 | 0.65 | 2.09 |

| | | | |
|---|---|---|---|
| glycochenodeoxycholate | 1.00 | 0.38 | 2.62 |
| taurochenodeoxycholate | 0.10 | 1.88 | 0.05 |
| taurocholate | 0.05 | 2.16 | 0.02 |
| acetylcholine | 0.77 | 1.20 | 0.64 |
| 2-hydroxyhippurate (salicylurate) | 1.56 | 0.60 | 2.61 |
| azelate (C9-DC) | 0.44 | 1.73 | 0.25 |
| eicosapentaenoate (EPA; 20:5n3) | 0.04 | 2.19 | 0.02 |
| methylsuccinate | 0.76 | 1.17 | 0.65 |
| tricarballylate | 0.56 | 1.39 | 0.40 |
| ethylmalonate | 0.62 | 1.56 | 0.40 |
| carnitine | 1.21 | 0.82 | 1.46 |
| benzoate | 2.22 | 0.26 | 8.58 |
| 3-phenylpropionate (hydrocinnamate) | 0.35 | 1.81 | 0.19 |
| hippurate | 1.46 | 0.58 | 2.54 |
| xanthurenate | 0.89 | 1.02 | 0.88 |
| suberate (C8-DC) | 0.62 | 1.56 | 0.40 |
| 3-methyl-2-oxovalerate | 1.31 | 0.75 | 1.74 |
| methionine sulfoxide | 0.54 | 1.50 | 0.36 |
| 3-methylhistidine | 0.92 | 1.08 | 0.85 |
| anserine | 0.12 | 1.67 | 0.07 |
| 5-hydroxylysine | 0.38 | 2.33 | 0.16 |
| 2-ketogulonate | 1.31 | 0.62 | 2.11 |
| 2-phosphoglycerate | 0.82 | 1.00 | 0.82 |
| 4-guanidinobutanoate | 0.94 | 1.03 | 0.92 |
| 5-(2-hydroxyethyl)-4-methylthiazole | 0.03 | 2.45 | 0.01 |
| agmatine | 0.91 | 1.05 | 0.87 |
| cadaverine | 0.62 | 1.25 | 0.50 |
| 2'-deoxycytidine | 0.69 | 1.43 | 0.48 |
| flavin mononucleotide (FMN) | 1.20 | 0.82 | 1.47 |
| glucuronate | 1.97 | 0.15 | 12.87 |
| glycerol 3-phosphate | 0.84 | 1.24 | 0.68 |
| imidazole lactate | 1.18 | 0.73 | 1.62 |
| kynurenine | 0.55 | 1.37 | 0.40 |
| glycerophosphorylcholine (GPC) | 1.01 | 0.92 | 1.10 |
| allo-threonine | 0.83 | 0.65 | 1.27 |
| N-acetylglutamate | 1.04 | 1.00 | 1.04 |
| tartarate | 0.67 | 1.20 | 0.56 |
| xanthosine | 1.15 | 0.77 | 1.50 |
| galactose 1-phosphate | 1.53 | 0.39 | 3.98 |
| ribitol | 0.92 | 1.04 | 0.88 |
| 2-isopropylmalate | 0.91 | 1.05 | 0.87 |
| quinate | 0.59 | 1.25 | 0.47 |
| gentisate | 0.55 | 1.70 | 0.32 |

| | | | |
|---|---|---|---|
| 5-aminovalerate | 0.94 | 1.14 | 0.83 |
| indolelactate | 0.54 | 1.39 | 0.39 |
| 3-indoxyl sulfate | 1.00 | 0.89 | 1.13 |
| glycylvaline | 1.00 | 0.99 | 1.01 |
| gamma-glutamylphenylalanine | 1.84 | 0.62 | 2.98 |
| 4-methyl-2-oxopentanoate | 1.62 | 0.63 | 2.59 |
| 1,5-anhydroglucitol (1,5-AG) | 1.16 | 0.84 | 1.38 |
| 1-palmityl-GPC (O-16:0) | 0.31 | 1.88 | 0.16 |
| 1-stearoyl-2-arachidonoyl-GPI (18:0/20:4) | 0.95 | 1.10 | 0.86 |
| 1-stearoyl-2-oleoyl-GPS (18:0/18:1) | 1.91 | 0.12 | 15.54 |
| 1-palmitoyl-2-oleoyl-GPG (16:0/18:1) | 1.20 | 0.93 | 1.29 |
| 1,2-dipalmitoyl-GPG (16:0/16:0) | 1.28 | 0.85 | 1.51 |
| 1-stearoyl-GPI (18:0) | 0.68 | 1.00 | 0.68 |
| 1,2-dipalmitoyl-GPC (16:0/16:0) | 1.14 | 0.85 | 1.34 |
| docosahexaenoate (DHA; 22:6n3) | 0.02 | 2.16 | 0.01 |
| 1-myristoyl-2-palmitoyl-GPC (14:0/16:0) | 1.52 | 0.51 | 2.99 |
| alpha-hydroxyisocaproate | 1.16 | 0.80 | 1.45 |
| maleate | 0.24 | 2.04 | 0.12 |
| 4-acetylphenol sulfate | 1.02 | 0.61 | 1.67 |
| 1-methylguanidine | 0.14 | 2.03 | 0.07 |
| 2-hydroxyoctanoate | 1.09 | 0.92 | 1.18 |
| levulinate (4-oxovalerate) | 0.42 | 1.01 | 0.41 |
| mevalonolactone | 1.04 | 0.71 | 1.47 |
| 3-hydroxyoctanoate | 1.00 | 1.10 | 0.91 |
| phenyllactate (PLA) | 1.73 | 0.42 | 4.15 |
| palmitoylcarnitine (C16) | 0.89 | 1.00 | 0.89 |
| hyodeoxycholate | 1.00 | 0.83 | 1.20 |
| N-acetylaspartate (NAA) | 0.53 | 1.55 | 0.34 |
| harmane | 0.18 | 2.11 | 0.08 |
| acetylcarnitine (C2) | 1.93 | 0.34 | 5.75 |
| cysteine s-sulfate | 1.00 | 0.42 | 2.35 |
| lactobionate | 1.85 | 0.52 | 3.56 |
| 1-palmitoylglycerol (16:0) | 4.04 | 0.67 | 6.05 |
| oxalate (ethanedioate) | 0.69 | 1.37 | 0.50 |
| erythritol | 0.70 | 1.02 | 0.68 |
| 4-hydroxybenzoate | 0.87 | 1.14 | 0.76 |
| adipate (C6-DC) | 0.82 | 1.03 | 0.80 |
| sinapate | 0.54 | 1.84 | 0.29 |
| galacturonate | 1.18 | 0.70 | 1.69 |
| galactarate (mucic acid) | 1.47 | 0.63 | 2.35 |
| 3-hydroxypyridine | 0.81 | 1.20 | 0.68 |
| galactinol | 1.93 | 0.38 | 5.05 |
| biochanin A | 0.65 | 1.30 | 0.50 |

| | | | |
|---|---|---|---|
| naringenin | 1.04 | 0.85 | 1.23 |
| 1-oleoylglycerol (18:1) | 1.07 | 1.00 | 1.07 |
| 1-stearoylglycerol (18:0) | 1.00 | 0.90 | 1.11 |
| apigenin | 0.88 | 1.03 | 0.86 |
| 3-methyl-2-oxobutyrate | 1.05 | 0.32 | 3.24 |
| 2-oleoylglycerol (18:1) | 0.95 | 1.00 | 0.95 |
| 8-hydroxyoctanoate | 0.53 | 1.92 | 0.28 |
| homocitrulline | 0.05 | 1.00 | 0.05 |
| alpha-glutamylglutamate | 1.00 | 0.72 | 1.38 |
| glycylproline | 1.07 | 0.98 | 1.09 |
| 1-kestose | 0.97 | 1.00 | 0.97 |
| 2-linoleoylglycerol (18:2) | 0.63 | 1.12 | 0.56 |
| N-acetylglycine | 1.39 | 0.69 | 2.00 |
| ribonate | 0.95 | 1.04 | 0.92 |
| threonate | 0.86 | 1.19 | 0.72 |
| galactonate | 0.86 | 1.10 | 0.78 |
| glycerol 2-phosphate | 0.54 | 1.20 | 0.45 |
| beta-sitosterol | 0.82 | 1.27 | 0.64 |
| indoleacetate | 0.75 | 1.19 | 0.63 |
| 1-linoleoylglycerol (18:2) | 0.59 | 1.10 | 0.54 |
| 2-palmitoylglycerol (16:0) | 3.14 | 0.50 | 6.35 |
| 1-methylhistidine | 0.56 | 1.52 | 0.37 |
| butyrylcarnitine (C4) | 1.64 | 0.40 | 4.06 |
| isobutyrylcarnitine (C4) | 1.84 | 0.34 | 5.39 |
| 7-ketodeoxycholate | 0.67 | 1.00 | 0.67 |
| 6-oxolithocholate | 1.00 | 0.88 | 1.14 |
| 7-ketolithocholate | 1.00 | 0.92 | 1.09 |
| 2-pyrrolidinone | 0.65 | 1.39 | 0.46 |
| N-(2-furoyl)glycine | 1.34 | 0.60 | 2.25 |
| trigonelline (N'-methylnicotinate) | 1.12 | 0.97 | 1.16 |
| dodecanedioate (C12-DC) | 0.70 | 1.31 | 0.53 |
| N-acetyltyrosine | 0.78 | 1.11 | 0.70 |
| pyridoxate | 0.61 | 1.17 | 0.52 |
| threonylphenylalanine | 1.06 | 0.93 | 1.14 |
| gamma-glutamylvaline | 1.23 | 0.90 | 1.37 |
| pyroglutamylvaline | 1.23 | 0.90 | 1.36 |
| propionylglycine | 1.19 | 0.19 | 6.12 |
| butyrylglycine | 0.98 | 1.09 | 0.90 |
| 2-hydroxyadipate | 0.28 | 2.16 | 0.13 |
| 2-methylbutyrylglycine | 1.19 | 0.48 | 2.46 |
| propionylcarnitine (C3) | 1.75 | 0.26 | 6.77 |
| 3-hydroxy-2-ethylpropionate | 1.03 | 0.70 | 1.47 |
| docosapentaenoate (n3 DPA; 22:5n3) | 0.29 | 1.10 | 0.26 |

| | | | |
|---|---|---|---|
| docosadienoate (22:2n6) | 0.70 | 1.00 | 0.70 |
| adrenate (22:4n6) | 3.29 | 0.85 | 3.86 |
| myristoleate (14:1n5) | 0.72 | 1.19 | 0.60 |
| 4-imidazoleacetate | 0.80 | 1.04 | 0.77 |
| 1-methyl-4-imidazoleacetate | 0.89 | 1.20 | 0.74 |
| sebacate (C10-DC) | 0.56 | 1.61 | 0.35 |
| delta-tocopherol | 0.85 | 1.10 | 0.78 |
| glycitin (glycitein 7-O-glucoside) | 1.06 | 1.00 | 1.06 |
| daidzin (daidzein 7-O-glucoside) | 0.90 | 1.00 | 0.90 |
| daidzein | 0.62 | 1.21 | 0.51 |
| genistin (genistein 7-O-glucoside) | 1.24 | 0.71 | 1.74 |
| genistein | 0.82 | 1.00 | 0.82 |
| glycitein | 0.49 | 1.32 | 0.37 |
| stearidonate (18:4n3) | 0.15 | 1.95 | 0.07 |
| nervonate (24:1n9)* | 0.37 | 1.12 | 0.33 |
| 5-dodecenoate (12:1n7) | 1.01 | 0.98 | 1.04 |
| N-acetylglutamine | 1.20 | 0.89 | 1.34 |
| N-acetyltryptophan | 0.39 | 1.51 | 0.26 |
| N-acetylphenylalanine | 0.75 | 1.28 | 0.59 |
| N-acetylasparagine | 0.73 | 1.13 | 0.65 |
| glycylleucine | 1.08 | 0.94 | 1.15 |
| S-methylglutathione | 1.04 | 0.20 | 5.28 |
| 1-palmitoyl-GPC (16:0) | 0.80 | 1.15 | 0.69 |
| N-acetylarginine | 0.95 | 1.02 | 0.93 |
| piperine | 1.80 | 0.23 | 7.76 |
| campesterol | 0.82 | 1.08 | 0.76 |
| 1-stearoyl-GPC (18:0) | 1.09 | 0.93 | 1.17 |
| 1-oleoyl-GPC (18:1) | 0.91 | 1.15 | 0.79 |
| N-acetylthreonine | 1.06 | 0.99 | 1.07 |
| phenylacetylglycine | 1.63 | 0.52 | 3.14 |
| N-acetylisoleucine | 0.70 | 1.33 | 0.53 |
| 10-nonadecenoate (19:1n9) | 1.35 | 1.00 | 1.35 |
| 10-heptadecenoate (17:1n7) | 1.18 | 1.00 | 1.18 |
| hyocholate (gamma-muricholate) | 1.00 | 0.73 | 1.38 |
| N-acetylhistidine | 0.23 | 1.79 | 0.13 |
| gamma-glutamylglycine | 1.00 | 0.87 | 1.14 |
| gamma-glutamyltryptophan | 1.38 | 0.81 | 1.70 |
| stachydrine | 0.72 | 1.36 | 0.53 |
| alpha-hydroxyisovalerate | 1.04 | 0.97 | 1.07 |
| ophthalmate | 1.68 | 0.41 | 4.12 |
| gamma-glutamylmethionine | 1.03 | 0.98 | 1.05 |
| gamma-glutamylthreonine | 0.96 | 1.15 | 0.83 |
| p-cresol sulfate | 1.60 | 0.47 | 3.40 |

| | | | |
|---|---|---|---|
| nicotinate ribonucleoside | 1.51 | 0.62 | 2.43 |
| erythronate* | 0.55 | 1.71 | 0.32 |
| N-acetylproline | 0.68 | 1.53 | 0.44 |
| eicosenoate (20:1) | 0.41 | 1.03 | 0.40 |
| linolenate [alpha or gamma; (18:3n3 or 6)] | 0.67 | 1.07 | 0.63 |
| aconitate [cis or trans] | 1.27 | 0.80 | 1.59 |
| 1-myristoyl-GPC (14:0) | 0.42 | 1.75 | 0.24 |
| 1-arachidoyl-GPC (20:0) | 0.93 | 1.15 | 0.81 |
| laurylcarnitine (C12) | 0.72 | 1.14 | 0.64 |
| isovalerylcarnitine (C5) | 1.56 | 0.62 | 2.54 |
| 1-linoleoyl-GPC (18:2) | 0.89 | 1.12 | 0.79 |
| 2-myristoylglycerol (14:0) | 3.01 | 0.87 | 3.45 |
| beta-guanidinopropanoate | 1.15 | 0.89 | 1.29 |
| N2,N2-dimethylguanosine | 0.95 | 1.00 | 0.95 |
| N6-carbamoylthreonyladenosine | 1.05 | 0.94 | 1.11 |
| orotidine | 1.05 | 0.90 | 1.16 |
| 5,6-dihydrouridine | 1.18 | 0.86 | 1.37 |
| 3-(3-amino-3-carboxypropyl)uridine* | 0.91 | 1.03 | 0.88 |
| 1-pentadecanoylglycerol (15:0) | 1.67 | 1.00 | 1.67 |
| 1-arachidonylglycerol (20:4) | 1.81 | 0.94 | 1.92 |
| 1-linolenoylglycerol (18:3) | 0.58 | 1.05 | 0.55 |
| cysteine-glutathione disulfide | 1.00 | 0.71 | 1.40 |
| 1-palmitoyl-GPA (16:0) | 0.85 | 1.10 | 0.77 |
| 5-methyluridine (ribothymidine) | 1.07 | 0.98 | 1.10 |
| adenosine 3'-monophosphate (3'-AMP) | 0.64 | 1.74 | 0.37 |
| isovalerylglycine | 0.78 | 1.08 | 0.73 |
| 3-hydroxydodecanedioate* | 0.76 | 1.19 | 0.64 |
| 7-methylguanine | 0.71 | 1.27 | 0.56 |
| 1-stearoyl-GPE (18:0) | 1.24 | 0.78 | 1.59 |
| 1-stearoyl-GPG (18:0) | 1.19 | 1.00 | 1.19 |
| mead acid (20:3n9) | 2.21 | 1.00 | 2.22 |
| 1-docosahexaenoylglycerol (22:6) | 0.07 | 1.04 | 0.06 |
| gamma-glutamylisoleucine* | 1.58 | 0.67 | 2.35 |
| oleoylcarnitine (C18:1) | 0.83 | 1.00 | 0.83 |
| gamma-glutamyl-2-aminobutyrate | 1.00 | 0.54 | 1.85 |
| 4-hydroxybutyrate (GHB) | 0.87 | 1.17 | 0.75 |
| 2-methylbutyrylcarnitine (C5) | 1.54 | 0.55 | 2.83 |
| phenol sulfate | 1.55 | 0.44 | 3.53 |
| 1-palmitoleoyl-GPC (16:1)* | 0.38 | 1.92 | 0.20 |
| hexanoylglycine | 0.98 | 1.02 | 0.95 |
| glutamine_degradant* | 1.23 | 0.88 | 1.40 |
| 2-hydroxy-3-methylvalerate | 0.95 | 1.07 | 0.89 |
| ferulate | 0.44 | 1.54 | 0.29 |

| | | | |
|---|---|---|---|
| homostachydrine* | 0.93 | 1.22 | 0.76 |
| 1-arachidonoyl-GPC (20:4n6)* | 0.58 | 1.50 | 0.39 |
| 1-dihomo-linolenoyl-GPC (20:3n3 or 6)* | 1.07 | 0.95 | 1.12 |
| 1-dihomo-linoleoyl-GPC (20:2)* | 0.99 | 1.00 | 0.99 |
| 2-stearoyl-GPC (18:0)* | 0.44 | 1.14 | 0.39 |
| 2-oleoyl-GPC (18:1)* | 0.51 | 1.14 | 0.45 |
| 2-linoleoyl-GPC (18:2)* | 0.51 | 1.16 | 0.44 |
| 2-palmitoleoyl-GPC (16:1)* | 0.91 | 4.08 | 0.22 |
| 2-palmitoyl-GPC (16:0)* | 0.40 | 1.13 | 0.35 |
| 2-myristoyl-GPC (14:0)* | 0.46 | 1.66 | 0.28 |
| 1-docosahexaenoyl-GPC (22:6)* | 0.01 | 2.62 | 0.00 |
| 1-palmitoyl-GPE (16:0) | 1.04 | 0.94 | 1.11 |
| 1-myristoyl-GPE (14:0) | 0.72 | 1.31 | 0.55 |
| 1-oleoyl-GPE (18:1) | 1.00 | 1.00 | 1.00 |
| 1-linoleoyl-GPE (18:2)* | 1.06 | 0.93 | 1.14 |
| 1-arachidonoyl-GPE (20:4n6)* | 0.89 | 1.00 | 0.89 |
| 2-hydroxypalmitate | 1.19 | 1.00 | 1.19 |
| docosapentaenoate (n6 DPA; 22:5n6) | 0.33 | 1.03 | 0.32 |
| gulonate* | 1.28 | 0.76 | 1.68 |
| isobutyrylglycine | 0.73 | 1.40 | 0.52 |
| phenylpropionylglycine | 1.03 | 0.80 | 1.28 |
| beta-hydroxyisovaleroylcarnitine | 1.39 | 0.53 | 2.63 |
| catechol sulfate | 1.44 | 0.55 | 2.63 |
| cholesterol sulfate | 1.01 | 0.90 | 1.11 |
| octadecanedioate (C18-DC) | 1.23 | 0.82 | 1.50 |
| undecanedioate (C11-DC) | 0.63 | 1.53 | 0.41 |
| 1-myristoylglycerol (14:0) | 3.38 | 0.89 | 3.81 |
| glycerophosphoglycerol | 1.37 | 0.69 | 1.97 |
| glycerophosphoethanolamine | 0.96 | 1.04 | 0.92 |
| glycerophosphoinositol* | 1.17 | 0.83 | 1.41 |
| sedoheptulose-7-phosphate | 1.00 | 0.80 | 1.25 |
| 2-dimethylaminoethanol | 0.72 | 1.23 | 0.59 |
| formononetin | 2.01 | 0.54 | 3.71 |
| vanillate | 0.47 | 1.66 | 0.28 |
| ectoine | 1.95 | 0.22 | 8.94 |
| 2-oleoyl-GPE (18:1)* | 0.69 | 1.00 | 0.69 |
| 1-arachidonoyl-GPI (20:4)* | 0.49 | 1.06 | 0.46 |
| 1-palmitoyl-GPI (16:0) | 0.75 | 1.00 | 0.75 |
| deoxycarnitine | 0.96 | 1.03 | 0.93 |
| N6-succinyladenosine | 0.73 | 1.10 | 0.66 |
| 4-hydroxycinnamate | 0.54 | 1.43 | 0.38 |
| pheophorbide A | 1.39 | 0.78 | 1.78 |
| adenosine 2'-monophosphate (2'-AMP) | 0.55 | 1.59 | 0.34 |

| | | | |
|---|---|---|---|
| 2,4,6-trihydroxybenzoate | 1.15 | 0.66 | 1.75 |
| N2-acetyllysine | 0.06 | 1.05 | 0.06 |
| alpha-hydroxycaproate | 0.46 | 1.02 | 0.45 |
| 3,4-dihydroxybutyrate | 0.28 | 1.78 | 0.16 |
| N6-acetyllysine | 0.54 | 1.52 | 0.35 |
| dihomo-linolenate (20:3n3 or n6) | 2.42 | 0.99 | 2.43 |
| mannitol/sorbitol | 0.87 | 1.06 | 0.82 |
| 4-ethylphenylsulfate | 1.63 | 0.53 | 3.10 |
| pyrraline | 0.09 | 2.45 | 0.04 |
| N6-carboxymethyllysine | 0.05 | 2.33 | 0.02 |
| N6-carboxyethyllysine | 0.22 | 1.90 | 0.12 |
| 2-linoleoyl-GPE (18:2)* | 0.80 | 1.00 | 0.80 |
| 1-oleoyl-GPI (18:1) | 0.65 | 1.07 | 0.60 |
| 1-linoleoyl-GPI (18:2)* | 0.71 | 1.00 | 0.71 |
| 1-palmitoleoyl-GPE (16:1)* | 0.67 | 1.54 | 0.43 |
| glycylisoleucine | 0.91 | 0.94 | 0.98 |
| methylphosphate | 2.02 | 0.26 | 7.71 |
| dimethylarginine (SDMA + ADMA) | 1.09 | 0.86 | 1.27 |
| dihydrokaempferol | 1.05 | 0.99 | 1.05 |
| gamma-glutamylalanine | 1.09 | 0.96 | 1.13 |
| N-acetylserine | 0.97 | 1.02 | 0.96 |
| 1-stearoyl-2-oleoyl-GPE (18:0/18:1) | 1.51 | 0.44 | 3.42 |
| chiro-inositol | 1.15 | 0.95 | 1.21 |
| verbascose | 1.23 | 0.82 | 1.50 |
| 1-stearoyl-2-arachidonoyl-GPC (18:0/20:4) | 1.45 | 0.79 | 1.84 |
| 1-palmitoyl-2-linoleoyl-GPE (16:0/18:2) | 1.25 | 0.72 | 1.75 |
| glycosyl-N-stearoyl-sphingosine (d18:1/18:0) | 1.37 | 0.82 | 1.67 |
| alanylproline | 0.95 | 1.06 | 0.90 |
| alanylleucine | 0.94 | 1.00 | 0.94 |
| syringic acid | 0.42 | 1.47 | 0.28 |
| cyclo(leu-pro) | 0.65 | 1.24 | 0.52 |
| cyclo(phe-pro) | 0.77 | 1.35 | 0.57 |
| cyclo(gly-pro) | 0.35 | 1.87 | 0.19 |
| succinylcarnitine (C4-DC) | 1.00 | 0.96 | 1.04 |
| N-methylproline | 0.92 | 1.00 | 0.92 |
| alpha-tocotrienol | 0.75 | 1.85 | 0.41 |
| gamma-tocotrienol | 0.78 | 1.38 | 0.56 |
| vanillin | 0.12 | 2.01 | 0.06 |
| 5-hydroxymethylfurfural | 0.24 | 1.52 | 0.16 |
| 1-docosahexaenoyl-GPE (22:6)* | 0.01 | 1.18 | 0.01 |
| 2-oleoyl-GPI (18:1)* | 0.59 | 1.02 | 0.58 |
| 2-hydroxyglutarate | 0.77 | 1.16 | 0.67 |
| indole-3-acetamide | 1.39 | 0.73 | 1.90 |

| | | | |
|---|---|---|---|
| palmitoyl sphingomyelin (d18:1/16:0) | 1.77 | 0.32 | 5.49 |
| 3-hydroxyhippurate | 1.07 | 0.38 | 2.81 |
| 5-HEPE | 0.03 | 1.11 | 0.03 |
| naringenin 7-O-glucoside | 1.18 | 0.72 | 1.64 |
| ergothioneine | 1.99 | 0.47 | 4.23 |
| 1-pentadecanoyl-GPC (15:0)* | 0.73 | 1.34 | 0.54 |
| indole-3-carboxylate | 0.84 | 1.16 | 0.73 |
| 13-HODE + 9-HODE | 0.68 | 1.21 | 0.56 |
| hydroquinone beta-glucopyranoside | 0.45 | 1.55 | 0.29 |
| 4-cholesten-3-one | 0.44 | 1.99 | 0.22 |
| N-acetyl-cadaverine | 0.34 | 1.02 | 0.33 |
| cinnamoylglycine | 1.40 | 0.68 | 2.06 |
| stearoyl ethanolamide | 1.79 | 0.20 | 8.81 |
| 2S,3R-dihydroxybutyrate | 0.76 | 1.18 | 0.64 |
| 2,4-dihydroxybutyrate | 0.53 | 1.12 | 0.48 |
| 5-methyl-2'-deoxycytidine | 0.58 | 1.50 | 0.39 |
| S-carboxymethyl-cysteine | 1.10 | 0.95 | 1.15 |
| alpha-tocopherol acetate | 0.37 | 2.30 | 0.16 |
| maltol | 0.73 | 1.14 | 0.64 |
| 2R,3R-dihydroxybutyrate | 0.65 | 1.47 | 0.44 |
| piperidine | 1.49 | 0.53 | 2.79 |
| 2,3-dihydroxyisovalerate | 1.91 | 0.41 | 4.65 |
| 3-methylglutaconate | 0.89 | 1.12 | 0.79 |
| 5-(galactosylhydroxy)-lysine | 0.77 | 1.00 | 0.77 |
| sulfate* | 1.14 | 0.93 | 1.23 |
| 4-hydroxyglutamate | 1.10 | 0.77 | 1.43 |
| homocitrate | 0.55 | 1.47 | 0.37 |
| N-methylhydantoin | 0.08 | 1.97 | 0.04 |
| pantoate | 0.39 | 1.87 | 0.21 |
| sedoheptulose | 1.13 | 0.92 | 1.22 |
| guanosine 3'-monophosphate (3'-GMP) | 0.57 | 1.00 | 0.57 |
| guanosine 2'-monophosphate (2'-GMP)* | 0.59 | 1.05 | 0.56 |
| S-methylcysteine | 0.93 | 0.77 | 1.21 |
| 5,6-DiHETrE | 1.38 | 0.40 | 3.44 |
| argininate* | 1.10 | 0.97 | 1.13 |
| 2-oxoarginine* | 0.69 | 1.32 | 0.52 |
| cis-4-decenoate (10:1n6)* | 0.82 | 0.96 | 0.85 |
| 1-behenoyl-GPC (22:0) | 0.95 | 1.00 | 0.95 |
| 1-erucoyl-GPC (22:1)* | 0.03 | 1.05 | 0.03 |
| 1-adrenoyl-GPC (22:4)* | 0.82 | 0.72 | 1.14 |
| 1-nervonoyl-GPC (24:1n9)* | 0.04 | 1.08 | 0.03 |
| malonylgenistin | 1.87 | 0.12 | 15.36 |
| 1-methyl-5-imidazoleacetate | 0.73 | 1.53 | 0.48 |

| | | | |
|---|---|---|---|
| 3-(4-hydroxyphenyl)propionate | 0.53 | 1.56 | 0.34 |
| S-methylcysteine sulfoxide | 0.84 | 1.11 | 0.76 |
| 16-hydroxypalmitate | 1.44 | 0.62 | 2.31 |
| UDP-N-acetylglucosamine/galactosamine | 1.80 | 0.39 | 4.63 |
| chrysoeriol (3'-O-methylluteolin) | 1.20 | 1.00 | 1.20 |
| oleoyl-linoleoyl-glycerol (18:1/18:2) [1] | 0.36 | 1.79 | 0.20 |
| oleoyl-linoleoyl-glycerol (18:1/18:2) [2] | 0.45 | 1.63 | 0.28 |
| 2-stearoyl-GPI (18:0)* | 0.58 | 1.03 | 0.56 |
| galactosylglycerol | 0.50 | 1.52 | 0.33 |
| digalactosylglycerol* | 1.42 | 0.74 | 1.92 |
| 2-oxindole-3-acetate | 0.55 | 1.27 | 0.44 |
| indoleacetylaspartate | 1.05 | 0.97 | 1.08 |
| N-oleoyltaurine | 0.73 | 1.00 | 0.73 |
| 6'-sialyllactose | 1.00 | 0.51 | 1.97 |
| N-methyl-GABA | 1.39 | 0.92 | 1.51 |
| isoleucylglycine | 0.91 | 1.02 | 0.89 |
| leucylalanine | 0.83 | 1.27 | 0.65 |
| leucylglutamine* | 1.46 | 0.74 | 1.99 |
| leucylglycine | 0.88 | 1.06 | 0.83 |
| lysylleucine | 1.36 | 0.76 | 1.79 |
| valylleucine | 0.96 | 1.08 | 0.88 |
| N-palmitoyltaurine | 0.79 | 1.00 | 0.79 |
| 2-O-methylascorbic acid | 1.54 | 0.56 | 2.73 |
| carboxyethyl-GABA | 0.71 | 1.37 | 0.52 |
| beta-citrylglutamate | 1.40 | 0.65 | 2.15 |
| hydroxymethylpyrimidine | 0.09 | 2.05 | 0.04 |
| trimethylamine N-oxide | 1.81 | 0.43 | 4.26 |
| 3'-sialyllactose | 1.22 | 0.04 | 29.03 |
| N6-methyllysine | 0.71 | 1.27 | 0.56 |
| cis-urocanate | 0.07 | 1.98 | 0.03 |
| equol sulfate | 1.00 | 0.86 | 1.16 |
| dihydroferulate | 1.85 | 0.47 | 3.97 |
| imidazole propionate | 0.03 | 2.32 | 0.01 |
| tyrosol | 1.74 | 0.33 | 5.33 |
| alanyllysine | 0.93 | 1.08 | 0.86 |
| lysylalanine | 1.13 | 0.88 | 1.28 |
| phenylalanylglycine | 0.88 | 1.01 | 0.87 |
| phenylalanylalanine | 0.95 | 1.16 | 0.82 |
| aspartylaspartate | 1.14 | 0.95 | 1.20 |
| valylglutamine | 0.98 | 1.01 | 0.96 |
| valylglycine | 0.87 | 1.09 | 0.80 |
| prolylalanine | 0.78 | 1.08 | 0.73 |
| prolylglycine | 0.43 | 1.08 | 0.40 |

| | | | |
|---|---|---|---|
| prolylproline | 1.11 | 0.98 | 1.13 |
| N-palmitoylglycine | 1.02 | 1.00 | 1.02 |
| mannonate* | 0.67 | 1.34 | 0.50 |
| 4-methylthio-2-oxobutanoate | 0.25 | 1.95 | 0.13 |
| glycohyodeoxycholate | 0.77 | 0.77 | 1.00 |
| lanthionine | 1.11 | 0.12 | 9.51 |
| equol glucuronide | 0.81 | 1.09 | 0.74 |
| 2-stearoyl-GPE (18:0)* | 0.48 | 1.17 | 0.41 |
| (R)-3-hydroxybutyrylcarnitine | 0.54 | 1.00 | 0.54 |
| nicotianamine | 1.71 | 0.41 | 4.18 |
| 2'-O-methyladenosine | 0.80 | 1.04 | 0.77 |
| histidine betaine (hercynine)* | 1.02 | 0.97 | 1.04 |
| 4-methylcatechol sulfate | 1.61 | 0.47 | 3.41 |
| 3-dehydrochenodeoxycholate | 1.27 | 0.75 | 1.69 |
| tryptophol | 1.05 | 0.97 | 1.08 |
| dimethyl sulfone | 1.64 | 0.47 | 3.51 |
| o-tyrosine | 0.81 | 1.18 | 0.68 |
| 2-piperidinone | 0.48 | 1.60 | 0.30 |
| 2,8-quinolinediol sulfate | 1.61 | 0.50 | 3.23 |
| 1-stearoyl-GPS (18:0)* | 0.49 | 0.49 | 1.00 |
| sphingomyelin (d18:1/14:0, d16:1/16:0)* | 0.86 | 1.09 | 0.79 |
| sphingomyelin (d18:2/16:0, d18:1/16:1)* | 1.06 | 0.18 | 6.06 |
| syringaresinol | 0.59 | 1.12 | 0.53 |
| cyclo(pro-tyr) | 0.57 | 1.50 | 0.38 |
| 3-hydroxyadipate | 1.34 | 0.40 | 3.40 |
| 1-methyl-beta-carboline-3-carboxylic acid | 1.50 | 0.64 | 2.34 |
| soyasaponin I | 1.18 | 0.94 | 1.25 |
| soyasaponin II | 1.03 | 1.00 | 1.03 |
| soyasaponin III | 1.29 | 0.79 | 1.63 |
| N-monomethylarginine | 1.18 | 0.77 | 1.52 |
| myristoyl ethanolamide | 1.24 | 0.71 | 1.74 |
| 6-oxopiperidine-2-carboxylate | 0.52 | 1.42 | 0.37 |
| N-delta-acetylornithine | 1.13 | 0.99 | 1.14 |
| 2-aminoheptanoate | 0.90 | 0.96 | 0.94 |
| 1-eicosapentaenoyl-GPE (20:5)* | 0.85 | 1.00 | 0.85 |
| N-formylanthranilic acid | 0.71 | 1.44 | 0.49 |
| 1-linoleoyl-GPS (18:2)* | 1.01 | 1.00 | 1.01 |
| docosahexaenoyl ethanolamide | 0.80 | 1.00 | 0.80 |
| methionine sulfone | 1.20 | 0.85 | 1.41 |
| cyclo(phe-pro) (L,D)* | 0.90 | 1.00 | 0.90 |
| 1-linolenoyl-GPC (18:3)* | 0.89 | 1.11 | 0.80 |
| 1-eicosapentaenoyl-GPC (20:5)* | 0.00 | 1.30 | 0.00 |
| 1-eicosenoyl-GPC (20:1)* | 0.68 | 1.59 | 0.43 |

| | | | |
|---|---|---|---|
| 1-dihomo-linolenoyl-GPE (20:3n3 or 6)* | 1.09 | 0.62 | 1.75 |
| fructosyllysine | 0.03 | 2.21 | 0.01 |
| 1-eicosenoyl-GPE (20:1)* | 0.55 | 1.72 | 0.32 |
| O-sulfo-tyrosine | 0.73 | 1.11 | 0.66 |
| ferulic acid 4-sulfate | 1.32 | 0.44 | 2.98 |
| salidroside | 1.76 | 0.34 | 5.15 |
| N-acetylmethionine sulfoxide | 0.29 | 2.06 | 0.14 |
| N-acetyltaurine | 0.54 | 1.57 | 0.35 |
| 1-linolenoyl-GPI (18:3)* | 0.84 | 1.00 | 0.84 |
| 2-linolenoyl-GPI (18:3)* | 0.99 | 0.85 | 1.16 |
| 1-linolenoyl-GPE (18:3)* | 1.15 | 0.85 | 1.35 |
| 1-oleoyl-GPG (18:1)* | 0.69 | 1.01 | 0.68 |
| 1-palmitoyl-GPG (16:0)* | 0.85 | 1.00 | 0.85 |
| 2-palmitoyl-GPG (16:0)* | 0.97 | 1.00 | 0.97 |
| 2-oleoyl-GPG (18:1)* | 0.82 | 1.00 | 0.82 |
| 1-palmitoyl-GPS (16:0)* | 0.94 | 0.87 | 1.07 |
| N-linoleoyltaurine* | 0.93 | 1.00 | 0.93 |
| methyl glucopyranoside (alpha + beta) | 1.00 | 0.86 | 1.16 |
| acetylagmatine | 0.71 | 1.38 | 0.51 |
| 2-keto-3-deoxy-gluconate | 0.43 | 1.43 | 0.30 |
| sphingomyelin (d18:1/24:1, d18:2/24:0)* | 0.48 | 1.60 | 0.30 |
| 7-hydroxycholesterol (alpha or beta) | 0.72 | 1.27 | 0.57 |
| diosmin (diosmetin 7-rutinoside) | 0.14 | 2.42 | 0.06 |
| myristoleoylcarnitine (C14:1)* | 0.36 | 1.06 | 0.34 |
| N-formylphenylalanine | 0.67 | 1.24 | 0.53 |
| cyclo(pro-val) | 0.73 | 1.20 | 0.61 |
| 3-hydroxypyridine sulfate | 1.00 | 0.82 | 1.22 |
| arabonate/xylonate | 0.78 | 1.32 | 0.59 |
| 1-dihomo-linolenylglycerol (20:3) | 1.80 | 0.99 | 1.81 |
| N-acetylhistamine | 0.32 | 1.54 | 0.21 |
| vanillactate | 0.59 | 1.43 | 0.41 |
| sphingomyelin (d18:1/20:0, d16:1/22:0)* | 1.76 | 0.21 | 8.40 |
| sphingomyelin (d18:1/20:1, d18:2/20:0)* | 1.12 | 0.48 | 2.32 |
| behenoyl sphingomyelin (d18:1/22:0)* | 1.74 | 0.23 | 7.63 |
| sphingomyelin (d18:1/22:1, d18:2/22:0, d16:1/24:1)* | 0.67 | 1.29 | 0.52 |
| lignoceroyl sphingomyelin (d18:1/24:0) | 1.93 | 0.19 | 10.40 |
| sphingomyelin (d17:1/16:0, d18:1/15:0, d16:1/17:0)* | 1.73 | 0.33 | 5.17 |
| kaempferol 3-O-glucoside/galactoside | 1.04 | 0.99 | 1.05 |
| 3-hydroxyhexanoate | 0.73 | 1.06 | 0.69 |
| N-carbamoylalanine | 0.31 | 1.64 | 0.19 |
| gentisic acid-5-glucoside | 1.40 | 0.83 | 1.68 |
| arabitol/xylitol | 0.79 | 1.22 | 0.64 |

| | | | |
|---|---|---|---|
| N-acetylglucosamine/N-acetylgalactosamine | 1.43 | 0.61 | 2.37 |
| citraconate/glutaconate | 0.22 | 1.99 | 0.11 |
| linoleoyl ethanolamide | 0.93 | 1.09 | 0.85 |
| cyclo(met-pro) | 0.66 | 1.38 | 0.48 |
| 1,2-dilinoleoyl-GPC (18:2/18:2) | 1.22 | 0.82 | 1.48 |
| 1-stearoyl-2-oleoyl-GPC (18:0/18:1) | 1.16 | 0.81 | 1.42 |
| 1,2-dioleoyl-GPE (18:1/18:1) | 1.25 | 0.79 | 1.59 |
| 1-palmitoyl-2-arachidonoyl-GPC (16:0/20:4n6) | 0.77 | 1.00 | 0.77 |
| 1-palmitoyl-2-docosahexaenoyl-GPC (16:0/22:6) | 0.03 | 1.16 | 0.02 |
| 1-stearoyl-2-docosahexaenoyl-GPC (18:0/22:6) | 0.02 | 1.16 | 0.02 |
| sphingomyelin (d18:1/17:0, d17:1/18:0, d19:1/16:0) | 1.04 | 0.12 | 8.98 |
| 1-palmitoyl-2-stearoyl-GPC (16:0/18:0) | 1.42 | 0.61 | 2.34 |
| 2-hydroxybutyrate/2-hydroxyisobutyrate | 0.82 | 1.23 | 0.67 |
| oleate/vaccenate (18:1) | 0.97 | 1.00 | 0.97 |
| 1-palmitoleoylglycerol (16:1)* | 0.86 | 1.00 | 0.86 |
| 2-palmitoleoylglycerol (16:1)* | 0.92 | 0.96 | 0.96 |
| palmitoyl dihydrosphingomyelin (d18:0/16:0)* | 1.72 | 0.26 | 6.68 |
| tricosanoyl sphingomyelin (d18:1/23:0)* | 1.94 | 0.16 | 12.45 |
| sphingomyelin (d18:2/23:0, d18:1/23:1, d17:1/24:1)* | 1.34 | 0.55 | 2.42 |
| sphingomyelin (d18:2/24:1, d18:1/24:2)* | 1.20 | 0.93 | 1.30 |
| uridine 2'-monophosphate (2'-UMP)* | 0.44 | 1.23 | 0.36 |
| 1-stearoyl-2-linoleoyl-GPE (18:0/18:2)* | 1.53 | 0.58 | 2.64 |
| 1-stearoyl-2-arachidonoyl-GPE (18:0/20:4) | 1.11 | 0.24 | 4.65 |
| 1-stearoyl-2-linoleoyl-GPC (18:0/18:2)* | 1.17 | 0.84 | 1.40 |
| 1-oleoyl-2-linoleoyl-GPC (18:1/18:2)* | 1.08 | 0.89 | 1.22 |
| 1-palmitoyl-2-palmitoleoyl-GPC (16:0/16:1)* | 0.84 | 1.34 | 0.63 |
| 1-palmitoyl-2-eicosapentaenoyl-GPC (16:0/20:5)* | 0.06 | 1.11 | 0.06 |
| 1-palmitoyl-2-arachidonoyl-GPE (16:0/20:4)* | 1.25 | 0.22 | 5.74 |
| 1-palmitoyl-2-docosahexaenoyl-GPE (16:0/22:6)* | 0.80 | 1.00 | 0.80 |
| 1-stearoyl-2-linoleoyl-GPI (18:0/18:2) | 1.32 | 0.71 | 1.85 |
| 1-palmitoyl-2-palmitoleoyl-GPE (16:0/16:1)* | 1.49 | 0.53 | 2.82 |
| gamma-tocopherol/beta-tocopherol | 0.83 | 1.26 | 0.66 |
| 1-(1-enyl-palmitoyl)-2-oleoyl-GPE (P-16:0/18:1)* | 1.13 | 0.52 | 2.18 |
| 1-palmityl-2-arachidonoyl-GPC (O-16:0/20:4)* | 0.81 | 0.81 | 1.00 |
| sphingomyelin (d18:1/21:0, d17:1/22:0, d16:1/23:0)* | 1.75 | 0.22 | 8.06 |
| behenoyl dihydrosphingomyelin (d18:0/22:0)* | 1.00 | 0.95 | 1.05 |
| sphingomyelin (d18:0/18:0, d19:0/17:0)* | 1.00 | 0.90 | 1.11 |
| N-palmitoyl-sphinganine (d18:0/16:0) | 1.08 | 0.92 | 1.19 |
| lactosyl-N-palmitoyl-sphingosine (d18:1/16:0) | 1.03 | 0.94 | 1.10 |
| 1-pentadecanoyl-2-linoleoyl-GPC (15:0/18:2)* | 1.35 | 0.79 | 1.70 |
| 1-margaroyl-2-oleoyl-GPC (17:0/18:1)* | 1.17 | 0.86 | 1.36 |
| 1-margaroyl-2-linoleoyl-GPC (17:0/18:2)* | 1.36 | 0.73 | 1.87 |

| | | | |
|---|---|---|---|
| myristoyl dihydrosphingomyelin (d18:0/14:0)* | 1.44 | 0.64 | 2.25 |
| palmitoyl-linoleoyl-glycerol (16:0/18:2) [1]* | 0.54 | 1.50 | 0.36 |
| palmitoyl-linoleoyl-glycerol (16:0/18:2) [2]* | 0.68 | 1.29 | 0.53 |
| (3'-5')-guanylyluridine | 0.51 | 1.06 | 0.48 |
| (3'-5')-uridylyluridine | 0.78 | 1.10 | 0.71 |
| (3'-5')-adenylyluridine | 0.30 | 1.21 | 0.24 |
| (3'-5')-cytidylyladenosine | 0.45 | 1.02 | 0.44 |
| 1-palmitoyl-2-oleoyl-GPI (16:0/18:1)* | 1.10 | 0.95 | 1.16 |
| 1-stearoyl-2-docosahexaenoyl-GPI (18:0/22:6)* | 0.61 | 1.00 | 0.61 |
| 1,2-dilinoleoyl-GPA (18:2/18:2)* | 1.00 | 0.92 | 1.09 |
| 1-palmitoleoyl-2-linoleoyl-GPC (16:1/18:2)* | 1.43 | 0.59 | 2.44 |
| 1-oleoyl-2-linoleoyl-GPE (18:1/18:2)* | 1.25 | 0.79 | 1.58 |
| 1-linoleoyl-GPA (18:2)* | 0.89 | 1.10 | 0.81 |
| 1-pentadecanoyl-2-docosahexaenoyl-GPC (15:0/22:6)* | 0.02 | 1.01 | 0.02 |
| 1-oleoyl-2-docosahexaenoyl-GPC (18:1/22:6)* | 0.84 | 1.00 | 0.84 |
| 1-linoleoyl-2-docosahexaenoyl-GPC (18:2/22:6)* | 0.81 | 1.00 | 0.81 |
| 1-palmityl-2-linoleoyl-GPC (O-16:0/18:2)* | 1.02 | 1.00 | 1.02 |
| 1-myristoyl-2-arachidonoyl-GPC (14:0/20:4)* | 0.82 | 1.00 | 0.82 |
| 1-stearoyl-2-docosapentaenoyl-GPC (18:0/22:5n3)* | 0.53 | 1.41 | 0.37 |
| 1-stearyl-GPC (O-18:0)* | 0.50 | 1.67 | 0.30 |
| 1-stearoyl-2-meadoyl-GPC (18:0/20:3n9)* | 1.36 | 0.62 | 2.18 |
| 1-palmitoyl-2-gamma-linolenoyl-GPC (16:0/18:3n6)* | 1.21 | 0.78 | 1.55 |
| 1-(1-enyl-palmitoyl)-2-palmitoyl-GPC (P-16:0/16:0)* | 0.99 | 1.03 | 0.96 |
| 1-palmitoyl-2-palmitoleoyl-GPI (16:0/16:1)* | 1.45 | 0.59 | 2.47 |
| 1-stearoyl-2-oleoyl-GPI (18:0/18:1)* | 1.44 | 0.51 | 2.81 |
| 1-stearoyl-2-dihomo-linolenoyl-GPI (18:0/20:3n3 or 6)* | 1.00 | 0.46 | 2.16 |
| 1,2-dilinoleoyl-GPI (18:2/18:2)* | 1.19 | 0.80 | 1.49 |
| 1,2-dipalmitoyl-GPE (16:0/16:0)* | 1.09 | 0.98 | 1.11 |
| 1,2-dilinoleoyl-GPE (18:2/18:2)* | 1.30 | 0.72 | 1.80 |
| 1-linoleoyl-2-linolenoyl-GPE (18:2/18:3)* | 1.47 | 0.59 | 2.50 |
| 1-linoleoyl-2-arachidonoyl-GPE (18:2/20:4)* | 0.84 | 1.00 | 0.84 |
| 1-linoleoyl-GPG (18:2)* | 0.84 | 1.00 | 0.84 |
| thioproline | 1.00 | 0.85 | 1.17 |
| palmitoylcholine | 0.73 | 1.37 | 0.53 |
| isocitric lactone | 0.78 | 1.07 | 0.73 |
| 2-methylserine | 0.87 | 1.07 | 0.81 |
| (S)-3-hydroxybutyrylcarnitine | 1.50 | 0.72 | 2.09 |
| glycosyl-N-palmitoyl-sphingosine (d18:1/16:0) | 1.56 | 0.48 | 3.25 |
| 14-HDoHE/17-HDoHE | 0.10 | 2.14 | 0.05 |
| 3-carboxyadipate | 0.60 | 1.35 | 0.44 |
| oleoylcholine | 0.54 | 1.65 | 0.33 |
| docosahexaenoylcholine | 0.70 | 1.00 | 0.70 |

| | | | |
|---|---|---|---|
| palmitoloelycholine | 0.24 | 2.21 | 0.11 |
| 1-linoleoyl-2-linolenoyl-GPC (18:2/18:3)* | 1.34 | 0.65 | 2.07 |
| 1,2-dilinolenoyl-GPC (18:3/18:3)* | 1.48 | 0.60 | 2.47 |
| 1-palmitoleoyl-2-linolenoyl-GPC (16:1/18:3)* | 1.40 | 0.63 | 2.22 |
| phosphatidylcholine (18:0/20:2, 20:0/18:2)* | 1.15 | 0.80 | 1.43 |
| 1,2-dilinolenoyl-digalactosylglycerol (18:3/18:3) | 1.29 | 0.85 | 1.51 |
| hexadecatrienoate (16:3n3) | 0.12 | 2.22 | 0.05 |
| hexadecadienoate (16:2n6) | 0.34 | 1.45 | 0.23 |
| 1-eicosapentaenoylglycerol (20:5)* | 0.81 | 1.00 | 0.81 |
| palmitoleoylcarnitine (C16:1)* | 0.91 | 1.00 | 0.91 |
| 3-methylglutarate/2-methylglutarate | 0.66 | 1.11 | 0.60 |
| 2-hydroxyoleate | 0.60 | 1.23 | 0.49 |
| 2,3-dihydroxy-2-methylbutyrate | 0.79 | 1.06 | 0.74 |
| 1-linoleoyl-2-linolenoyl-digalactosylglycerol (18:2/18:3)* | 1.04 | 0.89 | 1.17 |
| 1,2-dilinoleoyl-digalactosylglycerol (18:2/18:2)* | 1.00 | 1.02 | 0.98 |
| 1-palmitoyl-2-linoleoyl-digalactosylglycerol (16:0/18:2)* | 1.00 | 0.95 | 1.05 |
| 1-palmitoyl-2-linolenoyl-digalactosylglycerol (16:0/18:3) | 1.00 | 1.05 | 0.95 |
| 1,2-dilinolenoyl-galactosylglycerol (18:3/18:3)* | 0.94 | 1.08 | 0.87 |
| 1-linoleoyl-2-linolenoyl-galactosylglycerol (18:2/18:3)* | 0.95 | 1.03 | 0.92 |
| 1,2-dilinoleoyl-galactosylglycerol (18:2/18:2)* | 1.02 | 0.94 | 1.09 |
| 1-linoleoyl-galactosylglycerol (18:2)* | 0.78 | 1.00 | 0.78 |
| 1-linolenoyl-galactosylglycerol (18:3)* | 1.05 | 0.95 | 1.10 |
| 1-palmitoyl-galactosylglycerol (16:0)* | 0.74 | 1.02 | 0.72 |
| 1-linoleoyl-digalactosylglycerol (18:2)* | 0.76 | 1.00 | 0.76 |
| 1-linolenoyl-digalactosylglycerol (18:3)* | 0.99 | 1.05 | 0.94 |
| 1-palmitoyl-digalactosylglycerol (16:0)* | 0.68 | 1.01 | 0.68 |
| 1-palmitoyl-2-linoleoyl-galactosylglycerol (16:0/18:2)* | 1.15 | 0.85 | 1.35 |
| 1-palmitoyl-2-linolenoyl-galactosylglycerol (16:0/18:3)* | 1.05 | 0.93 | 1.13 |
| 2'-O-methylcytidine | 0.89 | 0.95 | 0.94 |
| 2'-O-methyluridine | 1.04 | 0.98 | 1.06 |
| gamma-glutamyl-alpha-lysine | 0.32 | 1.77 | 0.18 |
| coumaroylquinate (2) | 0.43 | 1.90 | 0.22 |
| coumaroylquinate (3) | 0.77 | 1.27 | 0.61 |
| coumaroylquinate (5) | 0.79 | 1.35 | 0.59 |
| palmitoyl-oleoyl-glycerol (16:0/18:1) [1]* | 0.68 | 1.28 | 0.53 |
| palmitoyl-oleoyl-glycerol (16:0/18:1) [2]* | 0.97 | 1.00 | 0.97 |
| oleoyl-oleoyl-glycerol (18:1/18:1) [1]* | 0.45 | 1.73 | 0.26 |
| oleoyl-oleoyl-glycerol (18:1/18:1) [2]* | 0.54 | 1.51 | 0.36 |
| stearoyl-linoleoyl-glycerol (18:0/18:2) [1]* | 0.75 | 1.33 | 0.57 |
| stearoyl-linoleoyl-glycerol (18:0/18:2) [2]* | 0.69 | 1.22 | 0.56 |
| linoleoyl-arachidonoyl-glycerol (18:2/20:4) [1]* | 0.64 | 3.02 | 0.21 |

| | | | |
|---|---|---|---|
| linoleoyl-arachidonoyl-glycerol (18:2/20:4) [2]* | 0.23 | 1.88 | 0.12 |
| palmitoyl-arachidonoyl-glycerol (16:0/20:4) [1]* | 1.00 | 0.61 | 1.64 |
| palmitoyl-arachidonoyl-glycerol (16:0/20:4) [2]* | 0.89 | 1.32 | 0.68 |
| linoleoyl-linolenoyl-glycerol (18:2/18:3) [1]* | 0.41 | 1.73 | 0.24 |
| linoleoyl-linolenoyl-glycerol (18:2/18:3) [2]* | 0.56 | 1.41 | 0.40 |
| linoleoyl-docosahexaenoyl-glycerol (18:2/22:6) [1]* | 0.67 | 1.00 | 0.67 |
| linoleoyl-docosahexaenoyl-glycerol (18:2/22:6) [2]* | 0.70 | 1.00 | 0.70 |
| palmitoleoyl-linoleoyl-glycerol (16:1/18:2) [1]* | 0.27 | 2.00 | 0.14 |
| linolenoyl-linolenoyl-glycerol (18:3/18:3) [1]* | 0.51 | 1.61 | 0.32 |
| linolenoyl-linolenoyl-glycerol (18:3/18:3) [2]* | 0.89 | 1.45 | 0.61 |
| diacylglycerol (14:0/18:1, 16:0/16:1) [1]* | 0.76 | 1.10 | 0.69 |
| diacylglycerol (14:0/18:1, 16:0/16:1) [2]* | 1.20 | 0.72 | 1.68 |
| oleoyl-arachidonoyl-glycerol (18:1/20:4) [1]* | 0.87 | 1.46 | 0.59 |
| oleoyl-arachidonoyl-glycerol (18:1/20:4) [2]* | 1.04 | 0.48 | 2.18 |
| palmitoyl-linolenoyl-glycerol (16:0/18:3) [2]* | 0.77 | 1.27 | 0.61 |
| diacylglycerol (16:1/18:2 [2], 16:0/18:3 [1])* | 0.54 | 1.51 | 0.36 |
| linoleoyl-linoleoyl-glycerol (18:2/18:2) [1]* | 0.37 | 1.68 | 0.22 |
| linoleoyl-linoleoyl-glycerol (18:2/18:2) [2]* | 0.51 | 1.55 | 0.33 |
| palmitoyl-palmitoyl-glycerol (16:0/16:0) [2]* | 0.92 | 0.82 | 1.12 |
| stearoyl-arachidonoyl-glycerol (18:0/20:4) [1]* | 1.00 | 1.00 | 1.00 |
| stearoyl-arachidonoyl-glycerol (18:0/20:4) [2]* | 1.17 | 0.77 | 1.51 |
| 1-stearoyl-2-dihomo-linolenoyl-GPS (18:0/20:3n3 or 6)* | 1.00 | 0.82 | 1.22 |
| 1-oleoyl-2-eicosenoyl-GPC (18:1/20:1)* | 0.79 | 1.29 | 0.61 |
| 1-palmityl-2-palmitoyl-GPC (O-16:0/16:0)* | 0.60 | 1.03 | 0.58 |
| 2-hydroxybehenate | 1.06 | 1.00 | 1.06 |
| 2-hydroxynervonate* | 0.69 | 1.01 | 0.68 |
| sphingadienine | 0.74 | 1.30 | 0.57 |
| glycerophosphoserine* | 1.06 | 0.95 | 1.12 |
| 1-stearoyl-2-(hydroxylinoleoyl)-GPC (18:0/18:2(OH))* | 1.17 | 0.88 | 1.32 |
| 1-palmitoyl-2-(hydroxylinoleoyl)-GPC (16:0/18:2(OH))* | 1.23 | 0.90 | 1.37 |
| ceramide (d16:1/24:1, d18:1/22:1)* | 0.25 | 1.89 | 0.13 |
| glycosyl ceramide (d16:1/24:1, d18:1/22:1)* | 0.02 | 1.06 | 0.02 |
| ceramide (d18:1/14:0, d16:1/16:0)* | 0.23 | 1.78 | 0.13 |
| ceramide (d18:2/24:1, d18:1/24:2)* | 1.28 | 0.65 | 1.97 |
| stearoylcholine* | 0.98 | 1.00 | 0.98 |
| linoleoylcholine* | 0.63 | 1.59 | 0.40 |
| 1-arachidoyl-GPE (20:0)* | 1.16 | 0.69 | 1.69 |
| sphingomyelin (d18:0/20:0, d16:0/22:0)* | 1.80 | 0.18 | 9.73 |
| sphingomyelin (d18:1/19:0, d19:1/18:0)* | 1.00 | 0.85 | 1.18 |
| sphingomyelin (d18:2/21:0, d16:2/23:0)* | 1.00 | 0.90 | 1.11 |
| heneicosapentaenoate (21:5n3) | 0.85 | 1.00 | 0.85 |
| eicosenoylcarnitine (C20:1)* | 0.86 | 1.00 | 0.86 |

| | | | |
|---|---|---|---|
| palmitoleoyl ethanolamide* | 1.00 | 1.03 | 0.97 |
| glycosyl ceramide (d18:1/20:0, d16:1/22:0)* | 1.54 | 0.32 | 4.82 |
| N,N,N-trimethyl-5-aminovalerate | 1.23 | 0.29 | 4.21 |
| ethyl alpha-glucopyranoside | 1.92 | 0.48 | 3.98 |
| carotene diol (1) | 1.14 | 0.87 | 1.31 |
| carotene diol (2) | 1.36 | 0.82 | 1.65 |
| carotene diol (3) | 0.85 | 1.21 | 0.70 |
| 1-palmitoyl-2-pentadecanoyl-GPC (16:0/15:0)* | 1.51 | 0.51 | 2.93 |
| (N(1) + N(8))-acetylspermidine | 0.73 | 1.07 | 0.68 |
| 2-linolenoyl-galactosylglycerol (18:3)* | 1.10 | 0.91 | 1.21 |
| N-palmitoyl-phytosphingosine (t18:0/16:0) | 0.99 | 1.02 | 0.97 |
| linolenoylcholine* | 0.76 | 1.33 | 0.57 |
| 2-hydroxyarachidate* | 1.34 | 1.00 | 1.34 |
| cytidine 2' or 3'-monophosphate (2' or 3'-CMP) | 0.66 | 1.00 | 0.66 |
| N-linoleoylserine* | 1.34 | 0.97 | 1.38 |
| 2-hydroxyheptanoate* | 0.64 | 1.47 | 0.44 |
| lyxonate | 0.72 | 1.33 | 0.54 |
| dodecenedioate (C12:1-DC)* | 0.64 | 1.68 | 0.38 |
| octadecenedioate (C18:1-DC) | 1.27 | 0.80 | 1.59 |
| octadecadienedioate (C18:2-DC)* | 0.75 | 1.40 | 0.54 |
| N,N,N-trimethyl-alanylproline betaine (TMAP) | 0.97 | 1.00 | 0.97 |
| 3-formylindole | 0.34 | 1.95 | 0.18 |
| pyroglutamylalanine* | 1.14 | 0.92 | 1.24 |
| tetradecadienoate (14:2)* | 0.64 | 1.37 | 0.47 |
| 3-amino-2-piperidone | 0.49 | 1.08 | 0.45 |
| N,N-dimethylalanine | 0.95 | 1.02 | 0.93 |
| 3-indoleglyoxylic acid | 0.71 | 1.41 | 0.50 |
| ethyl beta-glucopyranoside | 0.55 | 1.27 | 0.43 |
| 2-hydroxysebacate | 0.46 | 1.72 | 0.27 |
| enterolactone sulfate | 1.97 | 0.28 | 6.98 |
| N6,N6-dimethyllysine | 0.83 | 1.00 | 0.83 |
| N-acetyl-isoputreanine | 1.01 | 0.89 | 1.13 |
| 4-ethylcatechol sulfate | 1.60 | 0.57 | 2.79 |
| 4-allylcatechol sulfate | 1.00 | 0.85 | 1.17 |
| 2-linolenoyl-digalactosylglycerol (18:3)* | 1.07 | 0.81 | 1.32 |
| 1-linolenoyl-GPG (18:3)* | 1.22 | 1.00 | 1.22 |
| 2-palmitoyl-galactosylglycerol (16:0)* | 0.96 | 0.89 | 1.09 |
| 2-palmitoyl-digalactosylglycerol (16:0)* | 0.74 | 0.97 | 0.77 |
| heptadecatrienoate (17:3)* | 0.70 | 1.10 | 0.64 |
| 1-nonadecenoyl-GPC (19:1)* | 0.82 | 1.17 | 0.70 |
| 2,6-dihydroxybenzoic acid | 0.28 | 1.83 | 0.15 |
| linolenoyl ethanolamide | 0.89 | 1.07 | 0.83 |
| methylthioadenosine sulfoxide | 1.33 | 0.82 | 1.62 |

| | | | |
|---|---|---|---|
| phenylacetylaspartate | 1.29 | 0.91 | 1.41 |
| N-oleoylalanine | 0.61 | 1.00 | 0.61 |
| cyclo(pro-arg)* | 0.60 | 1.37 | 0.44 |
| coixol | 0.29 | 1.26 | 0.23 |
| isoxanthohumol | 1.00 | 0.43 | 2.31 |
| eicosenedioate (C20:1-DC)* | 1.21 | 0.54 | 2.23 |
| dehydrophytosphingosine* | 0.77 | 1.54 | 0.50 |
| 5-hydroxypicolinic acid | 0.48 | 1.92 | 0.25 |
| leu-val-val | 0.77 | 1.00 | 0.77 |
| (S)-a-amino-omega-caprolactam | 0.18 | 1.99 | 0.09 |
| 2-hydroxy-4-(methylthio)butanoic acid | 0.74 | 1.05 | 0.70 |
| lariciresinol 4-O-glucoside | 1.20 | 0.98 | 1.23 |
| apigenin glucuronide (2) | 0.57 | 1.20 | 0.47 |
| pentose acid* | 1.11 | 0.94 | 1.18 |
| N-succinyl-leucine | 0.52 | 1.28 | 0.41 |
| N-succinyl-isoleucine | 1.20 | 0.94 | 1.28 |
| 1-methyl-5-imidazolelactate | 0.48 | 1.89 | 0.26 |
| feruloylquinate (1) | 1.00 | 1.00 | 1.00 |
| feruloylquinate (2) | 0.42 | 1.78 | 0.24 |
| feruloylquinate (3) | 0.92 | 1.14 | 0.81 |
| feruloylquinate (4) | 0.65 | 1.43 | 0.46 |
| feruloylquinate (5) | 0.60 | 1.67 | 0.36 |
| N-carbamoylputrescine | 0.94 | 1.00 | 0.94 |
| N1,N10-dicoumaroylspermidine | 0.56 | 1.50 | 0.37 |
| ala-ile-ala | 1.14 | 0.82 | 1.39 |
| val-val-ala | 0.86 | 1.07 | 0.80 |
| branched-chain, straight-chain, or cyclopropyl 10:1 fatty acid (1)* | 1.41 | 0.75 | 1.88 |
| N-acetylmethionylphenylalanine | 0.29 | 1.14 | 0.26 |
| N-acetyl-2-aminoadipate | 1.87 | 0.48 | 3.90 |
| flavone derivative C26H28O14 (1)* | 1.86 | 0.22 | 8.34 |
| flavone derivative C26H28O14 (2)* | 1.98 | 0.18 | 11.08 |
| flavone derivative C26H28O14 (3)* | 1.88 | 0.23 | 8.21 |
| flavone derivative C26H28O14 (4)* | 1.87 | 0.24 | 7.83 |
| N,N-dimethyl-pro-pro | 1.06 | 0.98 | 1.09 |
| oxindolylalanine | 0.42 | 1.82 | 0.23 |
| N-lactoyl histidine | 1.11 | 0.97 | 1.15 |
| N-lactoyl isoleucine | 1.17 | 0.87 | 1.35 |
| N-lactoyl leucine | 1.63 | 0.61 | 2.66 |
| N-lactoyl phenylalanine | 1.50 | 0.65 | 2.32 |
| N-lactoyl tyrosine | 1.08 | 0.95 | 1.14 |
| N-lactoyl valine | 1.14 | 0.87 | 1.30 |
| 1-oleyl-GPC (O-18:1)* | 0.33 | 1.98 | 0.17 |
| ceramide (d18:2/16:0, d18:1/16:1, d16:1/18:1)* | 1.46 | 0.49 | 2.99 |

*Supplemental Table 4: Differentially abundant stool metabolites of sl/sl fed the Protective or Detrimental diet at CHOP (ANCOVA contrast, p<0.05)*

| Chemical name | Prot. / Det. Fold-change | p-value | q-value |
|---|---|---|---|
| daidzein | 0.15 | 4.53E-02 | 0.32 |
| octadecenedioylcarnitine (C18:1-DC)* | 0.15 | 1.50E-03 | 0.17 |
| stearidonate (18:4n3) | 0.15 | 0.00E+00 | 0.00 |
| putrescine | 0.17 | 4.33E-02 | 0.32 |
| N1,N10-dicoumaroylspermidine | 0.19 | 4.30E-02 | 0.32 |
| laurylcarnitine (C12) | 0.21 | 4.60E-03 | 0.18 |
| undecenoylcarnitine (C11:1) | 0.27 | 5.80E-03 | 0.18 |
| thiamin (vitamin B1) | 0.28 | 5.00E-04 | 0.17 |
| gamma-glutamyl-epsilon-lysine | 0.28 | 1.30E-03 | 0.17 |
| myristoleoylcarnitine (C14:1)* | 0.29 | 6.40E-03 | 0.18 |
| decanoylcarnitine (C10) | 0.29 | 2.90E-03 | 0.18 |
| leucylalanine | 0.31 | 1.36E-02 | 0.23 |
| dihydrobiopterin | 0.31 | 1.61E-02 | 0.23 |
| succinylglutamine | 0.32 | 9.90E-03 | 0.21 |
| leucylglutamine* | 0.32 | 2.16E-02 | 0.25 |
| creatinine | 0.32 | 2.63E-02 | 0.27 |
| valylleucine | 0.33 | 5.60E-03 | 0.18 |
| alanylleucine | 0.34 | 2.59E-02 | 0.27 |
| N-acetylhomocitrulline | 0.34 | 1.03E-02 | 0.21 |
| linolenoylcarnitine (C18:3)* | 0.34 | 3.32E-02 | 0.30 |
| glutarylcarnitine (C5-DC) | 0.35 | 1.31E-02 | 0.23 |
| glutaminylleucine | 0.35 | 4.27E-02 | 0.32 |
| adipoylcarnitine (C6-DC) | 0.35 | 2.30E-02 | 0.26 |
| erucate (22:1n9) | 0.35 | 2.45E-02 | 0.27 |
| threonylphenylalanine | 0.36 | 1.41E-02 | 0.23 |
| phenylalanylalanine | 0.36 | 3.42E-02 | 0.30 |
| inositol hexakisphosphate | 0.37 | 6.70E-03 | 0.18 |
| leucylglycine | 0.37 | 2.00E-02 | 0.24 |
| N-acetyltryptophan | 0.40 | 2.27E-02 | 0.26 |
| 2-aminophenol | 0.40 | 2.28E-02 | 0.26 |
| N-acetylpyrraline | 0.40 | 2.87E-02 | 0.29 |
| octadecanedioylcarnitine (C18-DC)* | 0.42 | 1.16E-02 | 0.22 |
| valylglutamine | 0.42 | 4.16E-02 | 0.32 |
| N-acetylcysteine | 0.42 | 7.40E-03 | 0.19 |
| N-acetyl-beta-glucosaminylamine | 0.43 | 2.48E-02 | 0.27 |
| octanoylcarnitine (C8) | 0.43 | 3.02E-02 | 0.30 |
| hydantoin-5-propionate | 0.44 | 4.60E-03 | 0.18 |
| nicotinamide | 0.45 | 4.62E-02 | 0.32 |
| 1-methylguanidine | 0.45 | 1.86E-02 | 0.23 |
| phenol glucuronide | 0.45 | 3.40E-03 | 0.18 |
| pimeloylcarnitine/3-methyladipoylcarnitine (C7-DC) | 0.45 | 2.03E-02 | 0.24 |
| pyrraline | 0.46 | 1.80E-02 | 0.23 |
| octadecenedioate (C18:1-DC) | 0.47 | 1.03E-02 | 0.21 |
| biopterin | 0.48 | 4.42E-02 | 0.32 |
| cytidine 5'-monophosphate (5'-CMP) | 0.49 | 3.74E-02 | 0.31 |
| hexadecadienoate (16:2n6) | 0.51 | 7.10E-03 | 0.18 |
| 1-linoleoylglycerol (18:2) | 0.52 | 3.72E-02 | 0.31 |
| 1-linoleoyl-GPC (18:2) | 0.53 | 4.29E-02 | 0.32 |
| 13-HODE + 9-HODE | 0.55 | 3.91E-02 | 0.31 |
| flavone derivative C26H28O14 (5)* | 0.56 | 1.69E-02 | 0.23 |
| hexadecenedioate (C16:1-DC)* | 0.57 | 1.08E-02 | 0.21 |
| hexadecanedioate (C16-DC) | 0.62 | 4.60E-02 | 0.32 |
| 1-nervonoyl-GPC (24:1n9)* | 0.65 | 9.90E-03 | 0.21 |
| (3'-5')-uridylylguanosine | 0.76 | 3.54E-02 | 0.31 |
| androstenediol (3beta,17beta) monosulfate (2) | 1.41 | 3.37E-02 | 0.30 |

| | | | |
|---|---|---|---|
| pyridoxal | 1.45 | 4.43E-02 | 0.32 |
| nicotinate | 1.49 | 5.90E-03 | 0.18 |
| benzoate | 1.59 | 4.97E-02 | 0.33 |
| mevalonolactone | 1.69 | 3.19E-02 | 0.30 |
| 1,2-dipalmitoyl-GPE (16:0/16:0)* | 1.72 | 1.67E-02 | 0.23 |
| pseudouridine | 1.79 | 3.23E-02 | 0.30 |
| 2-hydroxyheptanoate* | 1.89 | 1.17E-02 | 0.22 |
| uracil | 1.89 | 4.22E-02 | 0.32 |
| 1-palmitoyl-2-oleoyl-GPE (16:0/18:1) | 1.92 | 1.40E-02 | 0.23 |
| diacetylchitobiose | 1.92 | 3.36E-02 | 0.30 |
| 2-palmitoylglycerol (16:0) | 1.96 | 1.54E-02 | 0.23 |
| 1-(1-enyl-palmitoyl)-2-oleoyl-GPE (P-16:0/18:1)* | 1.96 | 1.67E-02 | 0.23 |
| flavin adenine dinucleotide (FAD) | 1.96 | 2.63E-02 | 0.27 |
| phenylpyruvate | 1.96 | 3.20E-02 | 0.30 |
| 1-palmitoyl-2-dihomo-linolenoyl-GPE (16:0/20:3)* | 1.96 | 3.88E-02 | 0.31 |
| xanthine | 2.08 | 1.51E-02 | 0.23 |
| sedoheptulose | 2.08 | 2.40E-02 | 0.26 |
| 1-stearoyl-2-oleoyl-GPC (18:0/18:1) | 2.08 | 3.35E-02 | 0.30 |
| 1-palmitoyl-2-oleoyl-GPC (16:0/18:1) | 2.08 | 4.26E-02 | 0.32 |
| indolepropionate | 2.13 | 2.76E-02 | 0.28 |
| 3-methyl-2-oxobutyrate | 2.17 | 3.38E-02 | 0.30 |
| dehydrophytosphingosine* | 2.17 | 3.93E-02 | 0.31 |
| fucose | 2.17 | 4.85E-02 | 0.33 |
| beta-cryptoxanthin | 2.22 | 2.00E-03 | 0.17 |
| 1-stearoyl-2-dihomo-linolenoyl-GPC (18:0/20:3n3 or 6)* | 2.22 | 4.50E-02 | 0.32 |
| 1-stearoyl-2-linoleoyl-GPC (18:0/18:2)* | 2.33 | 3.75E-02 | 0.31 |
| 1-palmitoyl-2-docosahexaenoyl-GPE (16:0/22:6)* | 2.33 | 4.94E-02 | 0.33 |
| nicotinamide adenine dinucleotide (NAD+) | 2.38 | 2.20E-03 | 0.17 |
| 1-margaroyl-2-linoleoyl-GPC (17:0/18:2)* | 2.38 | 1.45E-02 | 0.23 |
| 1-pentadecanoyl-2-linoleoyl-GPC (15:0/18:2)* | 2.50 | 1.30E-03 | 0.17 |
| 5,6-dihydrouridine | 2.50 | 5.10E-03 | 0.18 |
| palmitoyl-oleoyl-glycerol (16:0/18:1) [2]* | 2.56 | 8.90E-03 | 0.21 |
| 2-methylcitrate/homocitrate | 2.56 | 1.20E-02 | 0.22 |
| 1-oleoyl-2-dihomo-linolenoyl-GPC (18:1/20:3)* | 2.63 | 1.41E-02 | 0.23 |
| 1-palmitoyl-2-arachidonoyl-GPE (16:0/20:4)* | 2.63 | 1.67E-02 | 0.23 |
| 1-(1-enyl-stearoyl)-2-docosahexaenoyl-GPE (P-18:0/22:6)* | 2.63 | 3.92E-02 | 0.31 |
| 1-palmitoyl-2-oleoyl-GPG (16:0/18:1) | 2.70 | 3.60E-03 | 0.18 |
| 1-stearoyl-2-dihomo-linolenoyl-GPE (18:0/20:3n3 or 6)* | 2.70 | 1.39E-02 | 0.23 |
| isovalerate (i5:0) | 2.78 | 4.03E-02 | 0.32 |
| 1-oleoyl-2-arachidonoyl-GPC (18:1/20:4)* | 2.86 | 4.79E-02 | 0.33 |
| 3-dehydroshikimate | 3.03 | 6.10E-03 | 0.18 |
| 3beta-hydroxy-5-cholenoate | 3.03 | 8.80E-03 | 0.21 |
| 1-(1-enyl-stearoyl)-2-arachidonoyl-GPE (P-18:0/20:4)* | 3.03 | 1.89E-02 | 0.23 |
| 1-palmitoyl-2-dihomo-linolenoyl-GPC (16:0/20:3n3 or 6)* | 3.03 | 2.14E-02 | 0.25 |
| pheophytin A | 3.13 | 1.80E-03 | 0.17 |
| stearoyl ethanolamide | 3.23 | 4.10E-03 | 0.18 |
| oxalate (ethanedioate) | 3.23 | 1.01E-02 | 0.21 |
| 1-stearoyl-2-linoleoyl-GPE (18:0/18:2)* | 3.23 | 1.71E-02 | 0.23 |
| alpha-ketoglutarate | 3.33 | 2.90E-03 | 0.18 |
| 1-(1-enyl-stearoyl)-2-oleoyl-GPE (P-18:0/18:1) | 3.33 | 4.30E-03 | 0.18 |
| 1-oleoyl-2-linoleoyl-GPE (18:1/18:2)* | 3.33 | 1.56E-02 | 0.23 |
| valerate (5:0) | 3.33 | 3.63E-02 | 0.31 |
| nicotinate ribonucleoside | 3.45 | 4.92E-02 | 0.33 |
| 1-stearoyl-2-arachidonoyl-GPI (18:0/20:4) | 3.57 | 1.02E-02 | 0.21 |
| 1-stearoyl-2-arachidonoyl-GPE (18:0/20:4) | 3.57 | 1.77E-02 | 0.23 |
| 1-stearoyl-2-oleoyl-GPE (18:0/18:1) | 3.85 | 3.70E-03 | 0.18 |
| ergothioneine | 4.00 | 1.30E-03 | 0.17 |
| gamma-glutamylglutamate | 4.00 | 3.67E-02 | 0.31 |
| butyrate/isobutyrate (4:0) | 4.76 | 4.50E-03 | 0.18 |

| | | | |
|---|---|---|---|
| maltol | 5.00 | 1.00E-03 | 0.17 |
| 1-stearoyl-2-oleoyl-GPS (18:0/18:1) | 5.00 | 5.10E-03 | 0.18 |
| 3,5-dihydroxybenzoic acid | 14.29 | 6.30E-03 | 0.18 |

*Supplemental Table 5: Differentially abundant stool metabolites of sl/sl fed the Protective or Detrimental diet at UQAM (ANCOVA contrast, p<0.05)*

| Chemical name | Prot. / Det. Fold-change | p-value | q-value |
|---|---|---|---|
| thiamin (vitamin B1) | 0.05 | 0.00E+00 | 0.00 |
| cadaverine | 0.06 | 1.15E-02 | 0.05 |
| putrescine | 0.07 | 4.20E-03 | 0.02 |
| 1-palmityl-GPE (O-16:0)* | 0.12 | 1.00E-04 | 0.00 |
| gamma-glutamyl-epsilon-lysine | 0.13 | 0.00E+00 | 0.00 |
| 1-oleyl-GPC (O-18:1)* | 0.13 | 2.10E-03 | 0.02 |
| 1-palmityl-GPC (O-16:0) | 0.13 | 4.70E-03 | 0.03 |
| stearidonate (18:4n3) | 0.14 | 0.00E+00 | 0.00 |
| succinylglutamine | 0.14 | 0.00E+00 | 0.00 |
| urate | 0.14 | 0.00E+00 | 0.00 |
| N6,N6-dimethyllysine | 0.16 | 6.40E-03 | 0.03 |
| docosahexaenoate (DHA; 22:6n3) | 0.17 | 6.80E-03 | 0.03 |
| allantoin | 0.17 | 1.14E-02 | 0.05 |
| erucate (22:1n9) | 0.18 | 5.00E-04 | 0.01 |
| cystine | 0.18 | 2.90E-03 | 0.02 |
| agmatine | 0.18 | 4.00E-03 | 0.02 |
| ferulic acid 4-sulfate | 0.18 | 2.88E-02 | 0.09 |
| ceramide (d18:2/24:1, d18:1/24:2)* | 0.18 | 3.07E-02 | 0.09 |
| 1-(1-enyl-oleoyl)-GPC (P-18:1)* | 0.19 | 1.50E-03 | 0.01 |
| oxindolylalanine | 0.2 | 0.00E+00 | 0.00 |
| eicosapentaenoate (EPA; 20:5n3) | 0.2 | 1.00E-04 | 0.00 |
| hydroxymethylpyrimidine | 0.2 | 5.00E-04 | 0.01 |
| flavone derivative C26H28O14 (1)* | 0.2 | 5.60E-03 | 0.03 |
| vanillactate | 0.22 | 5.00E-04 | 0.01 |
| 1-(1-enyl-palmitoyl)-GPC (P-16:0)* | 0.22 | 4.00E-03 | 0.02 |
| pyroglutamylvaline | 0.23 | 6.00E-04 | 0.01 |
| N-acetylputrescine | 0.23 | 1.02E-02 | 0.04 |
| 4-methylthio-2-oxobutanoate | 0.24 | 0.00E+00 | 0.00 |
| 3-formylindole | 0.24 | 4.00E-04 | 0.01 |
| cholate sulfate | 0.24 | 1.40E-03 | 0.01 |
| creatinine | 0.24 | 6.00E-03 | 0.03 |
| pyrraline | 0.25 | 1.00E-04 | 0.00 |
| cysteine s-sulfate | 0.25 | 4.00E-04 | 0.01 |
| 2-hydroxy-4-(methylthio)butanoic acid | 0.25 | 3.90E-03 | 0.02 |
| linoleoyl-arachidonoyl-glycerol (18:2/20:4) [2]* | 0.25 | 2.66E-02 | 0.08 |
| 3-methyl-2-oxobutyrate | 0.26 | 5.00E-04 | 0.01 |
| thymine | 0.26 | 1.50E-03 | 0.01 |
| uridine 5'-monophosphate (UMP) | 0.26 | 2.30E-03 | 0.02 |
| flavone derivative C26H28O14 (5)* | 0.27 | 0.00E+00 | 0.00 |
| 5-(2-hydroxyethyl)-4-methylthiazole | 0.27 | 1.50E-03 | 0.01 |
| N-acetylpyrraline | 0.27 | 2.20E-03 | 0.02 |
| glycylvaline | 0.27 | 4.10E-03 | 0.02 |
| 2,4-dihydroxybutyrate | 0.27 | 9.00E-03 | 0.04 |
| 5-hydroxylysine | 0.28 | 1.00E-04 | 0.00 |
| 4-methyl-2-oxopentanoate | 0.28 | 7.00E-04 | 0.01 |
| pregnenediol disulfate (C21H34O8S2)* | 0.28 | 3.70E-03 | 0.02 |
| p-cresol sulfate | 0.28 | 2.26E-02 | 0.07 |
| o-tyrosine | 0.29 | 0.00E+00 | 0.00 |
| 3-hydroxymargaroylglycine | 0.29 | 4.00E-04 | 0.01 |
| 4-hydroxyphenylpyruvate | 0.29 | 1.90E-03 | 0.02 |

| | | | |
|---|---|---|---|
| methionine sulfoxide | 0.3 | 1.00E-04 | 0.00 |
| pyroglutamylalanine* | 0.3 | 1.00E-04 | 0.00 |
| p-hydroxybenzaldehyde | 0.3 | 4.00E-04 | 0.01 |
| 3-methyl-2-oxovalerate | 0.3 | 7.00E-04 | 0.01 |
| 4-cholesten-3-one | 0.3 | 9.00E-04 | 0.01 |
| 1-methylhistidine | 0.3 | 2.10E-03 | 0.02 |
| 1-(1-enyl-palmitoyl)-GPE (P-16:0)* | 0.3 | 2.30E-03 | 0.02 |
| pantoate | 0.3 | 7.70E-03 | 0.04 |
| prolylglycine | 0.3 | 1.83E-02 | 0.06 |
| O-sulfo-tyrosine | 0.3 | 3.36E-02 | 0.09 |
| N6-acetyllysine | 0.31 | 2.00E-04 | 0.00 |
| valine | 0.31 | 2.00E-04 | 0.01 |
| histidine | 0.31 | 7.00E-04 | 0.01 |
| carboxymethylproline | 0.31 | 1.60E-03 | 0.01 |
| pyroglutamylglycine | 0.31 | 2.50E-03 | 0.02 |
| itaconate | 0.31 | 4.60E-03 | 0.03 |
| glycylleucine | 0.31 | 6.90E-03 | 0.03 |
| dihomo-linolenate (20:3n3 or n6) | 0.31 | 2.77E-02 | 0.08 |
| 2-oxoarginine* | 0.31 | 3.53E-02 | 0.10 |
| isoleucine | 0.32 | 2.00E-04 | 0.01 |
| 2,3-dihydroxyisovalerate | 0.32 | 2.40E-03 | 0.02 |
| N-acetylglucosaminylasparagine | 0.32 | 2.96E-02 | 0.09 |
| malate | 0.33 | 1.00E-04 | 0.00 |
| 1-linoleoylglycerol (18:2) | 0.33 | 7.00E-04 | 0.01 |
| 5-(galactosylhydroxy)-lysine | 0.33 | 6.20E-03 | 0.03 |
| glycylisoleucine | 0.33 | 1.64E-02 | 0.06 |
| phenol glucuronide | 0.34 | 1.00E-04 | 0.00 |
| serine | 0.34 | 2.00E-04 | 0.01 |
| tryptophan | 0.34 | 1.00E-03 | 0.01 |
| 1-methylguanidine | 0.34 | 2.00E-03 | 0.02 |
| gamma-glutamylleucine | 0.34 | 2.10E-03 | 0.02 |
| 1-(1-enyl-linoleoyl)-GPE (P-18:2)* | 0.34 | 5.40E-03 | 0.03 |
| 1-dihomo-linolenylglycerol (20:3) | 0.34 | 9.90E-03 | 0.04 |
| spermidine | 0.34 | 2.12E-02 | 0.07 |
| asparagine | 0.34 | 4.19E-02 | 0.11 |
| leucine | 0.35 | 9.00E-04 | 0.01 |
| 2-linoleoylglycerol (18:2) | 0.35 | 1.00E-03 | 0.01 |
| gamma-glutamylmethionine | 0.35 | 2.30E-03 | 0.02 |
| 1-(1-enyl-oleoyl)-GPE (P-18:1)* | 0.35 | 3.70E-03 | 0.02 |
| threonate | 0.35 | 1.00E-02 | 0.04 |
| glycylproline | 0.36 | 3.30E-03 | 0.02 |
| pyruvate | 0.36 | 8.40E-03 | 0.04 |
| prolylproline | 0.36 | 1.75E-02 | 0.06 |
| 1-(1-enyl-stearoyl)-GPC (P-18:0)* | 0.36 | 4.07E-02 | 0.11 |
| arachidonate (20:4n6) | 0.36 | 4.60E-02 | 0.12 |
| tyrosine | 0.37 | 1.00E-03 | 0.01 |
| threonine | 0.37 | 1.10E-03 | 0.01 |
| phenylpyruvate | 0.37 | 2.20E-03 | 0.02 |
| N6-methyllysine | 0.37 | 6.70E-03 | 0.03 |
| 2'-deoxycytidine 5'-monophosphate | 0.37 | 2.74E-02 | 0.08 |
| 5-oxoproline | 0.37 | 4.33E-02 | 0.11 |
| glycine | 0.38 | 4.00E-04 | 0.01 |
| phenylalanine | 0.38 | 1.10E-03 | 0.01 |
| 3-hydroxyadipate | 0.38 | 2.00E-03 | 0.02 |
| N-acetylmethionine sulfoxide | 0.38 | 7.30E-03 | 0.04 |
| caffeate | 0.38 | 2.25E-02 | 0.07 |
| methionine | 0.4 | 3.30E-03 | 0.02 |
| 1-nervonoyl-GPC (24:1n9)* | 0.41 | 0.00E+00 | 0.00 |
| ornithine | 0.41 | 3.09E-02 | 0.09 |

| | | | |
|---|---|---|---|
| 2'-deoxyadenosine 5'-monophosphate | 0.41 | 3.97E-02 | 0.11 |
| 2R,3R-dihydroxybutyrate | 0.42 | 3.00E-04 | 0.01 |
| alanine | 0.42 | 7.00E-04 | 0.01 |
| fumarate | 0.42 | 2.20E-03 | 0.02 |
| gamma-glutamyltyrosine | 0.42 | 6.40E-03 | 0.03 |
| palmitoleate (16:1n7) | 0.42 | 1.70E-02 | 0.06 |
| N-acetyl-beta-glucosaminylamine | 0.42 | 2.36E-02 | 0.07 |
| proline | 0.43 | 2.00E-04 | 0.01 |
| hexadecenedioate (C16:1-DC)* | 0.44 | 4.00E-04 | 0.01 |
| 2-oleoylglycerol (18:1) | 0.44 | 4.40E-03 | 0.03 |
| gamma-glutamylglycine | 0.44 | 4.81E-02 | 0.12 |
| linoleate (18:2n6) | 0.45 | 2.35E-02 | 0.07 |
| erythronate* | 0.45 | 4.46E-02 | 0.11 |
| kynurenine | 0.46 | 2.00E-04 | 0.01 |
| anthranilate | 0.46 | 9.00E-04 | 0.01 |
| glycerophosphoethanolamine | 0.46 | 1.25E-02 | 0.05 |
| N1-methyl-2-pyridone-5-carboxamide | 0.46 | 1.51E-02 | 0.06 |
| hexadecadienoate (16:2n6) | 0.47 | 2.90E-03 | 0.02 |
| N-acetylneuraminate | 0.47 | 1.03E-02 | 0.04 |
| adenosine 5'-monophosphate (AMP) | 0.47 | 1.51E-02 | 0.06 |
| uracil | 0.47 | 1.70E-02 | 0.06 |
| diaminopimelate | 0.48 | 2.43E-02 | 0.08 |
| 1-eicosenoyl-GPE (20:1)* | 0.48 | 3.37E-02 | 0.09 |
| cytidine 5'-monophosphate (5'-CMP) | 0.48 | 3.46E-02 | 0.10 |
| nervonoylcarnitine (C24:1)* | 0.48 | 4.37E-02 | 0.11 |
| 3-amino-2-piperidone | 0.49 | 2.72E-02 | 0.08 |
| 2-keto-3-deoxy-gluconate | 0.49 | 3.36E-02 | 0.09 |
| xanthine | 0.5 | 1.96E-02 | 0.07 |
| citrulline | 0.5 | 4.18E-02 | 0.11 |
| N2-acetyllysine | 0.51 | 3.80E-03 | 0.02 |
| lysine | 0.53 | 1.83E-02 | 0.06 |
| 1-oleoylglycerol (18:1) | 0.53 | 2.94E-02 | 0.09 |
| N-lactoyl phenylalanine | 0.54 | 2.25E-02 | 0.07 |
| tyrosol | 0.54 | 4.28E-02 | 0.11 |
| pseudouridine | 0.55 | 2.60E-02 | 0.08 |
| S-1-pyrroline-5-carboxylate | 0.56 | 6.00E-03 | 0.03 |
| trans-4-hydroxyproline | 0.6 | 1.25E-02 | 0.05 |
| pipecolate | 0.6 | 1.72E-02 | 0.06 |
| 3-sulfo-alanine | 0.61 | 1.66E-02 | 0.06 |
| benzoate | 0.61 | 4.01E-02 | 0.11 |
| sulindac | 0.62 | 2.07E-02 | 0.07 |
| gamma-glutamyltryptophan | 0.62 | 2.10E-02 | 0.07 |
| glutamate | 0.64 | 2.30E-02 | 0.07 |
| carnitine | 0.65 | 3.69E-02 | 0.10 |
| maleate | 0.69 | 1.16E-02 | 0.05 |
| (3'-5')-uridylylguanosine | 0.7 | 8.10E-03 | 0.04 |
| nicotinate | 1.43 | 1.41E-02 | 0.05 |
| 3-hydroxyhexanoate | 1.44 | 3.90E-03 | 0.02 |
| alpha-tocopherol | 1.58 | 4.00E-04 | 0.01 |
| linoleoylcholine* | 1.67 | 3.50E-02 | 0.10 |
| campesterol | 1.69 | 8.70E-03 | 0.04 |
| 1-palmitoyl-2-oleoyl-GPE (16:0/18:1) | 1.72 | 3.90E-02 | 0.11 |
| 1-oleoyl-GPA (18:1) | 1.79 | 4.56E-02 | 0.12 |
| flavin adenine dinucleotide (FAD) | 1.89 | 3.49E-02 | 0.10 |
| 1-palmitoyl-2-linoleoyl-GPE (16:0/18:2) | 1.9 | 3.69E-02 | 0.10 |
| carotene diol (3) | 1.91 | 4.74E-02 | 0.12 |
| hydroxy-undecanedioate (OH-C11:0-DC)* | 1.95 | 7.30E-03 | 0.04 |
| beta-cryptoxanthin | 1.95 | 8.80E-03 | 0.04 |
| docosadioate (C22-DC) | 1.97 | 6.50E-03 | 0.03 |

| | | | |
|---|---|---|---|
| N-carbamoylputrescine | 2.06 | 3.27E-02 | 0.09 |
| beta-sitosterol | 2.09 | 0.00E+00 | 0.00 |
| flavin mononucleotide (FMN) | 2.1 | 2.06E-02 | 0.07 |
| fucosterol | 2.16 | 5.00E-04 | 0.01 |
| ergosterol | 2.17 | 1.22E-02 | 0.05 |
| 1-linoleoyl-2-linolenoyl-GPC (18:2/18:3)* | 2.19 | 2.25E-02 | 0.07 |
| pterin | 2.21 | 4.47E-02 | 0.11 |
| octadecanedioylcarnitine (C18-DC)* | 2.22 | 2.03E-02 | 0.07 |
| 2-methylcitrate/homocitrate | 2.24 | 2.73E-02 | 0.08 |
| 1-(1-enyl-stearoyl)-2-oleoyl-GPE (P-18:0/18:1) | 2.25 | 4.63E-02 | 0.12 |
| 1-palmitoyl-2-linoleoyl-galactosylglycerol (16:0/18:2)* | 2.29 | 1.18E-02 | 0.05 |
| 1-palmitoyl-2-arachidonoyl-GPE (16:0/20:4)* | 2.33 | 3.36E-02 | 0.09 |
| stigmasterol | 2.35 | 4.00E-04 | 0.01 |
| N-acetyl-3-methylhistidine* | 2.35 | 3.25E-02 | 0.09 |
| 2-palmitoylglycerol (16:0) | 2.38 | 2.50E-03 | 0.02 |
| 1-linoleoyl-GPE (18:2)* | 2.41 | 9.00E-03 | 0.04 |
| N,N,N-trimethyl-5-aminovalerate | 2.41 | 1.93E-02 | 0.07 |
| pheophytin A | 2.42 | 1.26E-02 | 0.05 |
| 1-stearoyl-2-dihomo-linolenoyl-GPC (18:0/20:3n3 or 6)* | 2.43 | 2.51E-02 | 0.08 |
| 3,4-dihydroxybenzoate | 2.49 | 1.66E-02 | 0.06 |
| pantethine | 2.49 | 3.14E-02 | 0.09 |
| 1-stearoyl-2-linoleoyl-GPC (18:0/18:2)* | 2.51 | 2.55E-02 | 0.08 |
| indolin-2-one | 2.54 | 1.85E-02 | 0.06 |
| 1-palmitoyl-GPA (16:0) | 2.55 | 4.31E-02 | 0.11 |
| serotonin | 2.65 | 1.32E-02 | 0.05 |
| 1-oleoyl-2-linoleoyl-GPC (18:1/18:2)* | 2.66 | 1.18E-02 | 0.05 |
| protoporphyrin IX | 2.71 | 8.20E-03 | 0.04 |
| 1-pentadecanoyl-2-linoleoyl-GPC (15:0/18:2)* | 2.72 | 5.00E-04 | 0.01 |
| 3-hydroxyoleoylcarnitine | 2.73 | 1.50E-02 | 0.06 |
| 1-margaroyl-2-linoleoyl-GPC (17:0/18:2)* | 2.74 | 5.10E-03 | 0.03 |
| maltotriose | 2.79 | 4.15E-02 | 0.11 |
| 1-stearoyl-2-dihomo-linolenoyl-GPE (18:0/20:3n3 or 6)* | 2.81 | 1.03E-02 | 0.04 |
| pimeloylcarnitine/3-methyladipoylcarnitine (C7-DC) | 2.82 | 3.00E-03 | 0.02 |
| 1-palmitoyl-2-oleoyl-GPG (16:0/18:1) | 2.83 | 2.70E-03 | 0.02 |
| 2-aminoadipate | 2.86 | 1.00E-04 | 0.00 |
| 1-stearoyl-2-arachidonoyl-GPE (18:0/20:4) | 2.86 | 4.76E-02 | 0.12 |
| ectoine | 2.9 | 4.64E-02 | 0.12 |
| beta-hydroxyisovaleroylcarnitine | 2.98 | 1.08E-02 | 0.04 |
| heme | 2.98 | 4.43E-02 | 0.11 |
| gamma-tocopherol/beta-tocopherol | 3.12 | 2.60E-03 | 0.02 |
| maltol | 3.12 | 1.64E-02 | 0.06 |
| palmitoyl-oleoyl-glycerol (16:0/18:1) [2]* | 3.16 | 2.00E-03 | 0.02 |
| adenine | 3.19 | 2.18E-02 | 0.07 |
| 1,2-dilinoleoyl-GPE (18:2/18:2)* | 3.22 | 2.90E-03 | 0.02 |
| enterodiol | 3.42 | 1.05E-02 | 0.04 |
| tauroursodeoxycholate | 3.65 | 2.08E-02 | 0.07 |
| dehydrophytosphingosine* | 3.7 | 9.00E-04 | 0.01 |
| 3-(4-hydroxyphenyl)propionate | 4.02 | 9.40E-03 | 0.04 |
| 1-stearoyl-2-linoleoyl-GPE (18:0/18:2)* | 4.05 | 5.40E-03 | 0.03 |
| butyrate/isobutyrate (4:0) | 4.25 | 7.60E-03 | 0.04 |
| valerate (5:0) | 4.45 | 1.01E-02 | 0.04 |
| enterolactone | 4.56 | 8.50E-03 | 0.04 |
| 6beta-hydroxylithocholate | 4.82 | 9.20E-03 | 0.04 |
| 1-stearoyl-2-oleoyl-GPS (18:0/18:1) | 4.85 | 6.10E-03 | 0.03 |
| N-methylproline | 5.83 | 3.70E-03 | 0.02 |
| pantetheine | 6.02 | 6.00E-04 | 0.01 |
| N,N-dimethylalanine | 8.51 | 1.14E-02 | 0.05 |
| nicotinate ribonucleoside | 8.56 | 1.30E-03 | 0.01 |
| 3-(3-hydroxyphenyl)propionate | 8.7 | 4.99E-02 | 0.12 |

| | | | |
|---|---|---|---|
| equol | 11.78 | 2.89E-02 | 0.09 |
| 2,8-quinolinediol | 14.44 | 8.10E-03 | 0.04 |
| hyodeoxycholate | 15.28 | 6.90E-03 | 0.03 |
| 3,5-dihydroxybenzoic acid | 19.92 | 2.40E-03 | 0.02 |

*Supplemental Table 6: Relative abundance of all metabolites in stool of sl/sl fed the Protective or Detrimental diets regardless of location (>2-fold difference, q<0.05)*

| Chemical name | Prot. food relative abundance (mean) | Det. food relative abundance (mean) | Prot. / Det. Fold-change |
|---|---|---|---|
| putrescine | 0.09 | 0.85 | 0.11 |
| thiamin (vitamin B1) | 0.42 | 3.42 | 0.12 |
| stearidonate (18:4n3) | 0.22 | 1.56 | 0.14 |
| gamma-glutamyl-epsilon-lysine | 0.28 | 1.45 | 0.19 |
| succinylglutamine | 0.39 | 1.83 | 0.21 |
| erucate (22:1n9) | 0.58 | 2.27 | 0.25 |
| 1-palmityl-GPE (O-16:0)* | 0.65 | 2.33 | 0.28 |
| flavone derivative C26H28O14 (1)* | 0.22 | 0.79 | 0.28 |
| creatinine | 0.55 | 1.97 | 0.28 |
| cystine | 0.64 | 2.05 | 0.31 |
| N-acetylpyrraline | 0.22 | 0.67 | 0.33 |
| eicosapentaenoate (EPA; 20:5n3) | 0.58 | 1.77 | 0.33 |
| pyrraline | 0.36 | 1.07 | 0.34 |
| glycylvaline | 0.53 | 1.55 | 0.34 |
| N-acetylputrescine | 0.58 | 1.67 | 0.35 |
| p-cresol sulfate | 0.30 | 0.83 | 0.36 |
| cholate sulfate | 0.58 | 1.58 | 0.37 |
| glycylleucine | 0.50 | 1.33 | 0.38 |
| glycylisoleucine | 0.45 | 1.16 | 0.38 |
| uridine 5'-monophosphate (UMP) | 0.78 | 2.04 | 0.38 |
| phenol glucuronide | 0.25 | 0.65 | 0.39 |
| pregnenediol disulfate (C21H34O8S2)* | 0.55 | 1.40 | 0.39 |
| flavone derivative C26H28O14 (5)* | 0.30 | 0.76 | 0.39 |
| 1-methylguanidine | 0.47 | 1.19 | 0.39 |
| vanillactate | 0.43 | 1.09 | 0.40 |
| 2,4-dihydroxybutyrate | 0.93 | 2.35 | 0.40 |
| 1-linoleoylglycerol (18:2) | 0.69 | 1.66 | 0.41 |
| 5-hydroxylysine | 0.63 | 1.49 | 0.42 |
| hydroxymethylpyrimidine | 0.54 | 1.28 | 0.43 |
| N-acetyl-beta-glucosaminylamine | 0.54 | 1.26 | 0.43 |
| itaconate | 0.47 | 1.09 | 0.43 |
| pyroglutamylvaline | 0.45 | 1.03 | 0.43 |
| urate | 0.84 | 1.93 | 0.43 |
| N-acetyltryptophan | 0.52 | 1.17 | 0.44 |
| 2-linoleoylglycerol (18:2) | 0.74 | 1.67 | 0.45 |
| cysteine s-sulfate | 0.84 | 1.85 | 0.45 |
| methionine sulfoxide | 0.77 | 1.67 | 0.46 |
| 4-cholesten-3-one | 0.66 | 1.42 | 0.46 |
| oxindolylalanine | 0.43 | 0.94 | 0.46 |
| glycylproline | 0.75 | 1.56 | 0.48 |
| cytidine 5'-monophosphate (5'-CMP) | 0.85 | 1.76 | 0.48 |
| hexadecadienoate (16:2n6) | 0.72 | 1.48 | 0.48 |
| hexadecenedioate (C16:1-DC)* | 0.47 | 0.95 | 0.50 |
| 1-nervonoyl-GPC (24:1n9)* | 0.57 | 1.10 | 0.52 |
| o-tyrosine | 0.62 | 1.20 | 0.52 |
| linoleate (18:2n6) | 0.71 | 1.36 | 0.52 |
| isoleucine | 0.73 | 1.39 | 0.53 |

| | | | |
|---|---|---|---|
| tyrosine | 0.64 | 1.22 | 0.53 |
| valine | 0.70 | 1.31 | 0.53 |
| 4-methylthio-2-oxobutanoate | 0.58 | 1.08 | 0.54 |
| 2-oleoylglycerol (18:1) | 0.76 | 1.41 | 0.54 |
| tryptophan | 0.66 | 1.22 | 0.54 |
| leucine | 0.70 | 1.27 | 0.55 |
| 1-oleoylglycerol (18:1) | 0.77 | 1.36 | 0.56 |
| phenylalanine | 0.69 | 1.21 | 0.57 |
| hydantoin-5-propionate | 0.59 | 1.02 | 0.58 |
| 13-HODE + 9-HODE | 0.70 | 1.20 | 0.58 |
| N6-acetyllysine | 0.98 | 1.66 | 0.59 |
| pyroglutamylalanine* | 0.42 | 0.71 | 0.59 |
| alanine | 0.84 | 1.37 | 0.62 |
| 2R,3R-dihydroxybutyrate | 0.84 | 1.36 | 0.62 |
| kynurenine | 0.85 | 1.28 | 0.66 |
| proline | 0.86 | 1.28 | 0.67 |
| (3'-5')-uridylylguanosine | 0.62 | 0.86 | 0.73 |
| alpha-tocopherol | 1.08 | 0.77 | 1.40 |
| pyridoxal | 1.07 | 0.74 | 1.44 |
| nicotinate | 1.13 | 0.77 | 1.47 |
| fucosterol | 1.21 | 0.73 | 1.64 |
| beta-sitosterol | 1.22 | 0.73 | 1.68 |
| 1-(1-enyl-palmitoyl)-2-oleoyl-GPE (P-16:0/18:1)* | 1.16 | 0.66 | 1.76 |
| diacetylchitobiose | 1.11 | 0.62 | 1.80 |
| stigmasterol | 1.07 | 0.59 | 1.81 |
| 1-palmitoyl-2-oleoyl-GPE (16:0/18:1) | 1.19 | 0.65 | 1.82 |
| 1-palmitoyl-2-linoleoyl-GPE (16:0/18:2) | 1.28 | 0.70 | 1.82 |
| carotene diol (3) | 1.11 | 0.61 | 1.82 |
| flavin mononucleotide (FMN) | 1.33 | 0.69 | 1.91 |
| flavin adenine dinucleotide (FAD) | 0.92 | 0.48 | 1.92 |
| 2-aminoadipate | 0.60 | 0.31 | 1.92 |
| indolepropionate | 1.02 | 0.53 | 1.92 |
| ergosterol | 0.70 | 0.35 | 1.98 |
| 3,4-dihydroxybenzoate | 0.88 | 0.44 | 2.02 |
| N,N,N-trimethyl-5-aminovalerate | 1.53 | 0.75 | 2.02 |
| 1-palmitoyl-2-linoleoyl-galactosylglycerol (16:0/18:2)* | 1.36 | 0.67 | 2.04 |
| beta-cryptoxanthin | 1.21 | 0.58 | 2.09 |
| 1-oleoyl-2-linoleoyl-GPC (18:1/18:2)* | 1.13 | 0.53 | 2.13 |
| 2-palmitoylglycerol (16:0) | 1.34 | 0.62 | 2.16 |
| pantethine | 1.05 | 0.47 | 2.25 |
| stearoyl ethanolamide | 1.65 | 0.72 | 2.30 |
| 1-stearoyl-2-dihomo-linolenoyl-GPC (18:0/20:3n3 or 6)* | 1.39 | 0.60 | 2.31 |
| 1-(1-enyl-stearoyl)-2-arachidonoyl-GPE (P-18:0/20:4)* | 1.14 | 0.49 | 2.36 |
| 2-methylcitrate/homocitrate | 1.48 | 0.62 | 2.38 |
| 1-palmitoyl-2-dihomo-linolenoyl-GPC (16:0/20:3n3 or 6)* | 1.51 | 0.63 | 2.40 |
| 1-stearoyl-2-linoleoyl-GPC (18:0/18:2)* | 1.70 | 0.70 | 2.43 |
| ergothioneine | 1.24 | 0.51 | 2.45 |
| 1-palmitoyl-2-arachidonoyl-GPE (16:0/20:4)* | 1.39 | 0.57 | 2.46 |
| gamma-tocopherol/beta-tocopherol | 1.18 | 0.47 | 2.52 |
| oxalate (ethanedioate) | 1.42 | 0.56 | 2.53 |
| 1,2-dilinoleoyl-GPE (18:2/18:2)* | 1.19 | 0.47 | 2.55 |
| 1-margaroyl-2-linoleoyl-GPC (17:0/18:2)* | 0.99 | 0.39 | 2.56 |
| 1-pentadecanoyl-2-linoleoyl-GPC (15:0/18:2)* | 1.27 | 0.49 | 2.60 |
| 1-stearoyl-2-arachidonoyl-GPC (18:0/20:4) | 1.64 | 0.61 | 2.70 |
| 1-(1-enyl-stearoyl)-2-oleoyl-GPE (P-18:0/18:1) | 1.13 | 0.41 | 2.73 |
| 1-stearoyl-2-dihomo-linolenoyl-GPE (18:0/20:3n3 or 6)* | 0.81 | 0.29 | 2.74 |
| pheophytin A | 1.15 | 0.42 | 2.75 |
| 1-stearoyl-2-arachidonoyl-GPI (18:0/20:4) | 0.99 | 0.36 | 2.77 |
| 1-palmitoyl-2-oleoyl-GPG (16:0/18:1) | 1.31 | 0.47 | 2.78 |

| | | | |
|---|---|---|---|
| dehydrophytosphingosine* | 1.99 | 0.70 | 2.83 |
| 1-oleoyl-2-linoleoyl-GPE (18:1/18:2)* | 1.04 | 0.37 | 2.83 |
| palmitoyl-oleoyl-glycerol (16:0/18:1) [2]* | 1.46 | 0.51 | 2.86 |
| 1-stearoyl-2-oleoyl-GPE (18:0/18:1) | 1.48 | 0.51 | 2.92 |
| 1-stearoyl-2-arachidonoyl-GPE (18:0/20:4) | 1.44 | 0.45 | 3.20 |
| 1-stearoyl-2-linoleoyl-GPE (18:0/18:2)* | 1.51 | 0.42 | 3.63 |
| valerate (5:0) | 1.34 | 0.35 | 3.83 |
| maltol | 1.39 | 0.35 | 3.97 |
| butyrate/isobutyrate (4:0) | 1.62 | 0.36 | 4.47 |
| 1-stearoyl-2-oleoyl-GPS (18:0/18:1) | 1.39 | 0.28 | 4.94 |
| nicotinate ribonucleoside | 1.97 | 0.36 | 5.46 |
| 2,8-quinolinediol | 1.23 | 0.13 | 9.36 |
| 3,5-dihydroxybenzoic acid | 7.18 | 0.43 | 16.84 |
